# The 3-phosphoinositide-dependent protein kinase 1 is an essential upstream activator of protein kinase A in malaria parasites

**DOI:** 10.1101/2021.05.27.445967

**Authors:** Eva Hitz, Natalie Wiedemar, Armin Passecker, Nicolas M. B. Brancucci, Ioannis Vakonakis, Pascal Mäser, Till S. Voss

## Abstract

Cyclic AMP (cAMP) signalling is crucial for the propagation of asexual malaria blood stage parasites. Recent work on *Plasmodium falciparum* demonstrated that phosphorylation of the invasion ligand AMA1 by the catalytic subunit of cAMP-dependent protein kinase A (PfPKAc) is an essential step during parasite invasion into red blood cells. However, the exact mechanisms regulating PfPKAc activity are only partially understood and PfPKAc function has not been extensively studied in gametocytes, the sexual blood stage forms that are essential for malaria transmission. By studying a conditional PfPKAc knockdown mutant, we confirm the essential role for PfPKAc in erythrocyte invasion and demonstrate that PfPKAc is involved in regulating gametocyte deformability. Interestingly, we observed that the conditional overexpression of PfPKAc also caused a profound lethal phenotype by preventing intra-erythrocytic parasite multiplication. Whole genome sequencing of parasites selected to tolerate increased PfPKAc expression levels identified missense mutations exclusively in the gene encoding the putative parasite orthologue of 3-phosphoinositide-dependent protein kinase-1 (PfPDK1). Using targeted mutagenesis, we show that PfPDK1 is essential for PfPKAc activation, most likely by phosphorylating T189 in the PfPKAc activation loop. In summary, our results corroborate the importance of tight regulation of PfPKA signalling for parasite survival and identify PfPDK1 as a crucial upstream regulator in this pathway and potential new drug target.

## Introduction

Malaria is caused by protozoan parasites of the genus *Plasmodium*. Infections with *P. falciparum* are responsible for the vast majority of severe and fatal malaria cases. People get infected through female *Anopheles* mosquitoes that inject sporozoites into the skin tissue during their blood meal. After reaching the liver, sporozoites infect and multiply inside hepatocytes, generating thousands of merozoites that are released into the blood stream. Merozoites invade red blood cells (RBCs) and develop intracellularly through the ring stage into a trophozoite and finally a schizont stage parasite, which undergoes four to five rounds of nuclear division followed by cytokinesis to produce up to 32 new merozoites. Upon rupture of the infected RBC (iRBC), the released merozoites infect new erythrocytes to initiate another intra-erythrocytic developmental cycle (IDC). The consecutive rounds of RBC invasion and intraerythrocytic parasite proliferation are responsible for all malaria-related pathology and deaths. Importantly, during each round of replication, a small subset of trophozoites commits to sexual development and their ring stage progeny differentiates over the course of 10-12 days and five distinct morphological stages (stage I-V) into mature male or female gametocytes. Sexual commitment occurs in response to environmental triggers that activate expression of the transcription factor PfAP2-G, the master regulator of sexual conversion ^1-3^. When taken up by an *Anopheles* mosquito, mature stage V gametocytes develop into gametes and undergo fertilization. The resulting zygote develops into an ookinete that migrates through the midgut wall and transforms into an oocyst, generating thousands of infectious sporozoites ready to be injected into another human host.

Erythrocyte invasion by merozoites is a highly regulated multi-step process starting with the initial attachment of the merozoite to the RBC surface, followed by parasite reorientation and formation of a so-called tight junction ^4^. The tight junction is the intimate contact area between the merozoite and RBC membranes that moves along the merozoite surface during the actin-myosin motor-driven invasion process ^4^. Alongside the secreted rhoptry neck proteins, the micronemal transmembrane protein apical membrane antigen 1 (AMA1) is an integral component of the tight junction ^4^. The cytoplasmic domain of AMA1 bears an essential role in merozoites during RBC invasion ^5-7^. In particular, the phosphorylation of residues in the AMA1 cytoplasmic tail (S610 and T613) is essential for AMA1 function in RBC invasion ^5-7^. Recent research has shown that the *P. falciparum* cyclic adenosine monophosphate (cAMP)-dependent protein kinase A (PfPKA) is responsible for AMA1 phosphorylation at S610 and hence essential for successful erythrocyte invasion ^5,6,8,9^.

PKA was discovered in the 1970s and is one of the best characterized eukaryotic protein kinases ^10^. In its inactive state, the PKA holoenzyme is a tetramer consisting of two regulatory subunits (PKAr) and two catalytic subunits (PKAc) ^11^. Upon binding of cAMP to PKAr, the PKAc subunits are released. PKAc release thus depends on cAMP levels, which are regulated by adenylyl cyclases (ACs) and phosphodiesterases (PDEs) that synthesise and hydrolyse cAMP, respectively ^11^. Furthermore, phosphorylation of PKAc is essential for its activity. *In vitro*, PKAc was shown to be active upon release from PKAr due to auto-phosphorylation ^12,13^. However, research in budding yeast and human cells demonstrated that the 3-phosphoinositide-dependent protein kinase-1 (PDK1) phosphorylates and activates PKAc *in vivo* ^14-18^. PDK1 has originally been identified as the kinase responsible for activating PKB/Akt in response to growth factor-induced phosphoinositide-3-kinase (PI3K) signalling in human cells ^17,19-21^. Subsequent studies have shown that PDK1 also activates a large number of other AGC-type kinases including PKA, PKG and PKC ^17,21^. AGC kinases dock with PDK1 via their so called PDK1-interacting fragment (PIF), a hydrophobic motif that binds to the PIF-binding pocket in the N-terminal region of the PDK1 kinase domain, and this interaction allows PDK1 to activate its substrates by activation loop phosphorylation ^17,21,22^. In case of human PKAc, the PDK1-dependent phosphorylation of T197 in the activation loop plays a crucial role in controlling PKAc structure, activity and function ^15,18,23,24^

In *P. falciparum*, PfPKA kinase consists of only one catalytic (PfPKAc) and one regulatory (PfPKAr) subunit ^25,26^ and cAMP levels in blood stage parasites are regulated by PfPDEβ ^27^, which hydrolyses both cAMP and cGMP, and PfACβ that synthesises cAMP ^8^. Analysis of conditional loss-of-function mutants showed that PfPKAc is essential for the process of merozoite invasion, where it is required for the phosphorylation and timely shedding of the invasion ligand AMA1 from the merozoite surface ^5,6,8,9^. Likewise, depletion of cAMP levels through conditional disruption of *pfacβ* phenocopied the invasion defect observed for the *pfpka*c mutant ^8^. Interestingly, a conditional *pfpdeβ* null mutant, which displays increased cAMP levels and PfPKAc hyperactivation, also showed a severe merozoite invasion defect that was linked to elevated phosphorylation and premature shedding of AMA1 ^27^. These studies highlighted that tight regulation of PfPKAc activity is crucial for successful merozoite invasion and parasite proliferation. In addition, PfPKA seems to have additional functions in blood stage parasites. Global phosphoproteomic studies of *pfpdeβ, pfacβ* and *pfpkac* conditional knock out (KO) cell lines identified 39 proteins as high confidence targets of cAMP/PfPKA-dependent phosphorylation ^8,27^. These proteins include invasion factors (e.g. AMA1 and coronin) and several proteins with predicted roles in other processes (e.g. chromatin organization and protein transport) or with unknown functions ^8,27^. In addition, cAMP/PfPKA-dependent signalling has been implicated in the regulation of ion channel conductance and new permeability pathways (NPP) in asexual blood stage parasites ^28^ and gametocytes ^29^ as shown through the use of pharmacological approaches (PKA/PDE inhibitors, exogenous 8-Bromo-cAMP) and transgenic cell lines (deletion of PfPDEδ, overexpression of PfPKAr) ^28,29^. Similar experiments identified a putative role for cAMP/PfPKA-dependent signalling in regulating gametocyte-infected erythrocyte deformability ^30^.

While several studies demonstrated the importance of cAMP in activating PfPKAc, the role of PfPKAc phosphorylation in regulating PfPKAc activity remains elusive. High throughput phosphoproteomic approaches identified several phosphorylated residues in PfPKAc, including T189 that corresponds to the PDK1 target residue T197 in the activation segment of mammalian PKAc ^31-34^. However, if and to what extent phosphorylation of T189 is important for PfPKAc activation in *P. falciparum* and whether T189 phosphorylation is deposited via auto-phosphorylation or by another kinase, is unknown. Furthermore, besides the well-established role for PfPKAc in parasite invasion, other possible functions of PfPKA in asexual and sexual development are only poorly understood.

Here, we used reverse genetics approaches to study the function of PfPKAc in asexual blood stage parasites, sexual commitment and gametocytogenesis. Our results confirm the essential role for PfPKAc in merozoite invasion and show that while PfPKAc plays no obvious role in the control of sexual commitment or gametocyte maturation, it contributes to the regulation of gametocyte-infected erythrocyte deformability. We further demonstrate that overexpression of PfPKAc is lethal in asexual blood stage parasites. Intriguingly, whole genome sequencing (WGS) of parasites selected to tolerate PfPKAc overexpression identified mutations exclusively in the gene encoding the *P. falciparum* orthologue of phosphoinositide-dependent protein kinase-1 (PfPDK1). Using targeted mutagenesis, we show that the T189 residue is crucial for PfPKAc activity and that activation of PfPKAc is strictly dependent on PfPDK1-mediated phosphorylation.

## Results

### Generation of a conditional PfPKAc loss-of-function mutant

To study PfPKAc function, we generated a conditional PfPKAc knockdown (cKD) line using a selection-linked integration (SLI)-based gene editing approach ^35^. Successful engineering tags the *pfpkac* gene with the fluorescent marker gene *gfp* fused to the *fkbp* destabilization domain (*dd*) sequence ^36,37^, followed by a sequence encoding the 2A skip peptide ^38,39^, the blasticidin S deaminase (*bsd*) marker gene and finally the *glmS* ribozyme element ^40^ (Supplementary Fig. 1). The FKBP/DD system allows for protein destabilization in the absence of the small molecule ligand Shield-1 ^36,37^, whereas the *glmS* ribozyme in the 3’ untranslated region of the mRNA causes transcript degradation in the presence of glucosamine (GlcN) ^40^. To be able to easily quantify sexual commitment rates, we modified the *pfpkac* gene in NF54/AP2-G-mScarlet parasites (Brancucci et al., manuscript in preparation), which allows identifying sexually committed cells by visualising expression of fluorophore-tagged PfAP2-G ^3,41^. After drug selection of transgenic NF54/AP2-G-mScarlet/PKAc cKD parasites, correct editing of *pfpkac* and absence of the wild type (WT) locus was confirmed by PCR on genomic DNA (gDNA) (Supplementary Fig. 1). Live cell fluorescence imaging demonstrated that PfPKAc-GFPDD was expressed in the cytosol and nucleus in mid schizonts before it localized to the periphery of developing merozoites in late schizonts as also described elsewhere ^8,9^ (Supplementary Fig. 1). PfPKAc-GFPDD expression was efficiently depleted in parasites cultured under –Shield-1/+GlcN conditions compared to the matching control (+Shield-1/–GlcN) (Fig. 1a). To assess the growth phenotype of PfPKAc-GFPDD-depleted parasites, we split ring stage parasite cultures (0-6 hours post invasion, hpi), maintained them separately under PfPKAc-GFPDD-depleting (–Shield-1/+GlcN) and -stabilizing conditions (+Shield-1/–GlcN) and quantified parasitaemia over three generations using flow cytometry (Fig. 1b and Supplementary Fig. 2). As expected, PfPKAc-GFPDD-depleted parasites were unable to proliferate because the merozoites failed to invade new RBCs (Fig. 1c). Hence, by combining two inducible expression systems we engineered a PfPKAc cKD mutant that allows efficient depletion of PfPKAc-GFPDD expression and phenocopies the lethal invasion defect previously observed with conditional PfPKAc KO mutants ^8,9^.

**Fig 1.**
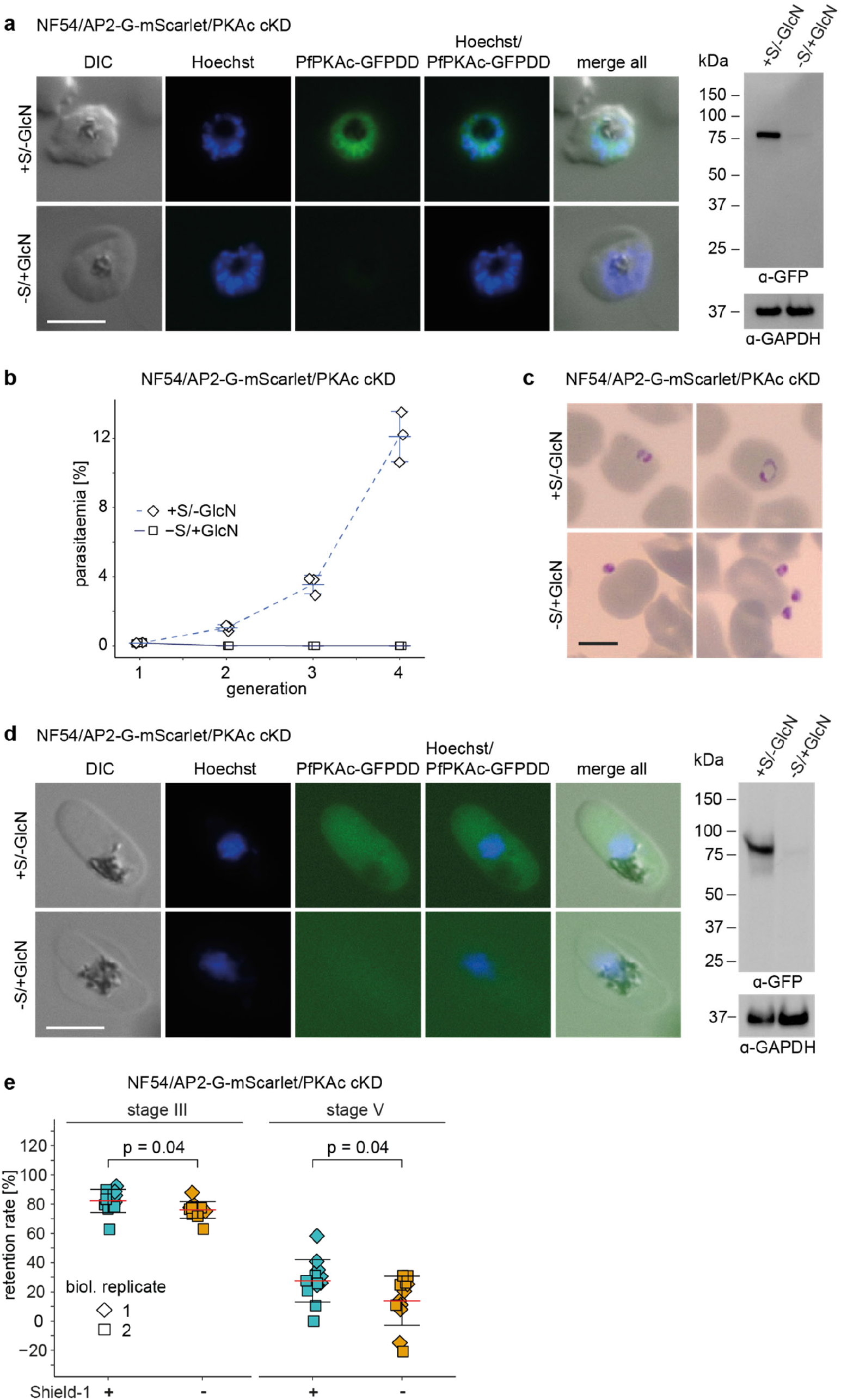
Depletion of PfPKAc in NF54/AP2-G-mScarlet/PKAc cKD parasites leads to a block in merozoite invasion and decreases gametocyte rigidity. **a**Expression of PfPKAc-GFPDD in late schizonts under protein- and RNA-depleting (–Shield-1/+GlcN) and control conditions (+Shield-1/– GlcN) as assessed by live cell fluorescence imaging and Western blot analysis. Synchronous parasites (0-8 hpi) were split (±Shield-1/±GlcN) 40 hours before sample collection. Representative fluorescence images are shown. Parasite DNA was stained with Hoechst. S, Shield-1; GlcN, glucosamine; DIC, differential interference contrast. Scale bar = 5 µm. For Western blot analysis, parasite lysates derived from equal numbers of parasites were loaded per lane. MW PfPKAc-GFPDD = 79.8 kDa, MW PfGAPDH = 36.6 kDa. **b** Increase in parasitaemia over three generations under PfPKAc-GFPDD-depleting (–Shield-1/+GlcN) and control conditions (+Shield-1/–GlcN). Open squares represent data points for individual replicates and the means and SD (error bars) of three biological replicates are shown. **c** Representative images from Giemsa-stained thin blood smears showing the progeny of parasites cultured under PfPKAc-GFPDD-depleting (–Shield-1/+GlcN) or control conditions (+Shield-1/–GlcN) conditions at 0-6 hpi (generation 2). Scale bar = 5 µm. **d** Expression of PfPKAc-GFPDD in stage V gametocytes (day 11) under protein- and RNA-depleting (–Shield-1/+GlcN) and control conditions (+Shield-1/–GlcN) as assessed by live cell fluorescence imaging and Western blot analysis. Representative fluorescence images are shown. Parasite DNA was stained with Hoechst. DIC, differential interference contrast. Scale bar = 5 µm. For Western blot analysis, parasite lysates derived from equal numbers of parasites were loaded per lane. MW PfPKAc-GFPDD = 79.8 kDa, MW PfGAPDH = 36.6 kDa. **e** Retention rates of stage III (day 6) and stage V (day 11) gametocytes cultured under PfPKAc-GFPDD-depleting (–Shield-1) (orange) and control conditions (+Shield-1) (blue). Coloured squares represent data points for individual replicates and the means and SD (error bars) of two biological replicates performed in six technical replicates each are shown. Differences in retention rates have been compared using an unpaired two-tailed Student’s t test (statistical significance cut-off: *p*<0.05).

**Fig 2.**
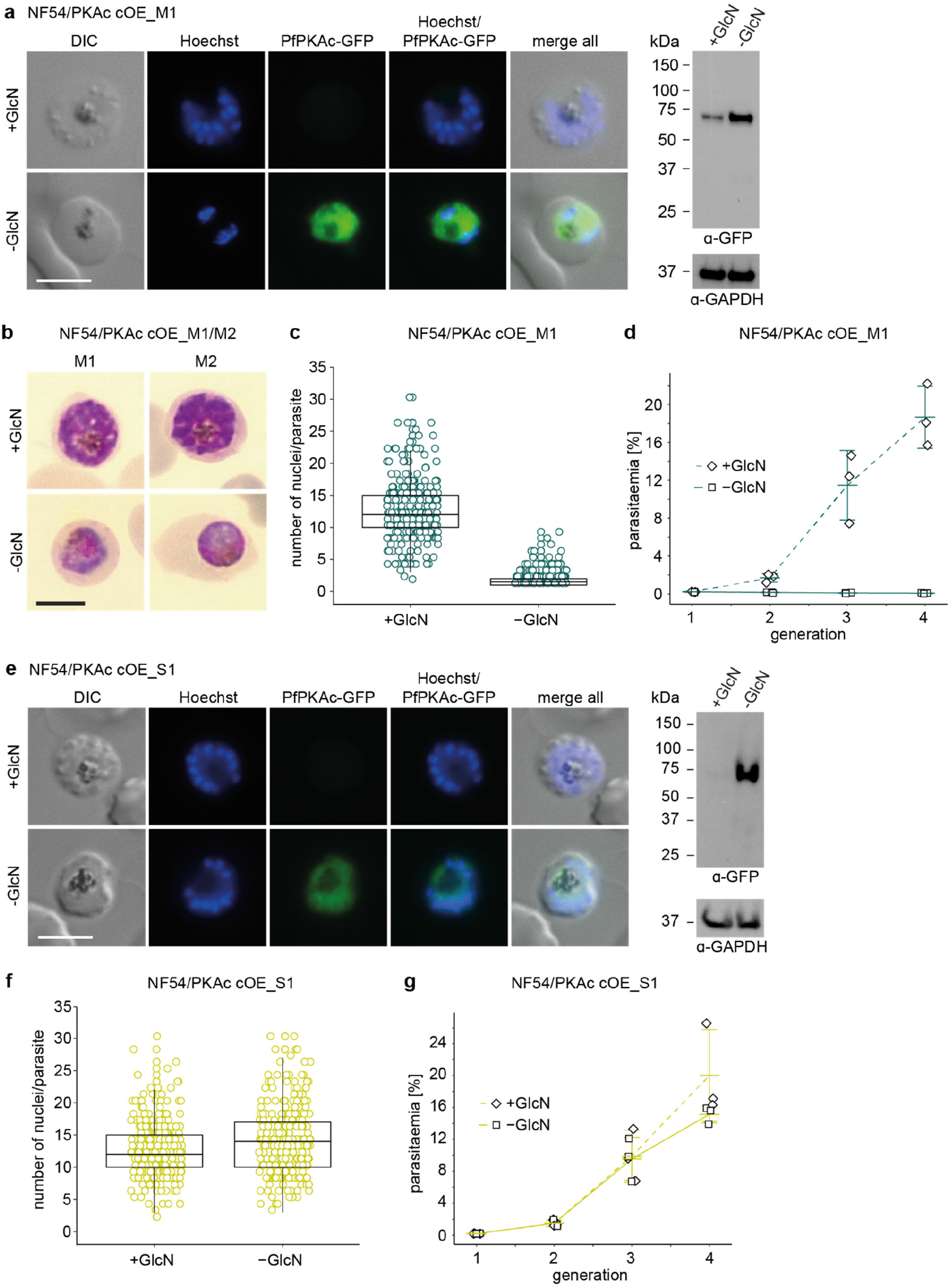
Overexpression of PfPKAc in NF54/PKAc cOE M1 parasites is lethal but survivor populations tolerant to PfPKAc OE can be selected. **a**Expression of PfPKAc-GFP in NF54/PKAc cOE M1 parasites under OE-inducing (–GlcN) and control conditions (+GlcN) as assessed by live cell fluorescence imaging and Western blot analysis. Synchronous parasites (0-8 hpi) were split (±GlcN) 40 hours before sample collection. Representative fluorescence images are shown. Parasite DNA was stained with Hoechst. DIC, differential interference contrast. Scale bar = 5 µm. GlcN, glucosamine. For Western blot analysis, lysates derived from equal numbers of parasites were loaded per lane. MW PfPKAc-GFP = 67.3 kDa, MW PfGAPDH = 36.6 kDa. **b** Representative images from Giemsa-stained thin blood smears showing NF54/PKAc cOE M1 and M2 parasites under PfPKAc-GFP OE-inducing (– GlcN) and control conditions (+GlcN). Synchronous parasites (0-8 hpi) were split (±GlcN) 40 hours before the images were captured. Scale bar = 5 µm. **c** Number of nuclei per schizont in NF54/PKAc cOE M1 parasites under PfPKAc-GFP OE-inducing (–GlcN) and control conditions (+GlcN). Each open circle represents one parasite. Data from three biological replicate experiments are shown and 100 parasites were counted in each experiment. The boxplots show data distribution (median, upper and lower quartile and whiskers). **d** Increase in parasitaemia in NF54/PKAc cOE M1 parasites over three generations under PfPKAc-GFP OE-inducing (–GlcN) and control (+GlcN) conditions. Synchronous parasites (0-6 hpi) were split (±GlcN) 18 hours before the first measurement in generation 1. Open squares represent data points for individual replicates and the means and SD (error bars) of three biological replicates are shown. **e** Expression of PfPKAc-GFP in NF54/PKAc cOE S1 survivor parasites under OE-inducing (–GlcN) and control conditions (+GlcN). Parasites were cultured and samples prepared as described in panel a. DIC, differential interference contrast. Scale bar = 5 µm. MW PfPKAc-GFP = 67.3 kDa, MW PfGAPDH = 36.6 kDa. **f** Number of nuclei per schizont in NF54/PKAc cOE S1 survivor parasites under PfPKAc-GFP OE-inducing (–GlcN) and control conditions (+GlcN). Each open circle represents one parasite. Data from three biological replicate experiments are shown and 100 parasites were counted in each experiment. The boxplots show data distribution (median, upper and lower quartile and whiskers). **g** Increase in parasitaemia in NF54/PKAc cOE S1 survivor parasites over three generations under PfPKAc-GFP OE-inducing (–GlcN) and control (+GlcN) conditions. Parasites were cultured as described in panel d. Open squares represent data points for individual replicates and the means and SD (error bars) of three biological replicates are shown.

### PfPKAc plays no major role in sexual commitment and gametocyte maturation but contributes to the regulation of gametocyte rigidity

To investigate whether PfPKAc activity regulates sexual commitment, we split ring stage parasites (0-6 hpi) and cultured them separately under PfPKAc-GFPDD-depleting (–Shield-1/+GlcN) and - stabilizing conditions (+Shield-1/–GlcN). Sexually committed parasites were identified based on PfAP2-G-mScarlet positivity in late schizonts (40-46 hpi) using high content imaging. We observed a significant increase in sexual commitment rates (SCRs) in PfPKAc-GFPDD-depleted (–Shield-1/+GlcN) compared to control parasites (+Shield-1/–GlcN) (1.56-fold ±0.13 SD) (Supplementary Fig. 3). We can exclude that these differences are caused by the Shield-1 compound as we have previously shown that Shield-1 treatment has no effect on SCRs ^42^. However, treatment with GlcN also caused a similar increase in SCRs in NF54/AP2-G-mScarlet control parasites (1.44-fold ±0.14 SD) (Supplementary Fig. 3). We therefore conclude that PfPKAc plays no major role in regulating sexual commitment, and that the increased SCRs observed in PfPKAc-GFPDD-depleted parasites are caused by the presence of 2.5 mM GlcN in the culture medium.

**Fig 3.**
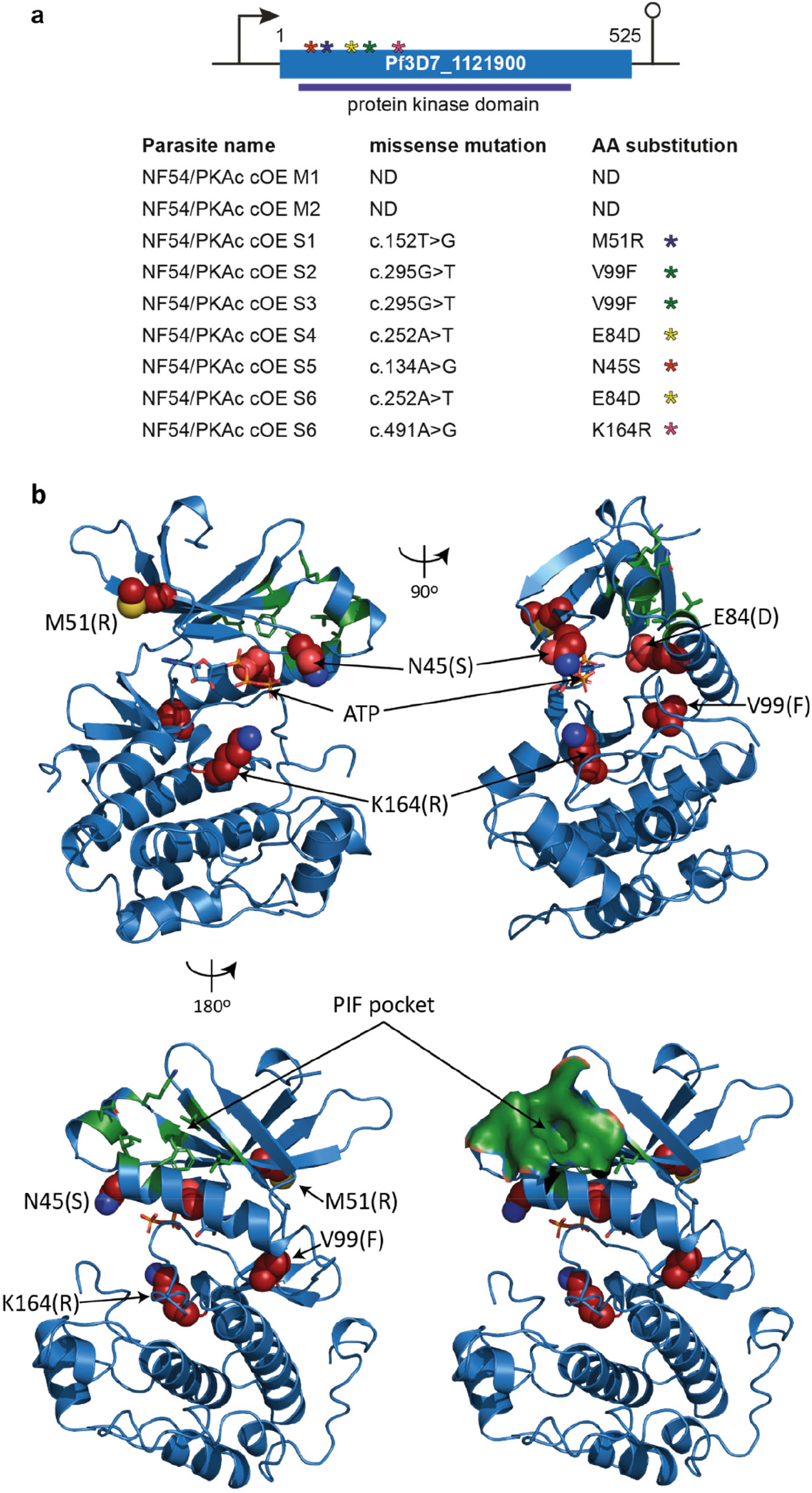
Whole genome sequencing reveals mutations in the Pf3D7_1121900/*pfpdk1* gene in six independently grown NF54/PKAc cOE survivor parasites. **a**Top: Schematic of the Pf3D7_1121900/*pfpdk1* gene. Asterisks indicate the approximate localization of the five missense mutations identified in the six different NF54/PKAc cOE survivors. Bottom: Summary of the sequence information obtained from WGS of gDNA of the two NF54/PKAc cOE clones (M1, M2) and the six independently grown survivors (S1-S6). Missense mutations and their positions within the *pfpdk1* coding sequence as well as the corresponding amino acid substitutions in the PfPDK1 protein sequence are shown. c., cDNA. **b** Predicted PfPDK1 structure shown in orthogonal (top left vs top right) or opposing (top left vs lower left) views. PfPDK1 was modelled on the crystallographic structure of human PDK1 (PDB ID 1UU9) ^51^. PfPDK1 segments with no correspondence in human PDK1 (amino acids 1-27, 188-307 and 423-525) were omitted from modelling. The ATP substrate (sticks), mutated amino acids (substitution in parenthesis; red spheres) and residues forming the PIF-binding pocket ^22^ (green sticks) are indicated. The PIF-binding pocket is shown in surface representation in the lower right view.

Next, we tested if PfPKAc-GFPDD depletion affects gametocyte morphology or male gametocyte exflagellation. To conduct these experiments, we induced sexual commitment by culturing parasites in serum-free minimal fatty acid medium (–SerM) ^3^. After reinvasion, parasites were split and cultured separately under PfPKAc-GFPDD-depleting (–Shield-1/+GlcN) or -stabilizing conditions (+Shield-1/– GlcN). For the first six days of gametocyte maturation, the growth medium was supplemented with 50 mM N-acetyl-D-glucosamine (GlcNac) to eliminate asexual parasites ^43^. Despite efficient depletion of PfPKAc-GFPDD expression (Fig. 1d and Supplementary Fig. 4), we could not detect morphological abnormalities in PfPKAc-GFPDD-depleted gametocytes in any of the five developmental stages (I-V) based on visual inspection of Giemsa-stained thin blood smears (Supplementary Fig. 4). While the exflagellation rates (ExRs) of PfPKAc-GFPDD-depleted male stage V gametocytes (–Shield-1/+GlcN) were significantly reduced by more than 50% compared to the control (+Shield-1/–GlcN), this effect was again caused by the presence of GlcN in the culture medium (Supplementary Fig. 4). When PfPKAc-GFPDD expression was depleted by Shield-1 removal only (–Shield-1/–GlcN), no significant differences in ExRs were observed even though PfPKAc-GFPDD expression was efficiently reduced (Supplementary Fig. 4). In line with this result, GlcN treatment also led to a significant reduction in ExRs of NF54 WT gametocytes (+GlcN) to less than 50% compared to the control (–GlcN) (Supplementary Fig. 4).

**Fig 4.**
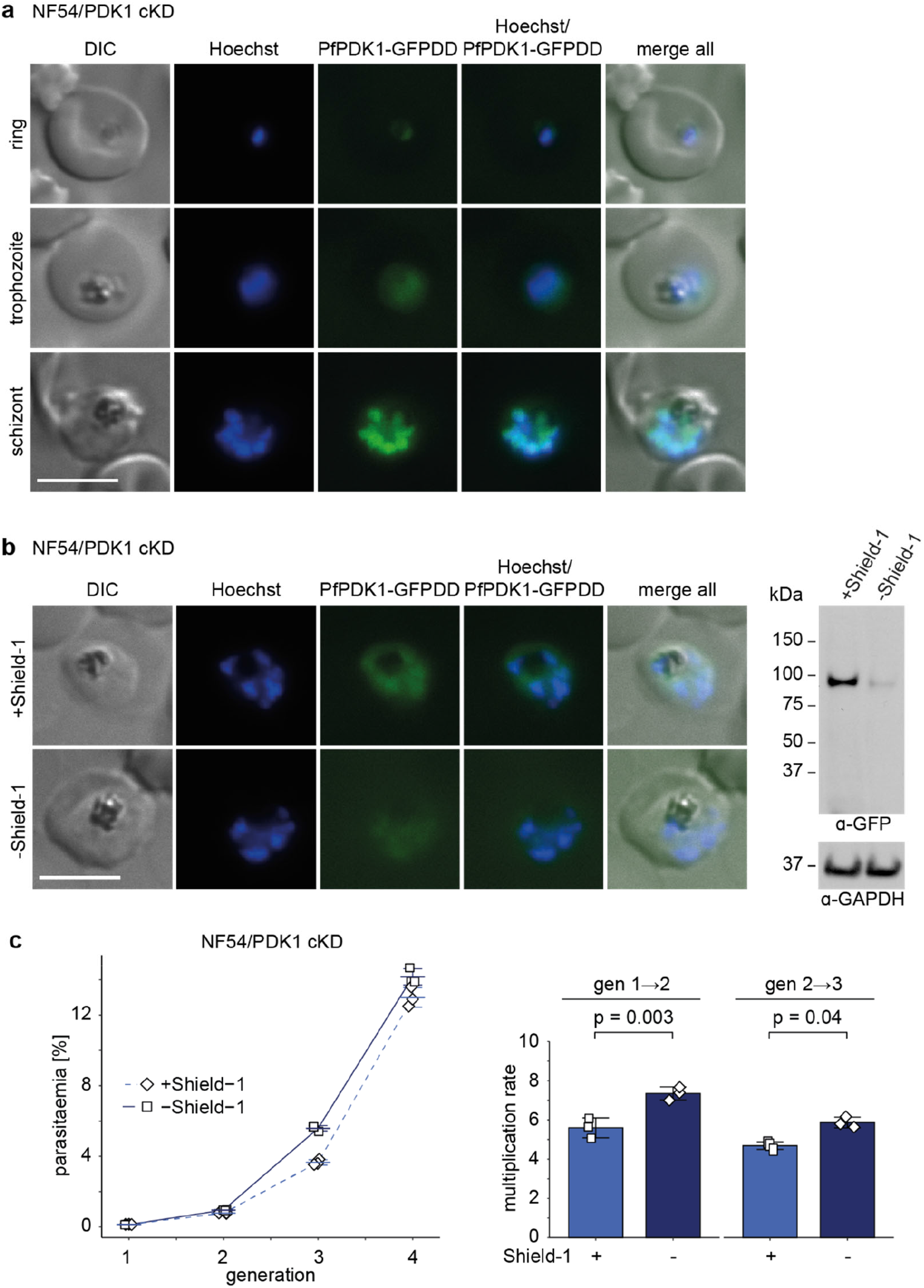
Knockdown of PfPDK1 expression in NF54/PDK1 cKD parasites has no major effect on asexual parasite growth. **a**Expression of PfPDK1-GFPDD in ring (18-24 hpi), trophozoite (24-30 hpi) and schizont (42-48 hpi) stages cultured under protein-stabilizing (+Shield-1) conditions by live cell fluorescence imaging. Representative fluorescence images are shown. Parasite DNA was stained with Hoechst. DIC, differential interference contrast. Scale bar = 5 µm. **b** Expression of PfPDK1-GFPDD under protein-depleting (–Shield-1) and control conditions (+Shield-1) by live cell fluorescence imaging and Western blot analysis. Synchronous parasites (0-8 hpi) were split (±Shield-1) 40 hours before collection of the samples. Representative fluorescence images are shown. Parasite DNA was stained with Hoechst. DIC, differential interference contrast. Scale bar = 5 µm. For Western blot analysis, lysates derived from an equal numbers of parasites were loaded per lane. MW PfPDK1-GFPDD = 101.1 kDa, MW PfGAPDH = 36.6 kDa. **c** Increase in parasitaemia (left) and parasite multiplication rates (right) under PfPDK1-GFPDD-depleting (–Shield-1) and control (+Shield-1) conditions. Synchronous parasites (0-6 hpi) were split (±Shield-1) 18 hours before the first measurement in generation 1. Open squares represent data points for individual replicates and the means and SD (error bars) of three biological replicates are shown. Differences in multiplication rates have been compared using a paired two-tailed Student’s t test (statistical significance cut-off: *p*<0.05).

Lastly, we tested if PfPKAc activity is involved in regulating gametocyte rigidity. Immature gametocytes display high cellular rigidity and sequester in the bone marrow and the spleen, whereas stage V gametocytes are more deformable and can re-enter the bloodstream to be taken up by feeding mosquitoes ^44-46^. Experiments employing PKA and PDE inhibitors, a transgenic cell line overexpressing PfPKAr, or treatment with exogenous 8-bromo-cAMP to increase cellular cAMP levels provided evidence for a potential role for PfPKAc in maintaining the rigidity of immature gametocyte-infected erythrocytes ^30^. To test if PfPKAc is indeed involved in controlling this process, we measured the deformability status of immature stage III and mature stage V gametocytes using microsphiltration experiments ^45^. Microsphiltration exploits the fact that differences in cellular rigidity correlate with cell retention rates in a microsphere-based artificial spleen system ^45,46^. We observed a significant decrease in the retention rates of PfPKAc-GFPDD-depleted gametocytes (–Shield-1) compared to the control (+Shield-1), both in immature stage III (day 6) (76.0% ±5.7 SD vs 82.2% ±8.0 SD) and mature stage V (day 11) gametocytes (13.9% ±16.8 SD vs 27.6% ±14.5 SD) (Fig. 1e). In contrast, NF54 WT gametocytes cultured in the presence or absence of Shield-1 showed no difference in retention rates (Supplementary Fig. 4).

In summary, our results demonstrate that PfPKAc plays no major role in the regulation of sexual commitment, gametocyte maturation or male gametogenesis but that gametocyte-infected RBC rigidity is at least partially regulated by PfPKAc. Furthermore, we discovered that the presence of 2.5 mM GlcN in the culture medium affects both sexual commitment and exflagellation rates, which needs to be taken into account when studying these processes in conditional mutants employing the *glmS* riboswitch system.

### PfPKAc overexpression in asexual blood stage parasites is lethal

The merozoite invasion and post-invasion developmental defects observed for *pfpdeβ* KO parasites had been linked to increased cAMP levels and PfPKAc hyperactivity ^27^. To further study the consequences of PfPKAc overexpression, we generated a PfPKAc conditional overexpression (cOE) line using CRIPSR/Cas9-based gene editing. To this end, we inserted an ectopic *pfpkac-gfp* transgene cassette into the non-essential *glp3* (*cg6*, Pf3D7_0709200) locus in NF54 WT parasites (NF54/PKAc cOE) (Supplementary Fig. 5). Here, the constitutive *calmodulin* (PF3D7_1434200) promoter and a *glmS* ribozyme element ^47^ control expression of the *pfpkac-gfp* gene. Since the initial transgenic NF54/PKAc cOE population still contained some parasites carrying the WT *glp3* locus, we obtained clonal lines by limiting dilution cloning ^48^. In two clones (NF54/PKAc cOE M1 and M2), correct integration of the inducible PfPKAc-GFP OE expression cassette and absence of WT parasites was confirmed by PCR on gDNA (Supplementary Fig. 5). Live cell fluorescence imaging and Western blot analysis confirmed the efficient induction of PfPKAc-GFP OE upon GlcN removal in both clones (Fig. 2a and Supplementary Fig. 6). Interestingly, PfPKAc-GFP OE (–GlcN) resulted in a complete block in parasite development half way through the IDC (Fig. 2b). To study this growth defect in more detail, we quantified the number of nuclei per parasite in late schizont stages (40-46 hpi), 40 hours after triggering PfPKAc-GFP OE (– GlcN) in young ring stage parasites (0-6 hpi). This experiment revealed that NF54/PKAc cOE M1 parasites overexpressing PfPKAc-GFP did not develop beyond the late trophozoite/early schizont stage as most parasites contained only one or two nuclei as opposed to the control population (+GlcN) that progressed normally through several rounds of nuclear division (Fig. 2c). To test whether these parasites were still able to produce progeny, we performed parasite multiplication assays. NF54/PKAc cOE M1 ring stage parasites were split at 0-6 hpi, cultured separately in the presence or absence of GlcN and parasite multiplication was quantified over three generations. PfPKAc-GFP OE (–GlcN) completely failed to multiply as no increase in parasitaemia was observed, in contrast to the control population (+GlcN) that multiplied normally (Fig. 2d and Supplementary Fig. 6). Notably, however, viable PfPKAc-GFP OE parasites emerged approximately two weeks after maintaining NF54/PKAc cOE M1 parasites constantly in culture medium lacking GlcN. We termed these parasites “PfPKAc OE survivors” (NF54/PKAc cOE S1). Importantly, NF54/PKAc cOE S1 parasites still overexpressed PfPKAc-GFP in absence of GlcN (Fig. 2e and Supplementary Fig. 6). Furthermore, Sanger sequencing confirmed that neither the endogenous nor the ectopic *pfpkac* genes in the NF54/PKAc cOE S1 survivor population carried any mutations. Quantifying the number of nuclei per schizont and parasite multiplication assays revealed that NF54/PKAc cOE S1 parasites were completely tolerant to PfPKAc-GFP OE and developed and multiplied identically irrespective of whether PfPKAc-GFP OE was induced (–GlcN) or not (+GlcN) (Figs. 2f, 2g and Supplementary Fig. 6).

**Fig 5.**
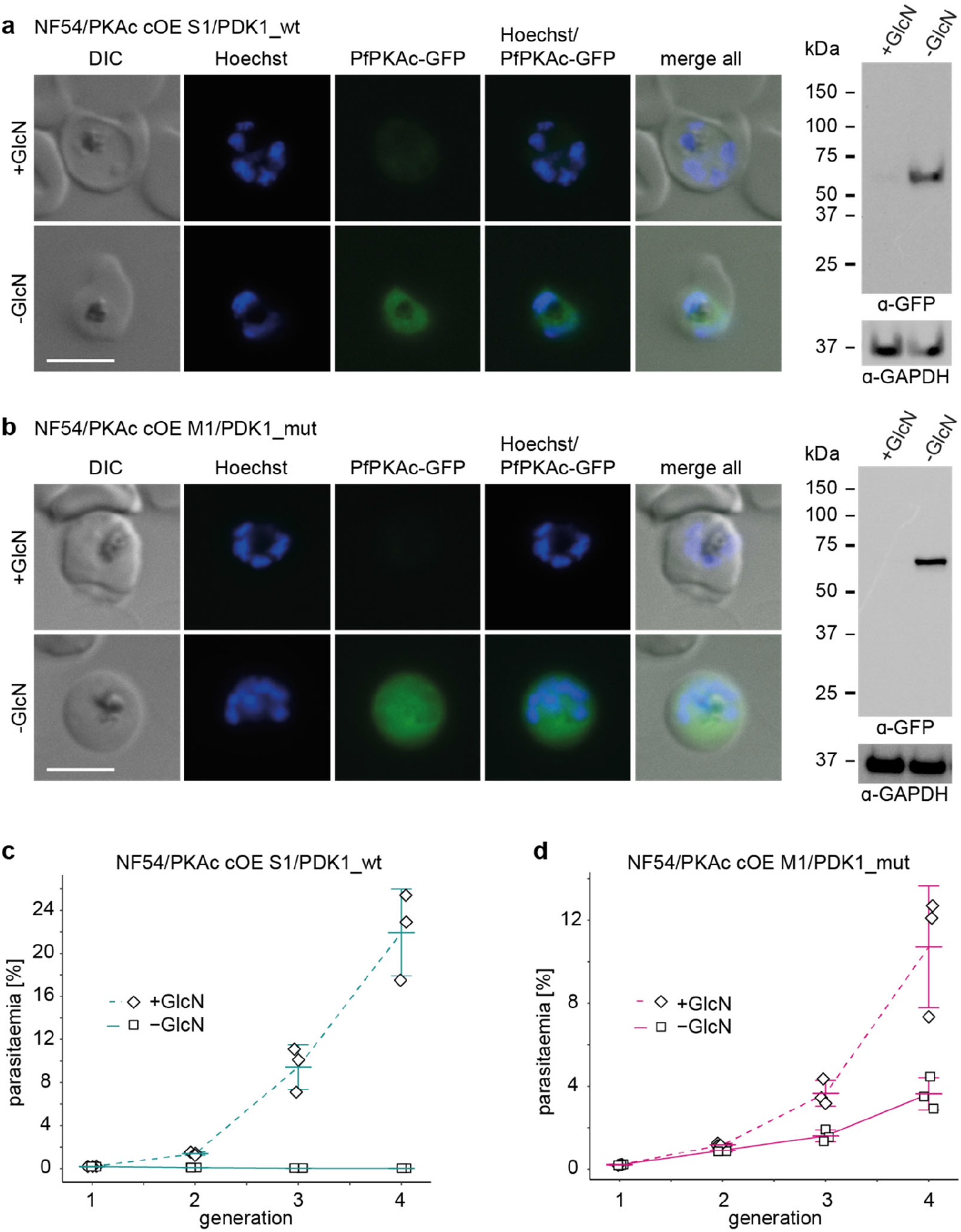
Targeted mutagenesis of PfPDK1 confirms its essential role in activating PfPKAc. **a**,**b** Expression of PfPKAc-GFP under OE-inducing (–GlcN) and control conditions (+GlcN) in NF54/PKAc cOE S1/PDK1_wt parasites (a) and NF54/PKAc cOE M1/PDK1_mut parasites (b) by live cell fluorescence imaging and Western blot analysis. Synchronous parasites (0-8 hpi) were split (±GlcN) 40 hours before collection of the samples. Representative fluorescence images are shown. Parasite DNA was stained with Hoechst. DIC, differential interference contrast. Scale bar = 5 µm. GlcN, glucosamine. For Western blot analysis, lysates derived from an equal number of parasites were loaded per lane. MW PfPKAc-GFP = 67.3 kDa, MW PfGAPDH = 36.6 kDa. **c**,**d** Increase in parasitaemia of NF54/PKAc cOE S1/PDK1_wt parasites (c) and NF54/PKAc cOE M1/PDK1_mut parasites (d) over three generations under PfPKAc-GFP OE-inducing (–GlcN) and control conditions (+GlcN). Synchronous parasites (0-6 hpi) were split (±GlcN) 18 hours before the first measurement in generation 1. Open squares represent data points for individual replicates and the means and SD (error bars) of three biological replicates are shown.

**Fig 6.**
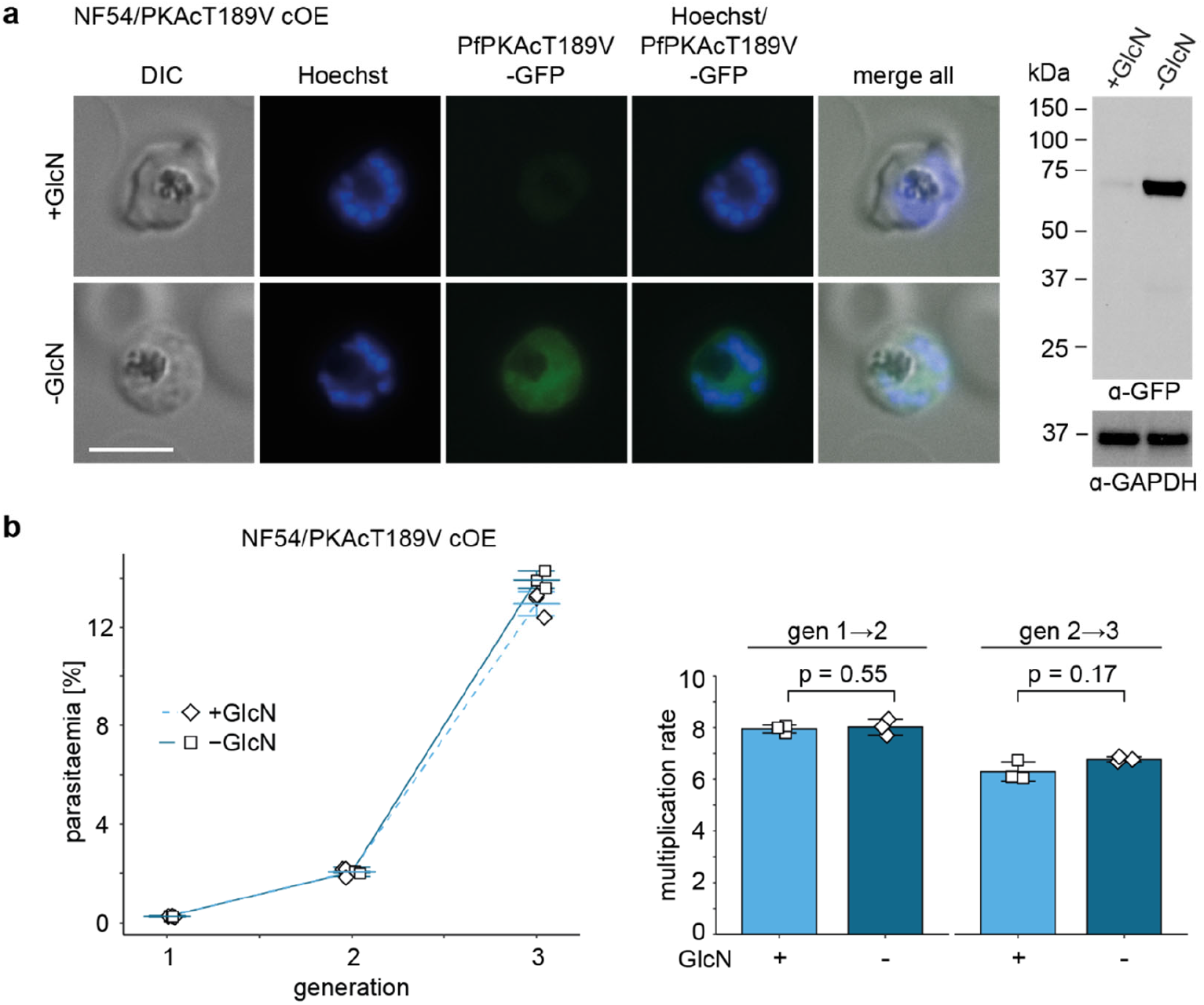
Overexpression of PfPKAcT189V has no effect on intra-erythrocytic parasite development and multiplication. **a**Expression of PfPKAcT189V-GFP under OE-inducing (–GlcN) and control conditions (+GlcN) by live cell fluorescence imaging and Western blot analysis. Synchronous parasites (0-8 hpi) were split (±GlcN) 40 hours before collection of the samples. Representative fluorescence images are shown. Parasite DNA was stained with Hoechst. DIC, differential interference contrast. Scale bar = 5 µm. GlcN, glucosamine. For Western blot analysis, lysates derived from equal numbers of parasites were loaded per lane. MW PfPKAc-GFP = 67.3 kDa, MW PfGAPDH = 36.6 kDa. **b** Increase in parasitaemia (left) and parasite multiplication rates (right) of NF54/PKAcT189V cOE parasites over three generations under PfPKAcT189V-GFP OE-inducing (–GlcN) and control conditions (+GlcN). Synchronous parasites (0-6 hpi) were split (±GlcN) 18 hours before the first measurement in generation 1. Open squares represent data points for individual replicates and the means and SD (error bars) of three biological replicates are shown. Differences in multiplication rates have been compared using a paired two-tailed Student’s t test (statistical significance cut-off: *p*<0.05).

In conclusion, PfPKAc OE causes a lethal phenotype in asexual parasites by preventing parasite development beyond the late trophozoite/early schizont stage. However, parasites tolerant to PfPKAc-GFP OE can be selected for and these parasites show no defect in intra-erythrocytic development and parasite multiplication.

### Parasites tolerant to PfPKAc overexpression carry mutations in the gene encoding *P. falciparum* 3-phosphoinositide-dependent protein kinase-1 (PfPDK1)

The above findings suggested that genetic mutations in the NF54/PKAc cOE S1 survivor population might cause their tolerance to elevated PfPKAc-GFP expression levels. To address this hypothesis, we performed whole genome sequencing (WGS) of the two unselected NF54/PKAc cOE clones M1 and M2 and the six independently grown survivor populations (NF54/PKAc cOE S1-S6), three each originating from the M1 and M2 clones, respectively. Intriguingly, we found that all six NF54/PKAc cOE survivors, but not the unselected M1 and M2 clones, carried missense mutations in the gene encoding a putative serine/threonine protein kinase (Pf3D7_1121900) (Fig. 3a and Supplementary Data 1). No other mutations were identified in the NF54/PKAc cOE survivor populations, consistent with the Sanger sequencing results, neither the endogenous nor the ectopic *pfpkac* gene carried mutations in any of the six survivors. The *P. vivax* orthologue of Pf3D7_1121900 (PVX_091715) is annotated as putative 3-phosphoinositide dependent protein kinase-1 (PDK1) (www.plasmodb.org), and a multiple sequence alignment suggested that Pf3D7_1121900 is indeed an orthologue of PDK1, a kinase widely conserved among eukaryotes and known as a master regulator of AGC kinases including PKA ^17,49^ (Supplementary Fig. 7). However, similar to PDK1 enzymes from most fungi, non-vascular plants and other alveolates, Pf3D7_1121900 and its *P. vivax* orthologue lack the C-terminal phospholipid-binding pleckstrin-homology (PH) domain that is found in PDK1s from animals and vascular plants and important to localise PDK1 to the plasma membrane for PKB activation in response to the PI3K-dependent production of phosphatidylinositol bis-/trisphosphates ^17,49,50^. Construction of a homology model of the Pf3D7_1121900 protein kinase domain based on the human PDK1 crystallographic structure (PDB ID 1UU9) ^51^ allowed us to visualize the location of the amino acids mutated in the six NF54/PKAc cOE survivors (Fig. 3b). None of these mutations affected the putative PIF-binding pocket. Rather, all mutated amino acids were located at or in the periphery of the predicted ATP-binding cleft, although only one mutation (N45S) may influence ATP coordination directly (Fig. 3b and Supplementary Fig. 7). We hence surmise that all identified mutations alter the catalytic activity but not the protein interaction preferences of PfPDK1.

In conclusion, we identified missense mutations in the Pf3D7_1121900 gene in six independently obtained NF54/PKAc cOE survivor parasite lines, which likely confer tolerance to PfPKAc OE. Bioinformatic analysis indicates that this gene encodes PfPDK1, the *P. falciparum* orthologue of PDK1, and modelling of the PfPDK1 structure predicts that the acquired mutations may affect its catalytic activity.

### Conditional depletion of PfPDK1 has no major effect on parasite multiplication, sexual commitment and gametocytogenesis

To gain further insight into the function of PfPDK1, we attempted to generate a *pfpdk1* KO line by gene disruption using a CRISPR/Cas9-based single plasmid approach ^52^. However, consistent with the results obtained in previous kinome- or genome-wide KO screens in *P. falciparum* and *P. berghei* ^31,53,54^, we failed to obtain a *pfpdk1* null mutant, indicating that the gene is essential in asexual parasites. We therefore engineered the PfPDK1 conditional knockdown line NF54/PDK1 cKD by tagging the endogenous *pfpdk1* gene with g*fp* fused to the *dd* sequence in NF54 WT parasites (Supplementary Fig. 8). Correct editing of the *pfpdk1* gene and absence of the wild type locus was confirmed by PCR on gDNA. Donor plasmid integration downstream of the *pfpdk1* gene was detected in a subset of parasites (Supplementary Fig. 8), but this is not expected to compromise *pfpdk1-gfpdd* expression since the 551 bp 3’ homology region (HR) used for homology-directed repair includes the native terminator as based on published RNA-seq data ^55^. Live cell fluorescence imaging and Western blot analysis revealed that PfPDK1-GFPDD is expressed in the cytosol and nucleus throughout the IDC with highest expression in schizonts, consistent with published gene expression data ^55,56^ (Fig. 4a and Supplementary Fig. 9). Furthermore, PfPDK1-GFPDD expression was efficiently reduced after Shield-1 removal compared to parasites grown in the presence of Shield-1 (Fig. 4b and Supplementary Fig. 9). Surprisingly, PfPDK1-GFPDD-depleted parasites (–Shield-1) showed slightly higher multiplication rates compared to the control (+Shield-1) (Fig. 4c). This increased multiplication rate cannot be attributed to the removal of Shield-1 itself, since NF54 WT parasites cultured in the presence or absence of Shield-1 multiplied equally (Supplementary Fig. 9). We also tested whether PfPDK1-GFPDD plays a role in gametocytogenesis but did not detect any differences in SCRs, gametocyte morphology or exflagellation rates when comparing PfPDK1-GFPDD-depleted (–Shield-1) with control parasites (+Shield-1) (Supplementary Fig. 10).

Taken together, we show that PfPDK1 is expressed in the parasite nucleus and cytosol throughout asexual development, reaching peak expression in schizonts. While PfPDK1 is likely essential, the results obtained with the PfPDK1 cKD mutant suggest that residual PfPDK1 expression levels are sufficient to sustain parasite viability. Furthermore, PfPDK1 seems to play no major role in regulating sexual commitment, gametocytogenesis or male gametogenesis, but it is again conceivable that residual PfPDK1 expression in the cKD mutant may have been sufficient to maintain these processes.

### Expression of WT PfPDK1 is incompatible with PfPKAc overexpression whereas expression of the M51R PfPDK1 mutant causes PfPKAc overexpression tolerance

To confirm the specific function of PfPDK1 in regulating PfPKAc activity, we used CRISPR/Cas9-based targeted mutagenesis to change the PfPDK1 M51R mutation back to the WT sequence in the NF54/PKAc cOE S1 survivor line (NF54/PKAc cOE S1/PDK1_wt). Sanger sequencing verified the successful reversion of the PfPDK1 M51R mutation (Supplementary Fig. 11), and live cell fluorescence imaging and Western Blot analysis confirmed that GlcN removal still triggered efficient PfPKAc-GFP OE in NF54/PKAc cOE S1/PDK1_wt parasites (Fig. 5a and Supplementary Fig. 11). Strikingly, PfPKAc-GFP OE (–GlcN) led to a complete block in parasite replication, showing that reverting the PfPDK1 M51R mutation completely restored the PfPKAc OE-sensitive phenotype (Fig. 5c and Supplementary Fig. 11). To complement these experiments, we also attempted to introduce the PfPDK1 M51R mutation into NF54 WT parasites and the NF54/PKAc cOE clone M1. Two independent attempts to mutate PfPDK1 in NF54 WT parasites failed, suggesting that fully functional PfPDK1 is strictly required for parasite viability under normal PfPKAc expression levels. In contrast, we readily succeeded in introducing the PfPDK1 M51R mutation into the NF54/PKAc cOE clone M1 (NF54/PKAc cOE M1/PDK1_mut) (Supplementary Fig. 11). Western Blot analysis and live cell fluorescence imaging confirmed that GlcN removal still triggered efficient PfPKAc-GFP OE (Fig. 5b and Supplementary Fig. 11). Parasite multiplication assays revealed that the PfPDK1 M51R point mutation rendered NF54/PKAc cOE M1/PDK1_mut parasites tolerant to PfPKAc-GFP OE (–GlcN) (Fig. 5d and Supplementary Fig. 11). However, in contrast to NF54/PKAc cOE S1 parasites, which carry the same M51R PfPDK1 mutation, the multiplication rates of NF54/PKAc cOE M1/PDK1_mut parasites overexpressing PfPKAc-GFP (–GlcN) reached only 60% to 80% compared to the matching control (+GlcN) (Fig. 5d and Supplementary Fig. 11). This observation suggested that an additional selection step had taken place in NF54/PKAc cOE S1 parasites that conferred full tolerance to PfPKAc-GFP OE (see Fig. 3g). Indeed, based on the WGS data the parental NF54/PKAc cOE M1 clone carries eight copies of the PfPKAc cOE transgene cassette, whereas the NF54/PKAc cOE S1 survivor population carries only two copies (Supplementary Fig. 12). We therefore believe that due to the negative impact of continuous PfPKAc-GFP OE on parasite viability, NF54/PfPKAc cOE S1 parasites carrying fewer PfPKAc cOE cassettes and hence lower overall PfPKAc expression levels had a comparative advantage during the selection process for PfPKAc-GFP OE tolerance.

These data collectively confirm the importance of the *pfpdk1* mutations identified in NF54/PKAc cOE survivor parasites in conferring resistance to PfPKAc OE and suggest that PfPDK1 is the kinase that phosphorylates and activates PfPKAc. They furthermore imply that the PfPDK1 M51R mutant kinase can still phosphorylate PfPKAc with reduced efficiency and therefore sustain parasite viability in the presence of elevated PfPKAc levels.

### The PfPKAc activation loop residue T189 is essential for PfPKAc activity

Previous research in human cell lines and yeast identified a specific threonine residue in the PKAc activation loop (T197 in mammals) as the target of PDK1-dependent phosphorylation ^15,18,23^. In PfPKAc, T189 likely represents the activation loop phosphorylation site corresponding to T197 in human PKAc. Hence, we tested whether the T189 residue is indeed important for PfPKAc activity. To achieve this, we employed the same approach as already used to obtain NF54/PKAc cOE parasites (Supplementary Fig. 5) to generate the NF54/PKAcT189V cOE line that conditionally overexpresses a mutated version of PfPKAc in which T189 has been substituted with a non-phosphorylatable valine residue ^57,58^ (Supplementary Fig. 13). Correct insertion of the PfPKAcT189V cOE cassette into the *glp3* locus was verified by PCR on gDNA (Supplementary Fig. 13). Live cell fluorescence microscopy and Western blot analysis confirmed that PfPKAcT189V-GFP OE was efficiently induced upon removal of GlcN (Fig. 6a and Supplementary Fig. 13). Strikingly, in contrast to the lethal effect provoked by OE of PfPKAc-GFP, OE of the PfPKAcT189V-GFP mutant (–GlcN) had no effect on intra-erythrocytic parasite development, multiplication and survival when compared to the control population (+GlcN) (Fig. 6b). Hence, these results suggest that PfPKAc activity is strictly dependent on phosphorylation of T189 in the activation loop segment. In combination with the findings obtained through the mutational analysis of PfPDK1, they further imply that PfPDK1 is directly or indirectly responsible for T189 phosphorylation and thus activation of PfPKAc.

### Kinase inhibitors targeting human PDK1 are active against asexual blood stage parasites

Many PDK1-dependent AGC kinases (e.g. AKT/PKB, RSK, PKC, S6K, SGK) are downstream effectors in the phosphoinositide 3-kinase (PI3K)/protein kinase B (AKT) or mitogen-activated protein kinase (MAPK) growth factor signalling pathways and are aberrantly activated in various types of cancer in humans ^59^. In addition, PDK1 expression itself is augmented in many tumors ^59^. For these reasons, human PDK1 is pursued as a potential drug target for cancer therapy and a large number of inhibitors targeting human PDK1 have been developed and patented over the past 15 years ^60-62^. For instance, BX-795 and BX-912, two related aminopyrimidine compounds, inhibit recombinant human PDK1 activity with half-maximal inhibitory concentrations (IC_50_) of 11 nM and 26 nM, respectively ^63^, and the aminopyrimidine-aminoindazole GSK2334470 has similar *in vitro* potency against PDK1 (15 nM IC_50_) ^64,65^. While BX-795 was shown to also inhibit several other human kinases *in vitro* ^66^, GSK2334470 displayed high specificity for PDK1 over a large panel of other recombinant human kinases ^64^. In cell-based assays, all three compounds inhibited the PDK1-dependent phosphorylation of several AGC kinase substrates at submicromolar concentrations ^63-65^. Here, we tested these three commercially available ATP-competitive inhibitors of human PDK1 for their potential to kill *P. falciparum* asexual blood stage parasites using a [^3^H]-hypoxanthine incorporation assay ^67^. We found that BX-795, BX-912 and GSK2334470 all inhibited parasite proliferation with IC_50_ values of 1.83 µM (± 0.23 SD), 1.31 µM (± 0.24 SD) and 1.83 µM (± 0.11 SD), respectively (Supplementary Fig. 14). Whether the lethal effect of these compounds is due to the specific inhibition of PfPDK1 or whether they target additional/other parasite kinases is unknown at this stage. However, based on the promising activities of these molecules against blood stage parasites and the expected essentiality of PfPDK1 for parasite survival, the screening of extended PDK1 inhibitor libraries as well as experimental validation of PfPDK1 as a drug target would be worthwhile activities to be pursued in future research.

## Discussion

PfPKA is essential for the proliferation of asexual blood stage parasites due to its indispensable role in phosphorylating the invasion ligand AMA1 ^5,6,8,9^. In addition, recent research described important roles for the phosphodiesterase PfPDEβ and the adenylyl cyclase PfACβ in regulating cAMP levels and hence PfPKA activity ^8,27^. Here, we studied the function and regulation of the catalytic PfPKA subunit PfPKAc to obtain further insight into PfPKA-dependent signalling in *P. falciparum* blood stage parasites. Using the NF54/AP2-G-mScarlet/PKAc cKD parasite line, we confirmed the previously described essential role for PfPKAc in merozoite invasion ^8,9^. We also demonstrated that PfPKAc plays no major role in regulating sexual commitment. The dispensability of PfPKAc in the sexual commitment pathway seems rather surprising since early studies performed over three decades ago claimed a potential involvement of cAMP signalling in regulating sexual commitment ^68,69^. However, these studies only indirectly suggested an involvement of cAMP/PfPKA-signalling in this process. For instance, Kaushal et al. determined the effect of high exogenous cAMP concentrations (1 mM) on sexual commitment and reported that under static culture conditions (high parasitaemia without addition of uRBCs) nearly all parasites developed into gametocytes ^68^. Rather than reflecting the true induction of sexual commitment by cAMP signalling, we suspect that observations may have been related to the selective killing of asexual stages by high cAMP concentrations (as indeed reported in their study) and/or the stimulation of high sexual commitment rates due to LysoPC depletion from the growth medium at high parasitaemia ^3^. Our results further suggest that PfPKAc is not required for the morphological maturation of gametocytes and for male gametogenesis. However, even though our cKD system allowed for efficient depletion of PfPKAc expression, we cannot exclude the possibility that residual PfPKAc expression levels still supported normal sexual development. At this point, we would also like to reiterate that our experiments conducted with the NF54/AP2-G-mScarlet/PKAc cKD line showed that GlcN (2.5 mM), but not Shield-1 (675 nM), acts as a confounding factor when studying sexual commitment and male gametocyte exflagellation. We therefore advise to use the FKBP/DD-Shield-1 ^36,37^ or DiCre/rapamycin ^70-72^ conditional expression systems when studying these processes.

The cellular rigidity of immature *P. falciparum* gametocytes is linked to a dynamic reorganization of the RBC spectrin and actin networks ^44^ as well as the presence of parasite-encoded STEVOR proteins at the iRBC membrane ^46,73^. In contrast, the increased deformability gained by stage V gametocytes is accompanied by the reversal of these cytoskeletal rearrangements ^44^ and dissociation of STEVOR from the iRBC membrane ^46,73^. Interestingly, results obtained from experiments using pharmacological agents to increase cellular cAMP levels or to inhibit PKA activity demonstrated that gametocyte rigidity is positively regulated by cAMP/PKA-dependent signalling ^30,73^. While potential PKA substrates involved in this process are largely unknown, the PKA-dependent phosphorylation of the cytoplasmic tail of STEVOR (S324) is important to maintain cellular rigidity of immature gametocytes and dephosphorylation of this residue is linked to the increased deformability of stage V gametocytes ^73^. Given that PfPKA is not known to be exported into the iRBC cytosol, however, PKA-dependent phosphorylation events in the RBC compartment are likely exerted by human rather than parasite PKA. Notably though, overexpression of the regulatory subunit PfPKAr, which is expected to lower PfPKAc activity, caused increased deformability of stage III gametocytes ^30^. Consistent with these data, we demonstrated that PfPKAc depletion caused a significant, yet only moderate, increase in stage III and V gametocyte deformability. While these results provide direct evidence for a role of PfPKAc-dependent phosphorylation in regulating the biomechanical properties of gametocyte-infected RBCs, they also suggest that cAMP signalling through PfPKA is not the only driver of this process. We envisage that PfPKAc activity may regulate the expression, trafficking or function of parasite-encoded proteins destined for export into the iRBC or of proteins of the inner membrane complex and/or microtubular and actin networks underneath the parasite plasma membrane that play important roles in determining cellular shape during gametocytogenesis ^74-76^. Comparative phosphoproteomic analyses of the conditional PfPKAc mutants generated here and elsewhere ^8,9^ may be a promising approach to test this hypothesis and identify the actual substrates involved.

Three recent studies employing DiCre-inducible KO parasites for PfPKAc ^8,9^, the phosphodiesterase PfPDEβ ^27^, and the adenylyl cyclase PfACβ ^8^ highlighted the importance for tight regulation of PfPKAc activity in asexual blood stage parasites. In these studies, induction of the corresponding gene KOs in ring stage parasites caused no immediate defects in intra-erythrocytic parasite development but resulted in a complete or severe block in RBC invasion by newly released merozoites due to prevention of PfPKAc activity (PfPKAc and PfACβ KOs) ^8,9^ or PfPKAc hyper-activation (PfPDEβ KO) ^27^, respectively. These findings are consistent with the specific expression pattern of all three cAMP signaling components in late schizont stages ^77^. Interestingly, however, Flueck and colleagues showed that some PfPDEβ KO merozoites successfully invaded RBCs but were then unable to develop into ring stage parasites ^27^, providing compelling evidence for a lethal effect of PfPKAc hyper-activity also on early intra-erythrocytic parasite development. Similarly, we showed that the OE of PfPKAc through a constitutively active heterologous promoter blocked parasite progression through schizogony. We believe this detrimental effect is due to illegitimate activity of PfPKAc and hence untimely phosphorylation of substrates prior to the intrinsic PfPKA activity window in late schizonts. While we did not engage in further explorations towards identifying the molecular mechanisms underlying the lethal consequences of PfPKAc overexpression, we discovered the upstream kinase that is required for PfPKAc activation. We identified this function by selecting for “PfPKAc OE survivor” parasites able to tolerate PfPKAc OE. All six independently selected NF54/PKAc cOE survivor populations carried mutations in the same gene encoding a putative serine/threonine kinase (Pf3D7_1121900). Bioinformatic analyses and structural modelling suggested this kinase is an orthologue of the eukaryotic phosphoinositide-dependent protein kinase 1 (PDK1), hence termed PfPDK1.

Interestingly, all PfPDK1 mutations identified in the various NF54/PKAc cOE survivors are positioned proximal to the ATP-binding cleft and do not coincide with the PIF-binding pocket, suggesting that these mutations impair the catalytic efficiency of PfPDK1 rather than its capacity to interact with substrates. It therefore seems that the most straightforward manner for the parasite to overcome the lethal effect of PfPKAc OE was to acquire mutations reducing PfPKAc activation through PfPDK1-mediated phosphorylation. We confirmed this scenario by (1) reverting the PfPDK1 M51R mutation in the NF54/PKAc cOE S1 survivor, which rendered these parasites again sensitive to PfPKAc OE; and (2) by introducing the PfPDK1 M51R mutation into the PfPKAc OE-sensitive clone M1, which rendered these parasites resistant to PfPKAc OE. Notably, we could also show that OE of the PfPKAcT189V mutant form of PfPKAc, which carries a non-phosphorylatable valine residue instead of the target threonine in the activation loop, had no negative effect on intra-erythrocytic parasite development and multiplication. Together, these striking results demonstrate that in addition to the cAMP-mediated release of PfPKAc from the regulatory subunit PfPKAr, phosphorylation of the T189 residue is essential for PfPKAc activity, and provide compelling evidence that PfPDK1 is the kinase that targets this residue. Conditional depletion of PfPDK1 did not result in any obvious multiplication or developmental defects in asexual and sexual blood stage parasites, showing that largely diminished PfPDK1 protein levels are still sufficient to sustain parasite viability and proliferation. However, several lines of evidence strongly argue for an essential role for PfPDK1 in asexual parasites and that at least one of its vital functions is to activate PfPKAc. First, previous studies ^31,53^ and our own attempts failed to obtain PfPDK1 null mutants via gene disruption approaches. Second, none of the different mutations identified in the six NF54/PKAc cOE survivors introduced a nonsense loss-of-function mutation into the *pfpdk1* open reading frame. This observation again supports the notion that PfPDK1 function is vital and that the PfPDK1 mutant enzymes retain residual kinase activity. Third, we were only able to introduce the M51R PfPDK1 mutation into parasites overexpressing PfPKAc but not into WT parasites, suggesting that parasites expressing functionally compromised PfPDK1 mutants can only survive if impaired PfPDK1-dependent PfPKAc activation is compensated for by elevated PfPKAc expression levels. Given that PDK1 is widely conserved in eukaryotes and required to activate AGC kinases at large ^17^, it is conceivable that PfPDK1 may also regulate the activity of PfPKG or PfPKB, the only other two known members of the AGC family in *P. falciparum*, which are both essential in blood stage parasites ^31,78^. Furthermore, the manifestation of the PfPKAc OE phenotype in late trophozoites/early schizonts implies that PfPDK1 is active and phosphorylates other substrates already at this stage, hours prior to the expression window of endogenous PfPKAc in late schizonts. Importantly, however, the fact that the NF54/PKAc cOE survivor parasites expressing functionally impaired PfPDK1 mutant enzymes are fully viable suggests that the PfPDK1-dependent phosphorylation of other substrates is either not essential or can still be executed at functionally relevant baseline levels by mutated PfPDK1.

In summary, we provide unprecedented functional insight into the cAMP/PfPKA signalling pathway in the malaria parasite *P. falciparum*. Our results complement earlier studies highlighting the importance of tight regulation of PfPKA activity for parasite survival, showing that diminished as well as augmented PfPKAc expression levels are lethal for asexual blood stage parasites. In addition to the well-established roles for the regulatory subunit PfPKAr, the adenylyl cyclase PfACβ and the phosphodiesterase PfPDEβ in regulating PfPKAc activity via cAMP levels, we identified PfPDK1 as the upstream kinase activating PfPKAc, most likely through activation loop phosphorylation at T189. In light of the essential role for PfPDK1 in this and possibly other parasite AGC kinase-dependent signalling pathways, and the promising anti-parasite activity of PDK1 kinase inhibitors, PfPDK1 represents an attractive candidate for further functional and structural studies and to be explored as a possible new antimalarial drug target.

## Methods

### Parasite culture

*P. falciparum* NF54 parasites were cultured and asexual growth was synchronized using 5% sorbitol as described previously ^79,80^. Parasites were cultured in AB+ or B+ human RBCs (Blood Donation Center, Zurich, Switzerland) at a hematocrit of 5%. The standard parasite culture medium (PCM) contains 10.44 g/L RPMI-1640, 25 mM HEPES, 100 μM hypoxanthine and is complemented with 24 mM sodium bicarbonate and 0.5% AlbuMAX II (Gibco #11021-037). 2 mM choline chloride (CC) was routinely added to PCM to block induction of sexual commitment ^3^. To induce sexual commitment, parasites were cultured in serum-free medium (–SerM) as described previously ^3^. –SerM medium contains fatty acid-free BSA (0.39%, Sigma #A6003) instead of AlbuMAX II and 30 μM oleic and 30 μM palmitic acid (Sigma #O1008 and #P0500) ^3^. Sexual commitment rate assays were performed using standardised –SerM medium complemented with 2 mM choline chloride (–SerM/CC) ^3^. Gametocytes were cultured in PCM complemented with 10% human serum (Blood Donation Center, Basel, Switzerland) instead of AlbuMAX II (+SerM). Parasite cultures were kept in gassed (4% CO_2_, 3% O_2_, 93% N_2_) airtight containers at 37 °C.

### Transfection constructs

The NF54/AP2-G-mScarlet/PKAc cKD parasite line was generated using a SLI-based gene editing approach ^35^. For this purpose, the SLI_PKAc_cKD transfection construct was generated in three successive cloning steps using Gibson assembly reactions. First, the SLI_cKD precursor plasmid was generated by joining four fragments in a Gibson assembly reaction: (1) the plasmid backbone amplified from pUC19 (primers PCRA_F and PCRA_R) as previously described ^52^; (2) the *SpeI/BamHI* cloning site followed by the *gfp-dd* sequence amplified from pD_cOE_DD_SpeI/BamHI (Hitz et al., unpublished) using primers gfp_F and dd_R; (3) the *2a*-*bsd* sequence amplified from pSLI-BSD ^35^ (primers 2A_F and bsd_R); and (4) the *glmS*-*hrp2* 3’ sequence amplified from pD_cOE (described below) (primers glmS1_F and term_R). Second, a four-fragment Gibson assembly reaction was performed using (1) the *SalI-* and *EcoRI*-digested SLI_cKD precursor plasmid; (2) the *calmodulin* (PF3D7_1434200) promoter amplified from pHcamGFP-DD ^52^ (primers cam1_F and cam1_R); (3) the yeast dihydroorotate dehydrogenase (y*dhodh*) resistance gene (conferring resistance to DSM1) amplified from the pUF1-Cas9 plasmid ^81^ (primers ydhodh_F and ydhodh_R); and (4) the *pbdt 3’* terminator amplified from pUF1-Cas9 ^81^ (primers pbdt3_F and pbdt3_R). Third, this SLI_cKD_ydhodh precursor plasmid was digested using *SpeI* and *BamHI* and joined with the 3’ HR of *pfpkac* amplified from NF54 gDNA (primers pka3’_F and pka3’_R) in a two-fragment Gibson assembly reaction resulting in the final SLI_PKAc_cKD transfection construct.

The CRISPR/Cas9 gene editing system employed here is based on co-transfection of suicide and donor plasmids ^52^. The pBF-gC- or pHF-gC-derived suicide plasmids encode the Cas9 enzyme, the single guide RNA (sgRNA) cassette and either the blasticidin deaminase (BSD; conferring resistance to BSD-S-HCl) or the human dihydrofolate reductase (hDHFR; conferring resistance to WR99210) resistance markers fused to the negative selection marker yeast cytosine deaminase/uridyl phosphoribosyl transferase (BSD-yFCU or hDHFR-yFCU) ^52^. The pD-derived donor plasmid ^52^ contains the sequence assembly essential for homology-directed repair of the DNA double-strand break induced by Cas9. To obtain the NF54/PKAc cOE parasite line, the pD_pkac_cOE donor plasmid was generated in several cloning steps using Gibson assembly reactions. First, a pD_cOE_DD precursor plasmid was generated using a two-fragment Gibson assembly joining (1) the plasmid backbone including the *glp3*-specific 5’ and 3’ HRs amplified from pD_*cg6_cam-gdv1-gfp-glmS* ^82^ (primers glp3_F and glp3_R) and (2) the *cam 5’-gfpdd-hrp2 3’* sequence amplified from pHcamGFP-DD ^52^ (primers cam_F and hrp2_R). Second, the subsequent precursor plasmid pD_cOE_glmS was generated by performing a Gibson assembly using two fragments (1) the pD_cOE_DD precursor plasmid digested using *SalI* and *AgeI* and the (2) *glmS* sequence amplified from pD_*cg6_cam-gdv1-gfp-glmS* ^82^ (primers glmS_F and glmS_R). To insert a new cloning site and generate the next precursor plasmid pD_cOE, another Gibson assembly reaction was performed joining (1) the pD_cOE_glmS plasmid digested using *BamHI* and *NotI* and (2) annealed complementary oligonucleotides clon_F and clon_R. The final pD_pkac_cOE plasmid was generated by assembling two Gibson fragments: (1) the *BamHI-* and *SpeI-*digested pD_cOE plasmid and (2) the *pfpkac* sequence amplified from NF54 gDNA using primers pka_F and pka_R. The pBF_gC-*cg6* suicide plasmid that was co-transfected with pD_pkac_cOE to generate this NF54/PKAc cOE parasite line encodes the sgRNA targeting the *glp3* locus ^82^. To obtain the NF54/PKAcT189V cOE parasite line, the pD_pkacT189V_cOE donor plasmid was generated in a Gibson assembly joining three fragments: (1) the *BamHI-* and *SpeI-*digested pD_cOE plasmid; (2) the 5’ fragment of the *pfpkac* sequence containing the single point mutation resulting in the T189V amino acid change (primers pka_F and T189V_R); and (3) the 3’fragment of the *pfpkac* sequence overlapping with the 5’ fragment and coding for the same amino acid change (primers T189V_F and pka_R). As described above, the *glp3*-specific suicide plasmid pBF_gC-*cg6* was used for co-transfection. The NF54/PDK1 cKD parasite line was obtained by co-transfection of pHF_gC_pdk1-gfpdd (encoding sgRNA_pdk1) and pD_pdk1-gfpdd. The previously published suicide mother plasmid pHF-gC ^52^ was used to insert the sgRNA sequence targeting the 3’ end of *pfpdk1* (sgt_pdk1). For this purpose, complementary oligonucleotides were annealed and the resulting double-stranded fragment was ligated into the *BsaI*-digested pHF-gC plasmid using T4 DNA ligase generating the pHF_gC_pdk1-gfpdd suicide plasmid. The pD_pdk1-gfpdd plasmid was generated by performing a four-fragment Gibson assembly joining (1) the plasmid backbone amplified from pUC19 (primers PCRA_F and PCRA_R) as previously described ^52^; (2) the 5’ HR amplified from NF54 gDNA (primers hr1KD_F and hr1KD_R); (3) the *gfpdd* sequence (primers gfpdd_F and gfpdd_R) amplified from pHcamGDV1-GFP-DD ^52^; and (4) the 3’ HR amplified from NF54 gDNA (hr2KD_F and hr2KD_R). The parasite lines NF54/PKAc cOE M1/PDK1_mut and NF54/PKAc cOE S1/PDK1_wt were generated as follows. Plasmids pHF_gC_S1rev and pHF_gC_M1mut were generated by inserting annealed complementary oligonucleotides encoding the respective sgRNAs (sgRNA_S1rev or sgRNA_M1mut) into the *BsaI*-digested pHF_gC suicide vector ^52^. The donor plasmids pD_S1rev and pD_M1mut were generated in a three-fragment Gibson assembly joining (1) the plasmid backbone amplified from pUC19 (primers PCRA_F and PCRA_R) as previously described ^52^; (2) the corresponding 5’ HRs amplified from NF54 gDNA (primers hr1_F and either rev_R or mut_R); and (3) the respective 3’ HRs amplified from NF54 gDNA (primers hr2_R and either rev_F or mut_F). The primers rev_F/rev_R or mut_F/mut_R encode the WT (M) or mutated (R) amino acid number 51 of PfPDK1, respectively.

All primers used for cloning of the described transfection constructs are listed in Supplementary Table 1.

### Transfection and transgenic cell lines

*P. falciparum* ring stage parasite transfection was performed as described ^52^. A total of 100 µg plasmid DNA was used to transfect NF54/AP2-G-mScarlet and NF54 WT parasites (100 µg of the SLI_PKAc_cKD plasmid; 50 µg each of all described CRISPR/Cas9 suicide and donor plasmids). 24 hours after transfection of the SLI_PKAc_cKD plasmid, parasites were cultured on 1.5 µM DSM1 until a stably growing parasite population was obtained. This culture was subsequently treated with 2.5 μg/mL BSD-S-HCl to select for parasites in which the *pfpkac* gene was successfully tagged. Similarly, 24 hours after transfection of CRISPR/Cas9-based plasmids, the cultures were treated with 2.5 μg/mL BSD-S-HCl (for ten subsequent days) or 4 nM WR99210 (for six subsequent days) depending on the resistance cassette encoded by the suicide plasmid. NF54/AP2-G-mScarlet/PKAc cKD and NF54/PDK1 cKD parasites were constantly cultured on 675 nM Shield-1 (+Shield-1) to stabilize the PfPKAc-GFPDD or PfPDK1-GFPDD protein, respectively. NF54/PKAc cOE, NF54/PKAcT189V cOE, NF54/PKAc cOE M1/PDK1_mut and NF54/PKAc cOE S1/PDK1_wt parasites were constantly cultured on 2.5 mM GlcN to block OE of PfPKAc or PfPKAcT189V. About two to three weeks after transfection, stably growing parasite cultures were obtained and diagnostic PCRs on gDNA were used to confirm correct genome editing. Primers used to verify for correct gene editing are listed in Supplementary Table 2. Correct targeted mutagenesis of the *pfpdk1* gene in NF54/PKAc cOE M1/PDK1_mut and NF54/PKAc cOE S1/PDK1_wt parasites was confirmed by Sanger sequencing. Sanger sequencing data analysis and visualisation was performed using SnapGene software 4.1.6 (Insightful Science).

### Limiting dilution cloning

Limiting dilution cloning was performed as previously described ^48^. In brief, synchronous ring stage parasite cultures were diluted with fresh PCM and RBCs to a haematocrit of 0.75 % and a parasitaemia of 0.0006 % (=parasite cell suspension). Each well of a flat-bottom 96 well microplate (Costar #3596) was filled with 200 µL PCM/0.75% haematocrit (=RBC suspension). In each well of row A, 100 µL of the parasite cell suspension was mixed with the 200 µL RBC suspension (1/3 dilution) resulting in a parasitaemia of 0.0002% which equals approximately 30 parasites per well. Subsequently, 100 µL of the row A parasite cell suspensions were mixed with 200 µL RBC suspension in the wells of row B resulting again in a 1/3 dilution (approximately 10 parasites/well). This serial dilution was continued until the last row of the plate was reached. The 96-well microplate was kept in a gassed airtight container at 37 °C for 11-14 days without medium change. Subsequently, using the Perfection V750 Pro scanner (Epson), the 96-well microplate was imaged to visualize plaques in the RBC layer. The content of wells containing a single plaque was then transferred individually into 5 mL cell culture plates and cultured using PCM until a stably growing parasite culture was obtained.

### Flow cytometry

Quantification of parasite multiplication was performed using flow cytometry measurements of fluorescence intensity. For this purpose, synchronous parasites cultures (0.2 % parasitaemia) were split at 0-6 hpi and cultured separately for the entire duration of the multiplication assay under (i) +Shield-1/–GlcN and –Shield-1/+GlcN conditions (NF54/AP2-G-mScarlet/PKAc cKD); (ii) –GlcN and +GlcN conditions (NF54/PKAc cOE M1, NF54/PKAc cOE S1, NF54/PKAc cOE S1/PDK1_wt, NF54/PKAc cOE M1/PDK1_mut, NF54/PKAcT189V cOE, NF54 WT); or (iii) +Shield-1 and –Shield-1 conditions (NF54/PDK1 cKD, NF54 WT). To determine the exact starting parasitaemia on day 1 of the assay (generation 1), gDNA of synchronous ring stage parasites (18-24 hpi) was stained for 30 min using SYBR Green DNA stain (Invitrogen, 1:10,000) at 37 °C. Subsequently, parasites were washed twice in PBS and fluorescence intensity was measured using the MACS Quant Analyzer 10. 200,000 RBCs were measured per sample. The measurement was repeated for two (day 3 and 5) or three (day 3, 5 and 7) subsequent generations. The FlowJo_v10.6.1 software was used to analyse the flow cytometry data. The measurements were gated to remove small debris (smaller than cell size) and doublets (two cells in a single measurement). Using an uninfected RBC (uRBC) control sample, uRBCs were separated from iRBCs based on their SYBR Green intensity. Representative plots showing the gating strategy used for all flow cytometry data are presented in Supplementary Fig. 2.

### Fluorescence microscopy

Live cell fluorescence imaging was performed to visualize protein expression as described ^83^. Parasite nuclei were stained using 5 µg/ml Hoechst (Merck) and Vectashield (Vector Laboratories) was used to mount the microscopy slides. Live cell fluorescence microscopy was performed using a Leica DM5000 B fluorescence microscope (20x, 40x and 63x objectives) and images were acquired using the Leica application suite (LAS) software Version 4.9.0 and the Leica DFC345 FX camera. Images were processed using Adobe Photoshop CC 2018 and for each experiment identical settings for both image acquisition and processing were used for all samples analysed.

For the quantification of the number of nuclei per schizont, synchronous NF54/PfPKAc cOE M1 and S1 ring stage parasites (0-6 hpi) were split and cultured either in presence or absence GlcN. 40 hours later (40-46 hpi), schizonts were stained using 5 µg/ml Hoechst and Vectashield-mounted slides visually inspected using the Leica fluorescence microscope and the LAS software. Three biological replicate experiments were performed and for each replicate the number of nuclei in 100 schizonts was determined by manual counting.

### Western blot analysis

Parasite pellets were obtained by lysing RBCs using 0.15% saponin in PBS (10 min on ice) followed by centrifugation and washed in ice-cold PBS until the supernatant was clear. Whole parasite protein lysates were generated by solubilising the parasite pellet in an UREA/SDS buffer (8 M Urea, 5% SDS, 50 mM Bis-Tris, 2 mM EDTA, 25 mM HCl, pH 6.5) complemented with 1x protease inhibitor cocktail (Merck) and 1 mM DTT. Protein lysates were separated on NuPage 5-12% Bis-Tris or 3-8% Tris-Acetate gels (Novex, Qiagen) using MES running buffer (Novex, Qiagen). Following protein transfer to a nitrocellulose membrane, the membrane was blocked for 30 min using 5% milk in PBS/0.1% Tween (PBS/Tween). Primary antibodies mouse mAb α-GFP (1:1,000) (Roche Diagnostics #11814460001) or mAb α-PfGAPDH ^84^ (1:20,000) diluted in blocking buffer were used for detection of GFP-tagged proteins or the GAPDH loading control, respectively. Primary antibody incubation was performed at 4 °C overnight in PBS/Tween and the membrane was subsequently washed three times. Incubation using the secondary antibody α-mouse IgG (H&L)-HRP (1:10,000) (GE healthcare #NXA931) diluted in blocking buffer was performed for two hours. The membrane was washed three times using PBS/Tween before chemiluminescent signal detection using KPL LumiGLO Chemiluminescent Substrate System (SeraCare).

### Quantification of sexual commitment rates

High content imaging was performed to quantify sexual commitment rates (SCRs) of NF54/AP2-G-mScarlet/PKAc cKD and NF54/AP2-G-mScarlet control parasites. Synchronous parasites (0-6 hpi) were split (±Shield-1/±GlcN or ±GlcN) and 18 hours later (18-24 hpi) the culture medium was replaced with standardised –SerM/CC medium (2% parasitaemia, 2.5% hematocrit). Twenty-two hours later (sexually committed schizonts) or 48 hours later (sexually committed ring stage progeny), 30 µL of culture suspension were mixed with 50 µL Hoechst/PBS (8.1 µM) and incubated in a 96-well plate for 30 min. Cells were pelleted (300 g, 5 min) and washed twice using 200 µL PBS. Subsequently, the cell pellet was resuspended using 180 µL PBS and 30 µL suspension was pipetted into the wells of a clear-bottom 96-well plate (Greiner CELLCOAT microplate 655948, Poly-D-Lysine, flat μClear bottom) containing 150 µL PBS per well. Prior to imaging, cells were allowed to settle for 30 min at 37 °C. The MetaXpress software (version 6.5.4.532, Molecular Devices), the ImageXpress Micro XLS widefield high content imaging system using a Plan-Apochromat 40x objective and Sola SE solid state white light engine (Lumencor) were used for automated image acquisition and data analysis. The Hoechst (Ex: 377/50 nm, Em: 447/60 nm, 80 ms exposure) and mScarlet (Ex: 543/22 nm, Em: 593/40 nm, 600 ms exposure) filter sets were used for imaging all iRBCs and AP2-G-mScarlet-expressing parasites, respectively. Thirty-six sites per well were imaged. Quantification of both Hoechst-positive and AP2-G-mScarlet-positive parasites allowed calculating SCRs (percentage of AP2-G-mScarlet-positive parasites amongst all Hoechst-positive parasites).

To quantify SCRs of NF54/PDK1 cKD parasites, N-acetyl-D-glucosamine (GlcNAc) assays were performed ^43^. For this purpose, synchronous parasites (0-6 hpi) were split (±Shield-1) and 18 hours later (18-24 hpi) the culture medium was replaced with standardised –SerM/CC medium (2% parasitaemia, 5% haematocrit). Upon reinvasion, the ring stage parasitaemia was quantified from Giemsa-stained thin blood smears prepared at 18-24 hpi. This parasitaemia corresponds to the cumulative counts of asexual ring stages and sexual ring stages (day one of gametocytogenesis). From 24-30 hpi onwards, parasites were cultured in +SerM medium supplemented with 50 mM GlcNAc (Sigma) to eliminate asexual parasites ^43^. On day four of gametocytogenesis (stage II gametocytes), the parasitaemia was again quantified from Giemsa-stained thin blood smears. SCRs were determined as the percentage of the day four parasitaemia (stage II gametocytes) compared the total parasitaemia observed on day one.

### Gametocyte cultures

Synchronous gametocyte cultures were used to study gametocyte morphology, to extract protein samples and to perform microsphiltration and exflagellation assays. Sexual commitment was induced at 18-24 hpi using –SerM medium. Upon reinvasion (0-6 hpi) (asexual and sexual ring stages; day one of gametocytogenesis), parasites were cultured in +SerM medium. Another 24 hours later (24-30 hpi) (trophozoites and stage I gametocytes, day two of gametocytogenesis), 50 mM GlcNAc was added (+SerM/GlcNAc) to eliminate asexual parasites ^43^. The +SerM/GlcNAc medium was changed daily for six consecutive days and thereafter gametocytes were cultured in +SerM medium that was replaced daily on a 37 °C heating plate to prevent gametocyte activation.

### Microsphiltration experiments

Synchronous NF54/AP2-G-mScarlet/PKAc cKD and NF54 WT parasites were split at 0-6 hpi and cultured separately in presence and absence of Shield-1 (±Shield-1). Subsequently, sexual commitment was induced at 18-24 hpi using –SerM medium and after reinvasion gametocytes were cultured using +SerM/GlcNAc and +SerM medium as described above. On day six (stage III) and eleven (mature stage V) of gametocytogenesis, microsphiltration experiments were conducted as described previously ^45^. Per sample and condition, either one (NF54 WT) or two (NF54/AP2-G-mScarlet/PKAc cKD) independent biological replicate experiments with six technical replicates each were conducted. The experiment starts by transferring gametocyte culture aliquots into 15 mL Falcon tubes and lowering the haematocrit to 1.5% by addition of PCM. Six microsphere filters (technical replicates) were loaded per sample and condition. After injection of 600 µL cell suspension, filters were washed with 5 mL +SerM medium at a speed of 60 mL per hour using a medical grade pump (Syramed µSP6000, Acromed AG, Switzerland). Gametocytaemia before (“UP”) and after (“DOWN”) the microsphiltration process were determined from Giemsa-stained thin blood smears by counting at least 1,000 RBCs. The “UP” gametocytaemia was determined as the mean gametocytaemia calculated from two independent Giemsa-stained thin blood smears. The “DOWN” gametocytaemia was determined for each filter separately. Gametocyte retention rates were calculated as 1-(“DOWN” gametocytaemia divided by “UP” gametocytaemia). Samples were kept at 37 °C whenever possible to prevent gametocyte activation and lack thereof was confirmed by visual inspection of Giemsa-stained thin blood smears.

### Exflagellation assays

On day 14 of gametocytogenesis (mature stage V), exflagellation assays were performed as described previously ^85^. Briefly, in a Neubauer chamber gametocytes were activated using 100 µM xanthurenic acid (XA) and a drop in temperature (from 37 °C to 22 °C). After 15 min of activation, the total number of RBCs per mL of culture and the number of exflagellation centers by activated male gametocytes were quantified by bright-field microscopy (40x objective). The gametocytaemia before activation was determined from Giemsa-stained thin blood smears. Exflagellation rates were calculated as the proportion of exflagellating gametocytes among all gametocytes. At least three biological replicates were performed per experiment.

### Illumina whole genome sequencing

To perform WGS, gDNA of the NF54/PKAc cOE clones M1 and M2 and the six independently grown NF54/PKAc cOE survivors (S1-S6) was isolated using a phenol/chloroform-based extraction protocol as described ^86^. To avoid an amplification bias due to the high AT-content of *P. falciparum* gDNA, DNA sequencing libraries were prepared using the PCR-free KAPA HyperPrep Kit (Roche). Libraries were sequenced on an Illumina NextSeq 500 and the quality of the raw sequencing reads was analysed with FastQC (version 0.11.4) ^87^. The raw reads were mapped to the *P. falciparum* 3D7 reference genome (PlasmoDB version 39) complemented with the corresponding transfection plasmid sequences using the Burrows-Wheeler Aligner (version 0.7.17) ^88^ with default parameters. The alignment files in SAM format were converted to binary BAM files with SAMtools (version 1.7) ^89^ and the BAM files were coordinate-sorted, indexed and read-groups were added with Picard (version 2.6.0) ^90^. Sequence variants (SNP and Indels) were directly called with the Genome Analysis Toolkit’s (GATK, version 4.0.7.0) HaplotypeCaller in the GVCF mode to allow multi-sample analysis ^91^. The resulting g.vcf files of the different samples were combined into one file and genotyped using GATK ^91^. To predict the consequences of the obtained variants, they were annotated with SnpEff (version 4.3T) ^92^ using the SnpEff database supplemented manually with the 3D7 genome annotation (PlasmoDB version 39). The detected variants were filtered for (i) “HIGH” or “MODERATE” impact (thus non-synonymous variants); (ii) absence in NF54/PKAc cOE clones M1 and M2; and (iii) an allele frequency of the alternative allele of >40% in at least one of the NF54/PKAc cOE survivors (S1-S6). The obtained list of 83 candidate variants was first screened manually (i) for variants present in all NF54/PKAc cOE survivors; and (ii) for genes mutated in all NF54/PKAc cOE survivors, leaving the variants identified in Pf3D7_1121900 as only candidates. Additionally, all 83 original candidate variants were inspected visually with the Integrative Genomics Viewer (version 2.7.0) ^93^. Variants were excluded if they were suspected to be false-positives because they were (i) only supported by a very small number of reads, and (iia) also detected in reads of the mother clones and not called because of low allele frequencies or (iib) insertions and deletions after/before large homopolymers or repeat tracts. This again left the variants in Pf3D7_1121900 as only candidates.

To analyse PfPKAc-GFP cOE cassette copy numbers, the sequencing coverage over the whole genome was determined in 50-nucleotide windows using the software igvtools (version 2.5.3) ^93^ and the mean coverage of (i) the whole genome, (ii) the endogenous *pfpkac* locus and (iii) the ectopic *pfpkac-gfp* cOE cassettes were calculated in RStudio (R version 3.6.2, RStudio release 1.2.5033). The mean coverage of the endogenous *pfpkac* locus and ectopic *pfpkac-gfp* cOE cassettes were summed up, as both sequences are identical and reads derived from the ectopic *pfpkac-gfp* cOE cassettes were mapped to both alleles. *pfpkac* coverage was normalized to the mean genome-wide coverage (assuming the copy number of the genome is 1) and to the *pfpkac* coverage of WT parasites. Finally, the endogenous *pfpkac* was subtracted (−1) to obtain the approximate copy number of ectopic PfPKAc-GFP cOE cassettes.

### Sequence alignments and modelling of protein structure

Alignment of the PfPDK1 (Pf3D7_1121900/UniProt ID Q8IIE7) amino acid sequence with PDK1 from *P. vivax* (PVX_091715, UniProt ID A5K4N1), *Arabidopsis thaliana* (UniProt ID Q9XF67), *Caenorhabditis elegans* (UniProt ID Q9Y1J3) and humans (UniProt ID O15530) was performed using Clustal Omega ^94^. A homology model of the PfPDK1 structure was built using SWISS-MODEL ^95^ based on the human PDK1 crystallographic structure (PDB ID 1UU9) ^51^. The mean homology model quality (Global Model Quality Estimation, GMQE) was assessed as 0.49, suggesting a model of average quality (possible GMQE values are 0-1 on a linear scale, higher values indicate better quality). A large, predicted disordered loop (amino acids 180-310) present in PfPDK1 but not in its homologues was not built in the model. Amino acids of the ATP-binding cleft were defined as those within 4 Å of any ATP atom in the human PDK1 structure. The PIF-binding pocket was defined as suggested by Biondi and colleagues ^22^.

### Drug assays

Activity of human PDK1 kinase inhibitors on asexual NF54 parasite multiplication was determined using an [^3^H] hypoxanthine incorporation assay ^67^. The mean IC_50_ values were determined from three biological replicate assays, each performed in technical duplicate. BX-795 (CAS Number 702675-74-9, Selleckchem #S1274), BX-912 (CAS Number 702674-56-4, Selleckchem #S1275 and GSK2334470 (CAS Number 1227911-45-6, Selleckchem #S7087), were resuspended in DMSO and used at a maximum starting concentration of 50 µM (replicates 1 and 2) or 10 µM (replicate 3). In a six-step serial dilution, the compound concentration was diluted to half in each step to span a concentration range between 50 µM and 0.8 µM or 10 µM and 0.15 µM, respectively. Chloroquine (CAS Number 50-63-5, Sigma #C6628) and artesunate (CAS Number 88495-63-0, Mepha #11665) served as control antimalarial compounds with a starting concentration of 100 nM and 50 nM, respectively. In a six-step serial dilution, the chloroquine and artesunate concentrations were diluted to half in each step to span a concentration range between 100 nM and 1.5 nM or between 50 nM and 0.8 nM, respectively.

### Statistical analysis

All data from assays quantifying parasite multiplication, number of nuclei per parasite, sexual commitment rates, gametocyte deformability, exflagellation rates and IC50 values are represented as means with error bars defining the standard deviation. All data were derived from at least three biological replicate experiments. Statistical significance (*p*<0.05) was determined using paired or unpaired Student’s *t* tests as indicated in the figure legends. The exact number of biological replicates performed per experiment and the number of cells analysed per sample are indicated in the figure legends and in the Source Data file. Data were analysed and plotted using RStudio Version 1.1.456 and package ggplot2 or GraphPad Prism Version 8.2.1 for Windows.

## Data availability

Whole genome sequencing data was deposited on the European Nucleotide Archive, accession number PRJEB40033. The source data for all the graphs and charts in the main and Supplementary figures are present in the Source Data file and any other remaining information can be available from the corresponding author upon reasonable request.

## Acknowledgements

We thank Christian Flueck for his valuable inputs on the manuscript and Christian Scheurer for assistance in performing the drug assays. This work was supported by funding from the Swiss National Science Foundation (BSCGI0_157729 to T.S.V. and 310030_156264 to P.M.) and the Rudolf Geigy Foundation. I.V. thanks the Medical Research Council UK (MR/N009274/1) and the EPA Cephalosporin Fund (CF 329) for their support.

## Author Contributions

E.H. produced transgenic parasite lines, designed and performed experiments, analysed and interpreted data, prepared illustrations and wrote the manuscript. N.W. analysed the WGS data and P.M. supervised these experiments and provided resources. N.M.B.B. generated the PfPKAc cKD parasite line and supervised sexual commitment experiments using high content microscopy. A.P. performed and analysed microsphiltration experiments supervised by N.M.B.B. I.V. performed the bioinformatic analysis and structure modelling of PfPDK1. T.S.V. conceived the study, designed and supervised experiments, prepared illustrations, provided resources and wrote the manuscript. All authors contributed to editing of the manuscript.

