## Supplementary Data 1 for "The 3-phosphoinositide-dependent protein kinase 1 is an essential upstream activator of protein kinase A in malaria parasites"

```
                    1                                                                                                  100
PF3D7_1121900       ATGAAGAAAGGATTTTTGTTGAATAAGAATATATATGATATTGAAGAAAATGTAATAAATGTTGAAAAAAATAACACAAATAAAAAAATATTGTAAGGATG
S1_c.152T>G         ATGAAGAAAGGATTTTTGTTGAATAAGAATATATATGATATTGAAGAAAATGTAATAAATGTTGAAAAAAATAACACAAATAAAAAAATATTGTAAGGATG
S2_c.295G>T         ATGAAGAAAGGATTTTTGTTGAATAAGAATATATATGATATTGAAGAAAATGTAATAAATGTTGAAAAAAATAACACAAATAAAAAAATATTGTAAGGATG
S3_c.295G>T         ATGAAGAAAGGATTTTTGTTGAATAAGAATATATATGATATTGAAGAAAATGTAATAAATGTTGAAAAAAATAACACAAATAAAAAAATATTGTAAGGATG
S4_c.252A>T         ATGAAGAAAGGATTTTTGTTGAATAAGAATATATATGATATTGAAGAAAATGTAATAAATGTTGAAAAAAATAACACAAATAAAAAAATATTGTAAGGATG
S5_c.134A>G         ATGAAGAAAGGATTTTTGTTGAATAAGAATATATATGATATTGAAGAAAATGTAATAAATGTTGAAAAAAATAACACAAATAAAAAAATATTGTAAGGATG
S6_c.252A>T         ATGAAGAAAGGATTTTTGTTGAATAAGAATATATATGATATTGAAGAAAATGTAATAAATGTTGAAAAAAATAACACAAATAAAAAAATATTGTAAGGATG
S6_c.491A>G         ATGAAGAAAGGATTTTTGTTGAATAAGAATATATATGATATTGAAGAAAATGTAATAAATGTTGAAAAAAATAACACAAATAAAAAAATATTGTAAGGATG

                    101                                                                                                200
PF3D7_1121900       ATTTTGAAATATATATGCACATAGGAACTGGGAATTTCAGTGATGTGTTTATGGTAAAATTAAAAAATGATCCTTCAAAAAAATATGCCTTAAAAATATT
S1_c.152T>G         ATTTTGAAATATATATGCACATAGGAACTGGGAATTTCAGTGATGTGTTTAGGGTAAAATTAAAAAATGATCCTTCAAAAAAATATGCCTTAAAAATATT
S2_c.295G>T         ATTTTGAAATATATATGCACATAGGAACTGGGAATTTCAGTGATGTGTTTATGGTAAAATTAAAAAATGATCCTTCAAAAAAATATGCCTTAAAAATATT
S3_c.295G>T         ATTTTGAAATATATATGCACATAGGAACTGGGAATTTCAGTGATGTGTTTATGGTAAAATTAAAAAATGATCCTTCAAAAAAATATGCCTTAAAAATATT
S4_c.252A>T         ATTTTGAAATATATATGCACATAGGAACTGGGAATTTCAGTGATGTGTTTATGGTAAAATTAAAAAATGATCCTTCAAAAAAATATGCCTTAAAAATATT
S5_c.134A>G         ATTTTGAAATATATATGCACATAGGAACTGGGAGTTTCAGTGATGTGTTTATGGTAAAATTAAAAAATGATCCTTCAAAAAAATATGCCTTAAAAATATT
S6_c.252A>T         ATTTTGAAATATATATGCACATAGGAACTGGGAATTTCAGTGATGTGTTTATGGTAAAATTAAAAAATGATCCTTCAAAAAAATATGCCTTAAAAATATT
S6_c.491A>G         ATTTTGAAATATATATGCACATAGGAACTGGGAATTTCAGTGATGTGTTTATGGTAAAATTAAAAAATGATCCTTCAAAAAAATATGCCTTAAAAATATT

                    201                                                                                                300
PF3D7_1121900       CAAAAAAGAAAAAGTGAATAAAATGAATAAAGTAAATGATGTGTTAACAGAAAAAAACGTTATGTCAAAATTAAATACCCCAGGACATGCAAATGTTATT
S1_c.152T>G         CAAAAAAGAAAAAGTGAATAAAATGAATAAAGTAAATGATGTGTTAACAGAAAAAAACGTTATGTCAAAATTAAATACCCCAGGACATGCAAATGTTATT
S2_c.295G>T         CAAAAAAGAAAAAGTGAATAAAATGAATAAAGTAAATGATGTGTTAACAGAAAAAAACGTTATGTCAAAATTAAATACCCCAGGACATGCAAATTTTATT
S3_c.295G>T         CAAAAAAGAAAAAGTGAATAAAATGAATAAAGTAAATGATGTGTTAACAGAAAAAAACGTTATGTCAAAATTAAATACCCCAGGACATGCAAATTTTATT
S4_c.252A>T         CAAAAAAGAAAAAGTGAATAAAATGAATAAAGTAAATGATGTGTTAACAGATAAAAACGTTATGTCAAAATTAAATACCCCAGGACATGCAAATGTTATT
S5_c.134A>G         CAAAAAAGAAAAAGTGAATAAAATGAATAAAGTAAATGATGTGTTAACAGAAAAAAACGTTATGTCAAAATTAAATACCCCAGGACATGCAAATGTTATT
S6_c.252A>T         CAAAAAAGAAAAAGTGAATAAAATGAATAAAGTAAATGATGTGTTAACAGATAAAAACGTTATGTCAAAATTAAATACCCCAGGACATGCAAATGTTATT
S6_c.491A>G         CAAAAAAGAAAAAGTGAATAAAATGAATAAAGTAAATGATGTGTTAACAGAAAAAAACGTTATGTCAAAATTAAATACCCCAGGACATGCAAATGTTATT

                    301                                                                                                400
PF3D7_1121900       AAATTAATAGAAACATTTAAAGACAAAGAAAATGTTTATTTATTATATGAATATGCTGATTATGATTTATGGGAATTTTTAAAAATTCGTAGTGTTGGTG
S1_c.152T>G         AAATTAATAGAAACATTTAAAGACAAAGAAAATGTTTATTTATTATATGAATATGCTGATTATGATTTATGGGAATTTTTAAAAATTCGTAGTGTTGGTG
S2_c.295G>T         AAATTAATAGAAACATTTAAAGACAAAGAAAATGTTTATTTATTATATGAATATGCTGATTATGATTTATGGGAATTTTTAAAAATTCGTAGTGTTGGTG
S3_c.295G>T         AAATTAATAGAAACATTTAAAGACAAAGAAAATGTTTATTTATTATATGAATATGCTGATTATGATTTATGGGAATTTTTAAAAATTCGTAGTGTTGGTG
S4_c.252A>T         AAATTAATAGAAACATTTAAAGACAAAGAAAATGTTTATTTATTATATGAATATGCTGATTATGATTTATGGGAATTTTTAAAAATTCGTAGTGTTGGTG
S5_c.134A>G         AAATTAATAGAAACATTTAAAGACAAAGAAAATGTTTATTTATTATATGAATATGCTGATTATGATTTATGGGAATTTTTAAAAATTCGTAGTGTTGGTG
S6_c.252A>T         AAATTAATAGAAACATTTAAAGACAAAGAAAATGTTTATTTATTATATGAATATGCTGATTATGATTTATGGGAATTTTTAAAAATTCGTAGTGTTGGTG
S6_c.491A>G         AAATTAATAGAAACATTTAAAGACAAAGAAAATGTTTATTTATTATATGAATATGCTGATTATGATTTATGGGAATTTTTAAAAATTCGTAGTGTTGGTG

                    401                                                                                                500
PF3D7_1121900       TTAATGAAAAAATTACATTTAATATAATACTTCAAATGGTACATGCATTAATATATATACATAACAAAAATATTATTCACAGAGACTTGAAATGTGAAAA
S1_c.152T>G         TTAATGAAAAAATTACATTTAATATAATACTTCAAATGGTACATGCATTAATATATATACATAACAAAAATATTATTCACAGAGACTTGAAATGTGAAAA
S2_c.295G>T         TTAATGAAAAAATTACATTTAATATAATACTTCAAATGGTACATGCATTAATATATATACATAACAAAAATATTATTCACAGAGACTTGAAATGTGAAAA
S3_c.295G>T         TTAATGAAAAAATTACATTTAATATAATACTTCAAATGGTACATGCATTAATATATATACATAACAAAAATATTATTCACAGAGACTTGAAATGTGAAAA
S4_c.252A>T         TTAATGAAAAAATTACATTTAATATAATACTTCAAATGGTACATGCATTAATATATATACATAACAAAAATATTATTCACAGAGACTTGAAATGTGAAAA
S5_c.134A>G         TTAATGAAAAAATTACATTTAATATAATACTTCAAATGGTACATGCATTAATATATATACATAACAAAAATATTATTCACAGAGACTTGAAATGTGAAAA
S6_c.252A>T         TTAATGAAAAAATTACATTTAATATAATACTTCAAATGGTACATGCATTAATATATATACATAACAAAAATATTATTCACAGAGACTTGAAATGTGAAAA
S6_c.491A>G         TTAATGAAAAAATTACATTTAATATAATACTTCAAATGGTACATGCATTAATATATATACATAACAAAAATATTATTCACAGAGACTTGAGATGTGAAAA

                    501                                                                                                600
PF3D7_1121900       TTTTCTTATAAATAAAGATGGAACCATAAAAATGATTGATTTTGGTAGTTCGAAAGATTTAGATAACATAAGCATCAAAACAAATAATGAGGAAGATACA
S1_c.152T>G         TTTTCTTATAAATAAAGATGGAACCATAAAAATGATTGATTTTGGTAGTTCGAAAGATTTAGATAACATAAGCATCAAAACAAATAATGAGGAAGATACA
S2_c.295G>T         TTTTCTTATAAATAAAGATGGAACCATAAAAATGATTGATTTTGGTAGTTCGAAAGATTTAGATAACATAAGCATCAAAACAAATAATGAGGAAGATACA
S3_c.295G>T         TTTTCTTATAAATAAAGATGGAACCATAAAAATGATTGATTTTGGTAGTTCGAAAGATTTAGATAACATAAGCATCAAAACAAATAATGAGGAAGATACA
S4_c.252A>T         TTTTCTTATAAATAAAGATGGAACCATAAAAATGATTGATTTTGGTAGTTCGAAAGATTTAGATAACATAAGCATCAAAACAAATAATGAGGAAGATACA
S5_c.134A>G         TTTTCTTATAAATAAAGATGGAACCATAAAAATGATTGATTTTGGTAGTTCGAAAGATTTAGATAACATAAGCATCAAAACAAATAATGAGGAAGATACA
S6_c.252A>T         TTTTCTTATAAATAAAGATGGAACCATAAAAATGATTGATTTTGGTAGTTCGAAAGATTTAGATAACATAAGCATCAAAACAAATAATGAGGAAGATACA
S6_c.491A>G         TTTTCTTATAAATAAAGATGGAACCATAAAAATGATTGATTTTGGTAGTTCGAAAGATTTAGATAACATAAGCATCAAAACAAATAATGAGGAAGATACA

                    601                                                                                                700
PF3D7_1121900       ATAAATCATGAAGAATTAAGTAAATTTGTATTAAAAAAAAAATAATAACAATAATTCAAACGAAGATTTAAAAAATGCAAATGAATTTAAAAATAATGATA
S1_c.152T>G         ATAAATCATGAAGAATTAAGTAAATTTGTATTAAAAAAAAAATAATAACAATAATTCAAACGAAGATTTAAAAAATGCAAATGAATTTAAAAATAATGATA
S2_c.295G>T         ATAAATCATGAAGAATTAAGTAAATTTGTATTAAAAAAAAAATAATAACAATAATTCAAACGAAGATTTAAAAAATGCAAATGAATTTAAAAATAATGATA
S3_c.295G>T         ATAAATCATGAAGAATTAAGTAAATTTGTATTAAAAAAAAAATAATAACAATAATTCAAACGAAGATTTAAAAAATGCAAATGAATTTAAAAATAATGATA
S4_c.252A>T         ATAAATCATGAAGAATTAAGTAAATTTGTATTAAAAAAAAAATAATAACAATAATTCAAACGAAGATTTAAAAAATGCAAATGAATTTAAAAATAATGATA
S5_c.134A>G         ATAAATCATGAAGAATTAAGTAAATTTGTATTAAAAAAAAAATAATAACAATAATTCAAACGAAGATTTAAAAAATGCAAATGAATTTAAAAATAATGATA
S6_c.252A>T         ATAAATCATGAAGAATTAAGTAAATTTGTATTAAAAAAAAAATAATAACAATAATTCAAACGAAGATTTAAAAAATGCAAATGAATTTAAAAATAATGATA
S6_c.491A>G         ATAAATCATGAAGAATTAAGTAAATTTGTATTAAAAAAAAAATAATAACAATAATTCAAACGAAGATTTAAAAAATGCAAATGAATTTAAAAATAATGATA

                    701                                                                                                800
PF3D7_1121900       GTTTAAATGGCGAAGAAATTAATAATAATGATTTAAATAATACTAATTTTCAAAAAAGTGATAAAAAATATAAAAGATCAAAATTCAAACAAATGCGTTCT
S1_c.152T>G         GTTTAAATGGCGAAGAAATTAATAATAATGATTTAAATAATACTAATTTTCAAAAAAGTGATAAAAAATATAAAAGATCAAAATTCAAACAAATGCGTTCT
S2_c.295G>T         GTTTAAATGGCGAAGAAATTAATAATAATGATTTAAATAATACTAATTTTCAAAAAAGTGATAAAAAATATAAAAGATCAAAATTCAAACAAATGCGTTCT
S3_c.295G>T         GTTTAAATGGCGAAGAAATTAATAATAATGATTTAAATAATACTAATTTTCAAAAAAGTGATAAAAAATATAAAAGATCAAAATTCAAACAAATGCGTTCT
S4_c.252A>T         GTTTAAATGGCGAAGAAATTAATAATAATGATTTAAATAATACTAATTTTCAAAAAAGTGATAAAAAATATAAAAGATCAAAATTCAAACAAATGCGTTCT
S5_c.134A>G         GTTTAAATGGCGAAGAAATTAATAATAATGATTTAAATAATACTAATTTTCAAAAAAGTGATAAAAAATATAAAAGATCAAAATTCAAACAAATGCGTTCT
S6_c.252A>T         GTTTAAATGGCGAAGAAATTAATAATAATGATTTAAATAATACTAATTTTCAAAAAAGTGATAAAAAATATAAAAGATCAAAATTCAAACAAATGCGTTCT
S6_c.491A>G         GTTTAAATGGCGAAGAAATTAATAATAATGATTTAAATAATACTAATTTTCAAAAAAGTGATAAAAAATATAAAAGATCAAAATTCAAACAAATGCGTTCT

                    801                                                                                                900
PF3D7_1121900       AGAAACACTTAAGAATAATGAAAGCTATGAGTTTAATAATCAAATTTTGAGTTCTTTCAAAAGTAACGATTATGCAGCAGATGCAAATAAACATAATAGC
S1_c.152T>G         AGAAACACTTAAGAATAATGAAAGCTATGAGTTTAATAATCAAATTTTGAGTTCTTTCAAAAGTAACGATTATGCAGCAGATGCAAATAAACATAATAGC
S2_c.295G>T         AGAAACACTTAAGAATAATGAAAGCTATGAGTTTAATAATCAAATTTTGAGTTCTTTCAAAAGTAACGATTATGCAGCAGATGCAAATAAACATAATAGC
S3_c.295G>T         AGAAACACTTAAGAATAATGAAAGCTATGAGTTTAATAATCAAATTTTGAGTTCTTTCAAAAGTAACGATTATGCAGCAGATGCAAATAAACATAATAGC
S4_c.252A>T         AGAAACACTTAAGAATAATGAAAGCTATGAGTTTAATAATCAAATTTTGAGTTCTTTCAAAAGTAACGATTATGCAGCAGATGCAAATAAACATAATAGC
S5_c.134A>G         AGAAACACTTAAGAATAATGAAAGCTATGAGTTTAATAATCAAATTTTGAGTTCTTTCAAAAGTAACGATTATGCAGCAGATGCAAATAAACATAATAGC
S6_c.252A>T         AGAAACACTTAAGAATAATGAAAGCTATGAGTTTAATAATCAAATTTTGAGTTCTTTCAAAAGTAACGATTATGCAGCAGATGCAAATAAACATAATAGC
S6_c.491A>G         AGAAACACTTAAGAATAATGAAAGCTATGAGTTTAATAATCAAATTTTGAGTTCTTTCAAAAGTAACGATTATGCAGCAGATGCAAATAAACATAATAGC
```

```
                  901                                                                                                 1000
PF3D7_1121900     TATAAAAAAAAAAAAAACATTCGAAAACTATGTTGGTTCTCCTAATTTCATTCCCCCTGAAGCATTAATAAATAAATGCAGTGGTAAGGCACGAGATTTTT
S1_c.152T>G       TATAAAAAAAAAAAAAACATTCGAAAACTATGTTGGTTCTCCTAATTTCATTCCCCCTGAAGCATTAATAAATAAATGCAGTGGTAAGGCACGAGATTTTT
S2_c.295G>T       TATAAAAAAAAAAAAAACATTCGAAAACTATGTTGGTTCTCCTAATTTCATTCCCCCTGAAGCATTAATAAATAAATGCAGTGGTAAGGCACGAGATTTTT
S3_c.295G>T       TATAAAAAAAAAAAAAACATTCGAAAACTATGTTGGTTCTCCTAATTTCATTCCCCCTGAAGCATTAATAAATAAATGCAGTGGTAAGGCACGAGATTTTT
S4_c.252A>T       TATAAAAAAAAAAAAAACATTCGAAAACTATGTTGGTTCTCCTAATTTCATTCCCCCTGAAGCATTAATAAATAAATGCAGTGGTAAGGCACGAGATTTTT
S5_c.134A>G       TATAAAAAAAAAAAAAACATTCGAAAACTATGTTGGTTCTCCTAATTTCATTCCCCCTGAAGCATTAATAAATAAATGCAGTGGTAAGGCACGAGATTTTT
S6_c.252A>T       TATAAAAAAAAAAAAAACATTCGAAAACTATGTTGGTTCTCCTAATTTCATTCCCCCTGAAGCATTAATAAATAAATGCAGTGGTAAGGCACGAGATTTTT
S6_c.491A>G       TATAAAAAAAAAAAAAACATTCGAAAACTATGTTGGTTCTCCTAATTTCATTCCCCCTGAAGCATTAATAAATAAATGCAGTGGTAAGGCACGAGATTTTT

                  1001                                                                                                1100
PF3D7_1121900     GGAGTCTTGGGTGTACTATTTACCAACTAGTGACATGTACCGTTCCATTTGATGGTTCAACAGAATGGTTCATCTATAACAAAATAAAAAGAAGGGAATT
S1_c.152T>G       GGAGTCTTGGGTGTACTATTTACCAACTAGTGACATGTACCGTTCCATTTGATGGTTCAACAGAATGGTTCATCTATAACAAAATAAAAAGAAGGGAATT
S2_c.295G>T       GGAGTCTTGGGTGTACTATTTACCAACTAGTGACATGTACCGTTCCATTTGATGGTTCAACAGAATGGTTCATCTATAACAAAATAAAAAGAAGGGAATT
S3_c.295G>T       GGAGTCTTGGGTGTACTATTTACCAACTAGTGACATGTACCGTTCCATTTGATGGTTCAACAGAATGGTTCATCTATAACAAAATAAAAAGAAGGGAATT
S4_c.252A>T       GGAGTCTTGGGTGTACTATTTACCAACTAGTGACATGTACCGTTCCATTTGATGGTTCAACAGAATGGTTCATCTATAACAAAATAAAAAGAAGGGAATT
S5_c.134A>G       GGAGTCTTGGGTGTACTATTTACCAACTAGTGACATGTACCGTTCCATTTGATGGTTCAACAGAATGGTTCATCTATAACAAAATAAAAAGAAGGGAATT
S6_c.252A>T       GGAGTCTTGGGTGTACTATTTACCAACTAGTGACATGTACCGTTCCATTTGATGGTTCAACAGAATGGTTCATCTATAACAAAATAAAAAGAAGGGAATT
S6_c.491A>G       GGAGTCTTGGGTGTACTATTTACCAACTAGTGACATGTACCGTTCCATTTGATGGTTCAACAGAATGGTTCATCTATAACAAAATAAAAAGAAGGGAATT

                  1101                                                                                                1200
PF3D7_1121900     AAAATATCCATCCATAATACCTTCTGAATTAATAGACTTAATAGAAAAGTTAACAACTATGAATCCAGAAGAAAGATTAGGGTTTAACGGAGGATGTGAG
S1_c.152T>G       AAAATATCCATCCATAATACCTTCTGAATTAATAGACTTAATAGAAAAGTTAACAACTATGAATCCAGAAGAAAGATTAGGGTTTAACGGAGGATGTGAG
S2_c.295G>T       AAAATATCCATCCATAATACCTTCTGAATTAATAGACTTAATAGAAAAGTTAACAACTATGAATCCAGAAGAAAGATTAGGGTTTAACGGAGGATGTGAG
S3_c.295G>T       AAAATATCCATCCATAATACCTTCTGAATTAATAGACTTAATAGAAAAGTTAACAACTATGAATCCAGAAGAAAGATTAGGGTTTAACGGAGGATGTGAG
S4_c.252A>T       AAAATATCCATCCATAATACCTTCTGAATTAATAGACTTAATAGAAAAGTTAACAACTATGAATCCAGAAGAAAGATTAGGGTTTAACGGAGGATGTGAG
S5_c.134A>G       AAAATATCCATCCATAATACCTTCTGAATTAATAGACTTAATAGAAAAGTTAACAACTATGAATCCAGAAGAAAGATTAGGGTTTAACGGAGGATGTGAG
S6_c.252A>T       AAAATATCCATCCATAATACCTTCTGAATTAATAGACTTAATAGAAAAGTTAACAACTATGAATCCAGAAGAAAGATTAGGGTTTAACGGAGGATGTGAG
S6_c.491A>G       AAAATATCCATCCATAATACCTTCTGAATTAATAGACTTAATAGAAAAGTTAACAACTATGAATCCAGAAGAAAGATTAGGGTTTAACGGAGGATGTGAG

                  1201                                                                                                1300
PF3D7_1121900     GAAATCTTGGAACATTTATATTTTCAAAAATATAATTATAATAAACTTAATTTCATCTTACCAGAAGTTTCCGAGTTAGAAAAACTGTATACAACTATTA
S1_c.152T>G       GAAATCTTGGAACATTTATATTTTCAAAAATATAATTATAATAAACTTAATTTCATCTTACCAGAAGTTTCCGAGTTAGAAAAACTGTATACAACTATTA
S2_c.295G>T       GAAATCTTGGAACATTTATATTTTCAAAAATATAATTATAATAAACTTAATTTCATCTTACCAGAAGTTTCCGAGTTAGAAAAACTGTATACAACTATTA
S3_c.295G>T       GAAATCTTGGAACATTTATATTTTCAAAAATATAATTATAATAAACTTAATTTCATCTTACCAGAAGTTTCCGAGTTAGAAAAACTGTATACAACTATTA
S4_c.252A>T       GAAATCTTGGAACATTTATATTTTCAAAAATATAATTATAATAAACTTAATTTCATCTTACCAGAAGTTTCCGAGTTAGAAAAACTGTATACAACTATTA
S5_c.134A>G       GAAATCTTGGAACATTTATATTTTCAAAAATATAATTATAATAAACTTAATTTCATCTTACCAGAAGTTTCCGAGTTAGAAAAACTGTATACAACTATTA
S6_c.252A>T       GAAATCTTGGAACATTTATATTTTCAAAAATATAATTATAATAAACTTAATTTCATCTTACCAGAAGTTTCCGAGTTAGAAAAACTGTATACAACTATTA
S6_c.491A>G       GAAATCTTGGAACATTTATATTTTCAAAAATATAATTATAATAAACTTAATTTCATCTTACCAGAAGTTTCCGAGTTAGAAAAACTGTATACAACTATTA

                  1301                                                                                                1400
PF3D7_1121900     TAAATAAGTATCATATATATATCAATGAAAAGAGGAAATTAAGACAAAATAATAATTCAACTGAAGAAAATATTAATAATGTAGAAGTACTGAAAAAAAA
S1_c.152T>G       TAAATAAGTATCATATATATATCAATGAAAAGAGGAAATTAAGACAAAATAATAATTCAACTGAAGAAAATATTAATAATGTAGAAGTACTGAAAAAAAA
S2_c.295G>T       TAAATAAGTATCATATATATATCAATGAAAAGAGGAAATTAAGACAAAATAATAATTCAACTGAAGAAAATATTAATAATGTAGAAGTACTGAAAAAAAA
S3_c.295G>T       TAAATAAGTATCATATATATATCAATGAAAAGAGGAAATTAAGACAAAATAATAATTCAACTGAAGAAAATATTAATAATGTAGAAGTACTGAAAAAAAA
S4_c.252A>T       TAAATAAGTATCATATATATATCAATGAAAAGAGGAAATTAAGACAAAATAATAATTCAACTGAAGAAAATATTAATAATGTAGAAGTACTGAAAAAAAA
S5_c.134A>G       TAAATAAGTATCATATATATATCAATGAAAAGAGGAAATTAAGACAAAATAATAATTCAACTGAAGAAAATATTAATAATGTAGAAGTACTGAAAAAAAA
S6_c.252A>T       TAAATAAGTATCATATATATATCAATGAAAAGAGGAAATTAAGACAAAATAATAATTCAACTGAAGAAAATATTAATAATGTAGAAGTACTGAAAAAAAA
S6_c.491A>G       TAAATAAGTATCATATATATATCAATGAAAAGAGGAAATTAAGACAAAATAATAATTCAACTGAAGAAAATATTAATAATGTAGAAGTACTGAAAAAAAA

                  1401                                                                                                1500
PF3D7_1121900     TTTGTTAAACTTAATTAATTCAGATACATTAGTTTGTGCTGAGGAGGAATACGAATCCATAACTTTAAAGAAGAAAATTTTTAAAAGTATTAATTTTATG
S1_c.152T>G       TTTGTTAAACTTAATTAATTCAGATACATTAGTTTGTGCTGAGGAGGAATACGAATCCATAACTTTAAAGAAGAAAATTTTTAAAAGTATTAATTTTATG
S2_c.295G>T       TTTGTTAAACTTAATTAATTCAGATACATTAGTTTGTGCTGAGGAGGAATACGAATCCATAACTTTAAAGAAGAAAATTTTTAAAAGTATTAATTTTATG
S3_c.295G>T       TTTGTTAAACTTAATTAATTCAGATACATTAGTTTGTGCTGAGGAGGAATACGAATCCATAACTTTAAAGAAGAAAATTTTTAAAAGTATTAATTTTATG
S4_c.252A>T       TTTGTTAAACTTAATTAATTCAGATACATTAGTTTGTGCTGAGGAGGAATACGAATCCATAACTTTAAAGAAGAAAATTTTTAAAAGTATTAATTTTATG
S5_c.134A>G       TTTGTTAAACTTAATTAATTCAGATACATTAGTTTGTGCTGAGGAGGAATACGAATCCATAACTTTAAAGAAGAAAATTTTTAAAAGTATTAATTTTATG
S6_c.252A>T       TTTGTTAAACTTAATTAATTCAGATACATTAGTTTGTGCTGAGGAGGAATACGAATCCATAACTTTAAAGAAGAAAATTTTTAAAAGTATTAATTTTATG
S6_c.491A>G       TTTGTTAAACTTAATTAATTCAGATACATTAGTTTGTGCTGAGGAGGAATACGAATCCATAACTTTAAAGAAGAAAATTTTTAAAAGTATTAATTTTATG

                  1501                                                                          1578
PF3D7_1121900     CTTGAAGAATTTGATAAACAGGAAATAAAAGAAATGGAAGAAGCAAGTAAATGGTTAGAACGATATCAAGGTACATAG
S1_c.152T>G       CTTGAAGAATTTGATAAACAGGAAATAAAAGAAATGGAAGAAGCAAGTAAATGGTTAGAACGATATCAAGGTACATAG
S2_c.295G>T       CTTGAAGAATTTGATAAACAGGAAATAAAAGAAATGGAAGAAGCAAGTAAATGGTTAGAACGATATCAAGGTACATAG
S3_c.295G>T       CTTGAAGAATTTGATAAACAGGAAATAAAAGAAATGGAAGAAGCAAGTAAATGGTTAGAACGATATCAAGGTACATAG
S4_c.252A>T       CTTGAAGAATTTGATAAACAGGAAATAAAAGAAATGGAAGAAGCAAGTAAATGGTTAGAACGATATCAAGGTACATAG
S5_c.134A>G       CTTGAAGAATTTGATAAACAGGAAATAAAAGAAATGGAAGAAGCAAGTAAATGGTTAGAACGATATCAAGGTACATAG
S6_c.252A>T       CTTGAAGAATTTGATAAACAGGAAATAAAAGAAATGGAAGAAGCAAGTAAATGGTTAGAACGATATCAAGGTACATAG
S6_c.491A>G       CTTGAAGAATTTGATAAACAGGAAATAAAAGAAATGGAAGAAGCAAGTAAATGGTTAGAACGATATCAAGGTACATAG
```
