## Supplementary Information for "The 3-phosphoinositide-dependent protein kinase 1 is an essential upstream activator of protein kinase A in malaria parasites"

The Supplementary Information file includes:

- Supplementary Figures 1-14
- Supplementary Tables 1 and 2
- Supplementary Data 1

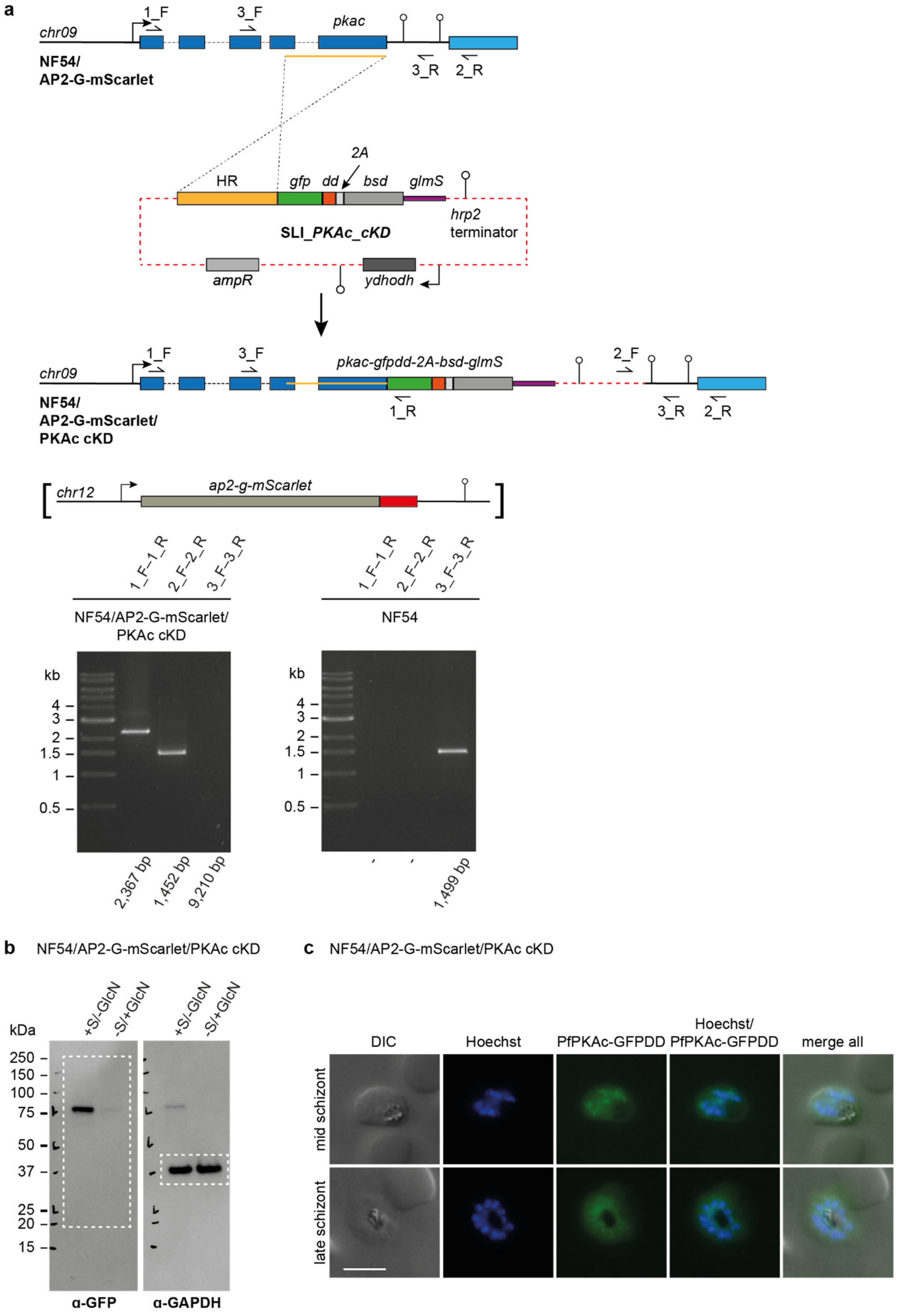


**Supplementary Figure 1 SLI-based engineering and characterisation of the NF54/AP2-G-mScarlet/PKAc cKD parasite line. a** Top: Scheme depicting the wild type *pfpkac* locus, the

SLI_PKAc_cKD construct transfected into NF54/AP2-G-mScarlet parasites and the edited *pfpkac* locus in NF54/AP2-G-mScarlet/PKAc cKD parasites. Primers used for diagnostic PCRs are indicated. Middle: The schematic map of the edited *pfap2-g-mScarlet* locus in the NF54/AP2-G-mScarlet parasite line (Brancucci et al., manuscript in preparation) is shown in brackets. Bottom: Results of PCR reactions performed on gDNA of NF54/AP2-G-mScarlet/PKAc cKD and NF54 WT control parasites confirm correct gene editing. **b** Full size Western blot showing expression of PfPKAc-GFPDD in late schizonts cultured under protein- and RNA-depleting (–Shield-1/+GlcN) and control conditions (+Shield-1/–GlcN). Lysates derived from an equal number of parasites were loaded per lane. The membrane was first probed with α-GFP followed by α-GAPDH control antibodies. MW PfPKAc-GFP = 67.3 kDa, MW PfGAPDH = 36.6 kDa. Dashed lines mark the blot sections shown in Fig. 1a. **c** Expression of PfPKAc-GFPDD in mid and late schizonts under protein- and RNA-stabilizing conditions (+Shield-1/–GlcN) as assessed by live cell fluorescence imaging. Parasites were previously synchronized to an 8-hour window and imaged at 32-40 hpi and 40-48 hpi. Representative fluorescence images are shown. Parasite DNA was stained with Hoechst. DIC, differential interference contrast. Scale bar = 5 μm.

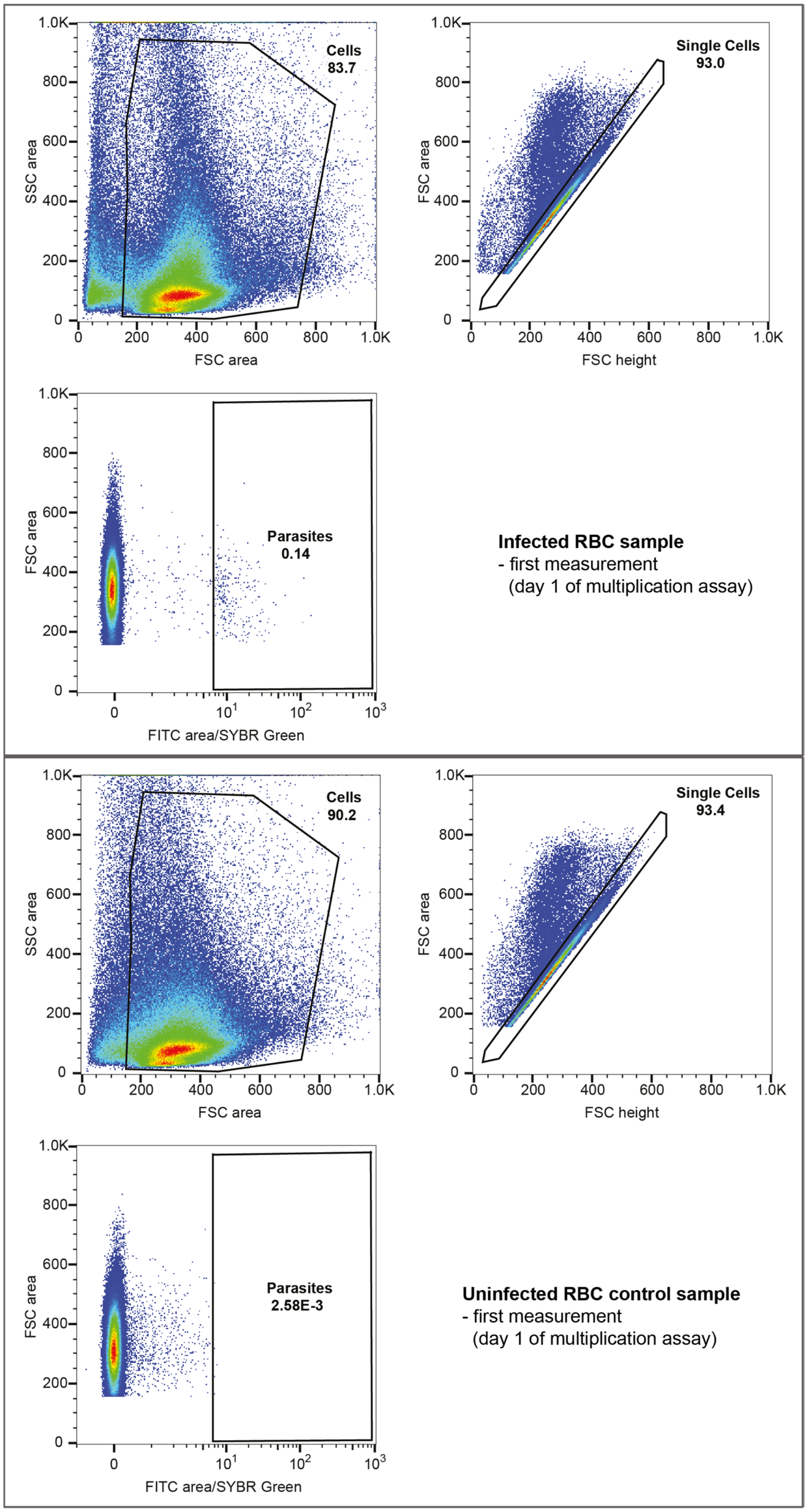


**Supplementary Figure 2 Gating strategy of flow cytometry data obtained from parasite multiplication assays.** Representative flow cytometry plots of a parasite culture (top frame; NF54/AP2-

G-mScarlet/PKAc cKD, +Shield-1/–GlcN) and an uninfected RBC control sample (bottom frame) on day one of the multiplication assay are shown. Events were consecutively gated for the expected cell size, singlets and infected (SYBR green positive) RBCs. The resulting parasite multiplication plots are shown in Figs. 1, 2, 4-6, and in Supplementary Figs. 6, 9, 11.

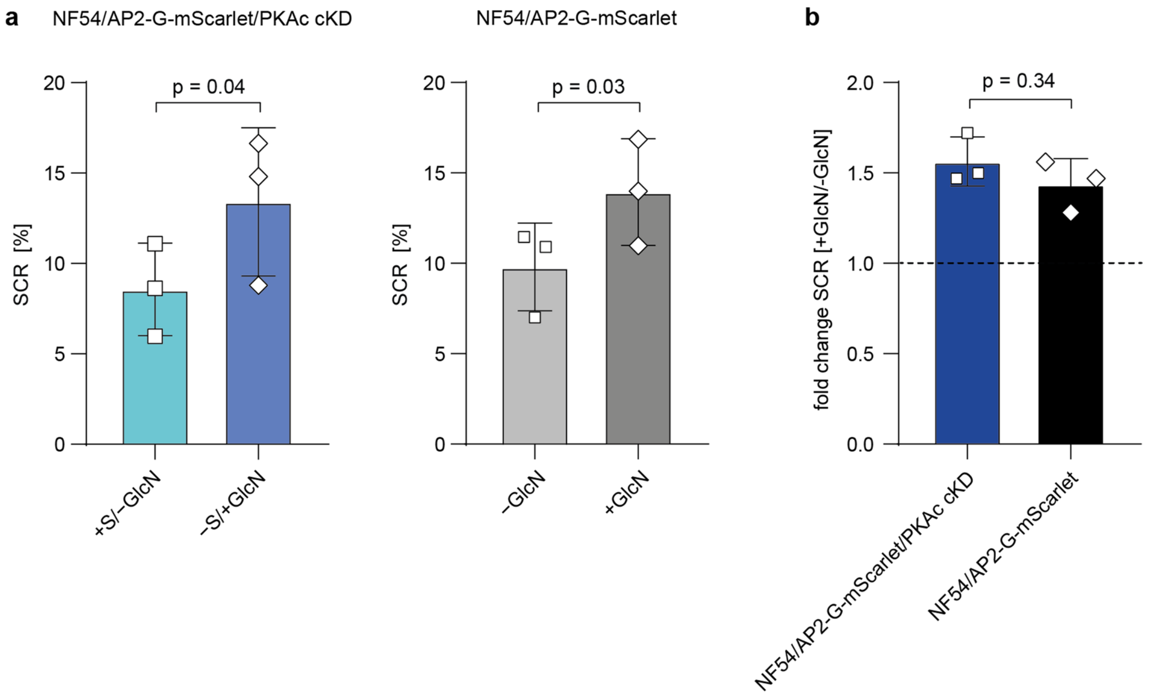


**Supplementary Figure 3 Sexual commitment rates of NF54/AP2-G-mScarlet/PKAc cKD and NF54/AP2-G-mScarlet control parasites.** **a** Left panel: Sexual commitment rates (SCRs) of NF54/AP2-G-mScarlet/PKAc cKD parasites cultured under protein- and RNA-depleting (–Shield-1/+GlcN) (turquoise) and control conditions (+Shield-1/–GlcN) (blue). Right panel: SCRs of NF54/AP2-G-mScarlet control parasites cultured in the presence (+GlcN) (light grey) or absence of GlcN (–GlcN) (dark grey). SCRs were determined by high content imaging and automated image analysis by assessing PfAP2-G-mScarlet positivity among the total number of Hoechst-stained iRBCs. For each experiment, at least 607 Hoechst-positive cells were assessed for PfAP2-G-mScarlet expression. Open squares represent data points for individual replicates and the means and SD (error bars) of three biological replicate experiments are shown. Differences in SCRs have been compared using a paired two-tailed Student's t test (statistical significance cut-off: $p<0.05$). **b** Mean fold change in SCRs of NF54/AP2-G-mScarlet/PKAc cKD parasites cultured under –Shield-1/+GlcN compared to +Shield-1/–GlcN conditions (dark blue) and of NF54/AP2-G-mScarlet control parasites cultured under +GlcN compared to –GlcN conditions (black). Differences in the fold change in SCRs between NF54/AP2-G-mScarlet/PKAc cKD and NF54/AP2-G-mScarlet control parasites have been compared using an unpaired two-tailed Student's t test (statistical significance cut-off: $p<0.05$). S, Shield-1; GlcN, glucosamine.

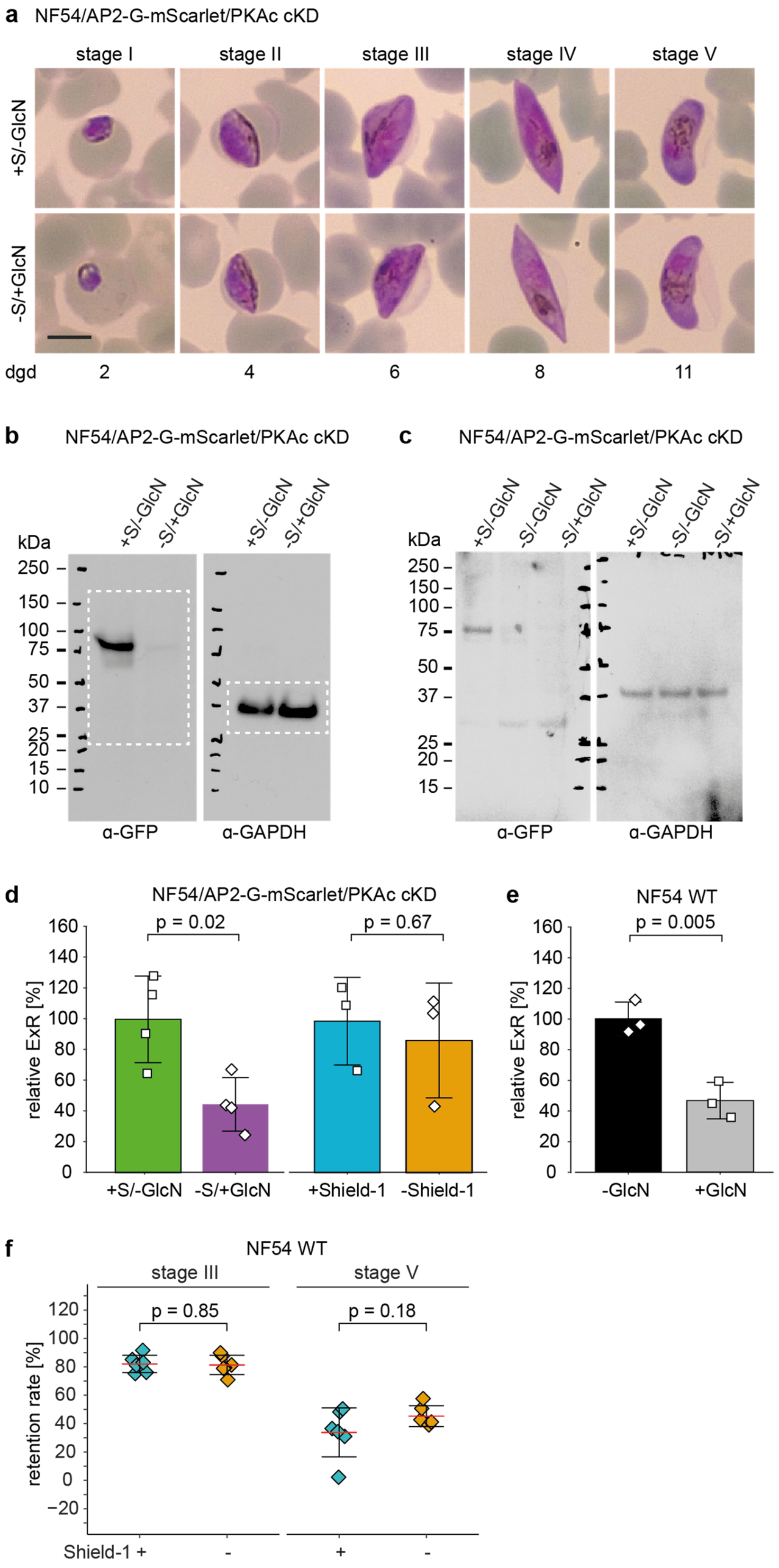


**Supplementary Figure 4 Gametocyte maturation, exflagellation and retention rates of NF54/AP2-G-mScarlet/PKAc cKD parasites. a** Representative images captured from Giemsa-stained blood smears showing the distinct morphology of stage I-V gametocytes cultured under PfPKAc-GFPDD-depleting (–Shield-1/+GlcN) and control conditions (+Shield-1/–GlcN) over eleven days of maturation. Synchronous parasites were split (±Shield-1/±GlcN) as sexual/asexual ring stage parasites 24 hours after the induction of sexual commitment in the preceding IDC. To eliminate asexual parasites, gametocytes were cultured in +SerM supplemented with 50 mM GlcNAc from day one to six of gametocytogenesis. Scale bar = 5 µm. dgd, day of gametocyte development. S, Shield-1; GlcN, glucosamine. **b** Full size Western blot showing expression of PfPKAc-GFPDD in mature stage V gametocytes (day 11) under protein- and RNA-depleting (–Shield-1/+GlcN) and control conditions (+Shield-1/–GlcN). Lysates derived from an equal number of parasites were loaded per lane. The membrane was first probed with α-GFP followed by α-GAPDH control antibodies. MW PfPKAc-GFPDD = 79.8 kDa, MW PfGAPDH = 36.6 kDa. Dashed lines mark the blot sections shown in Fig. 1d. **c** Full size Western blot comparing expression of PfPKAc-GFPDD under protein- and RNA-depleting (–Shield-1/+GlcN), protein-depleting (–Shield-1/–GlcN) and control conditions (+Shield-1/–GlcN) in mature stage V gametocytes (day 11). Lysates derived from an equal number of parasites were loaded per lane. **d** Relative exflagellation rates (ExRs) of NF54/AP2-G-mScarlet/PKAc cKD mature stage V gametocytes (day 14) cultured under protein- and RNA-depleting (–Shield-1/+GlcN) (purple) and control conditions (+Shield-1/–GlcN) (green) or under protein-depleting only (–Shield-1) (orange) and control conditions (+Shield-1) (blue). Open squares represent data points for individual replicates and the means and SD (error bars) of at least three biological replicate experiments are shown. Differences in exflagellation rates have been compared using an unpaired two-tailed Student's t test (statistical significance cut-off: $p<0.05$). **e** Relative exflagellation rates of mature NF54 WT stage V gametocytes (day 14) cultured in presence (+GlcN) (light grey) and absence of GlcN (–GlcN) (black). Parasites were cultured and the results obtained from three biological replicate experiments analysed as described in panel d. **f** Retention rates of NF54 WT stage III (day 6) and stage V (day 11) gametocytes cultured in absence (–Shield-1) (orange) and presence of Shield-1 (+Shield-1) (blue). Coloured squares represent data points for individual replicates and the means and SD (error bars) of one biological replicate experiment with six technical replicates each are shown. Differences in retention rates have been compared using an unpaired two-tailed Student's t test (statistical significance cut-off: $p<0.05$).

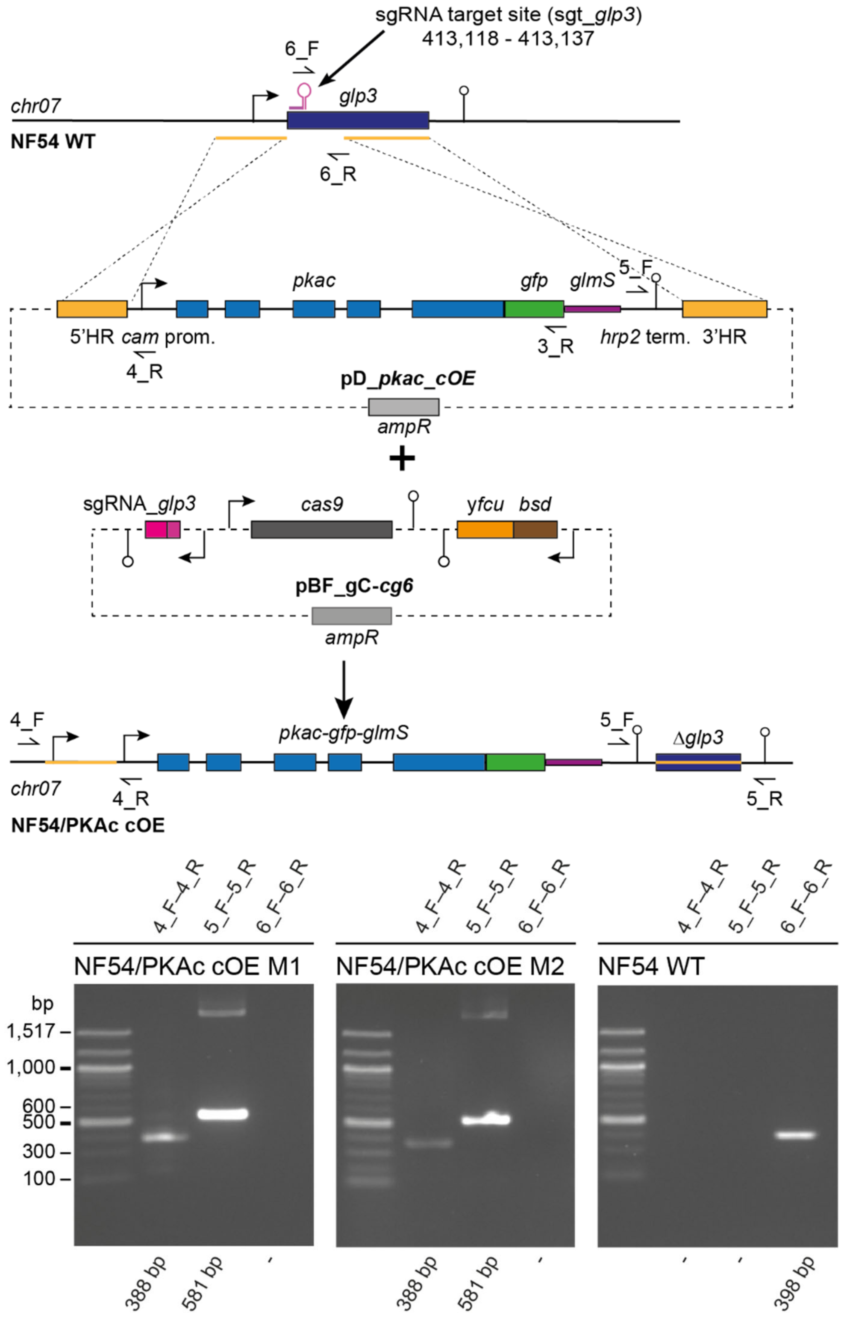


**Supplementary Figure 5 CRISPR/Cas9-based engineering of the NF54/PKAc cOE line.** Top: Scheme depicting the wild type *glp3* target locus, the donor (pD_pkac_cOE) and pBF_gC-cg6 suicide constructs transfected into NF54 WT parasites to generate the NF54/PKAc cOE parasite line and the edited *glp3* locus. Primers used for diagnostic PCRs are indicated. Bottom: Results of PCR reactions performed on gDNA of two clones (M1 and M2) of the NF54/PKAc cOE line and NF54 WT control parasites confirm successful insertion of the PfPKAc cOE cassettes into the *glp3* locus.

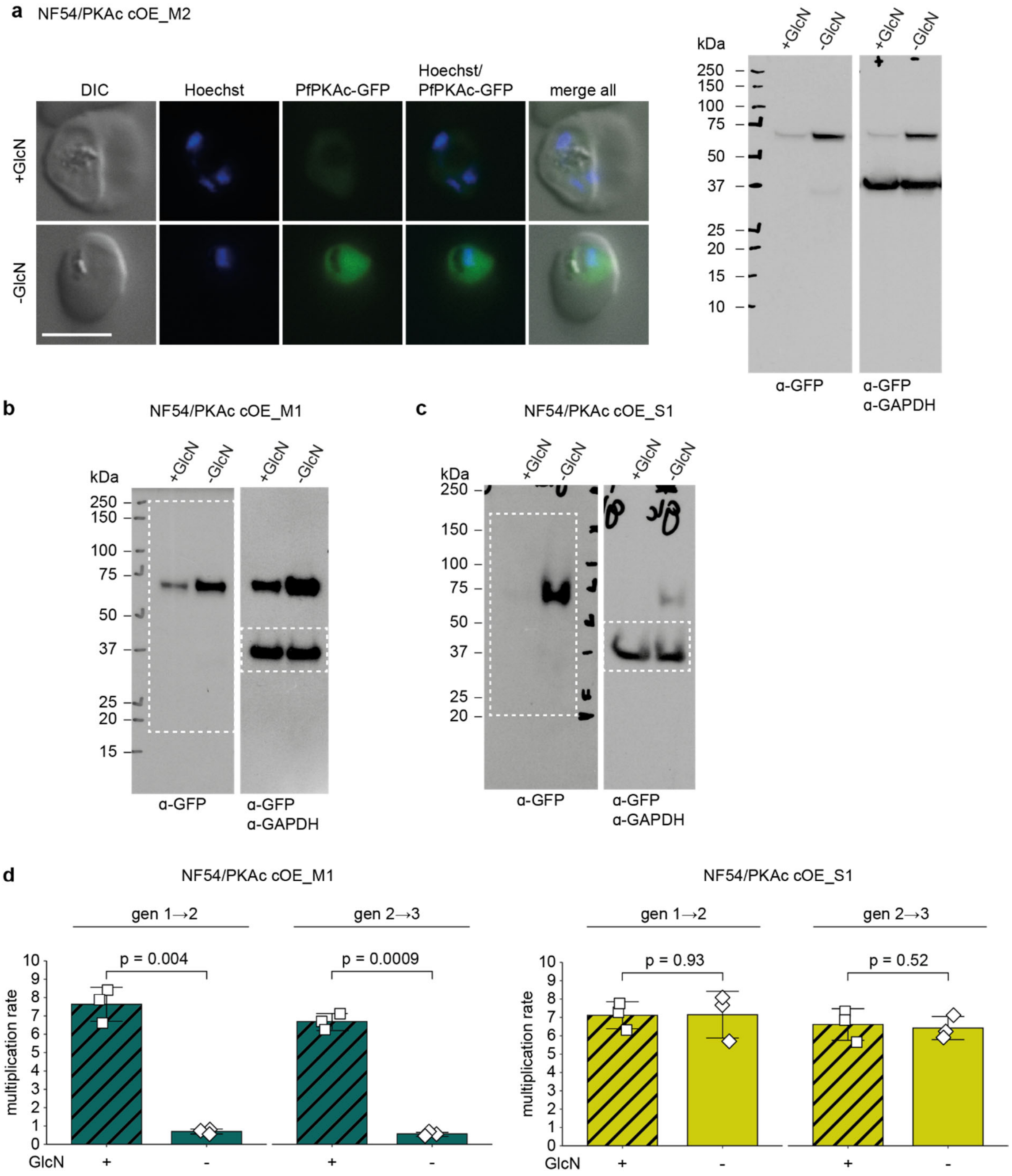


**Supplementary Figure 6 Overexpression of PfPKAc-GFP and multiplication rates of NF54/PKAc cOE M1, M2 and S1 parasites. a** Expression of PfPKAc-GFP in NF54/PKAc cOE M2 parasites under overexpression-inducing (–GlcN) and control conditions (+GlcN) as assessed by live cell fluorescence imaging and Western blot analysis. Synchronous parasites (0-8 hpi) were split (±GlcN) 40 hours before sample collection. Representative fluorescent images are shown. Parasite DNA was stained with Hoechst. DIC, differential interference contrast. Scale bar = 5 µm. For Western blot analysis, parasite lysates derived from equal numbers of parasites were loaded per lane. The membrane was first probed with α-GFP followed by α-GAPDH control antibodies. MW PfPKAc-GFP = 67.3 kDa, MW PfGAPDH = 36.6 kDa. **b** Full size Western blot showing expression of PfPKAc-GFP in NF54/PKAc cOE M1 parasites under overexpression-inducing (–GlcN) and control conditions (+GlcN) conditions. Parasites were cultured and samples prepared as described in panel a. The membrane was first probed with α-GFP followed by α-GAPDH control antibodies. MW PfPKAc-GFPDD = 79.8 kDa, MW PfGAPDH = 36.6 kDa. Dashed lines mark the blot sections shown in Fig. 2a. **c** Full size Western blot shows expression of PfPKAc-GFP in NF54/PKAc cOE S1 parasites under overexpression-inducing (–GlcN) and control (+GlcN) conditions. Parasites were cultured and samples prepared as described in panel a. The membrane was first probed with α-GFP followed by α-GAPDH control antibodies. MW PfPKAc-GFPDD = 79.8 kDa, MW PfGAPDH = 36.6 kDa. Dashed lines mark the blot sections shown in Fig. 2e. MW PfPKAc-GFP = 67.3 kDa, MW PfGAPDH = 36.6 kDa. **d** Parasite multiplication rates of NF54/PKAc cOE M1 (left) and S1 survivor parasites (right) under overexpression-inducing (–GlcN) and control conditions (+GlcN) over two generations. Open squares represent data points for individual replicates and the means and SD (error bars) of three biological replicates are shown. Differences in multiplication rates have been compared using a paired two-tailed Student's t test (statistical significance cut-off: $p<0.05$). Note that the same data is presented as an increase in parasitaemia over time in Figs. 2d and 2g.

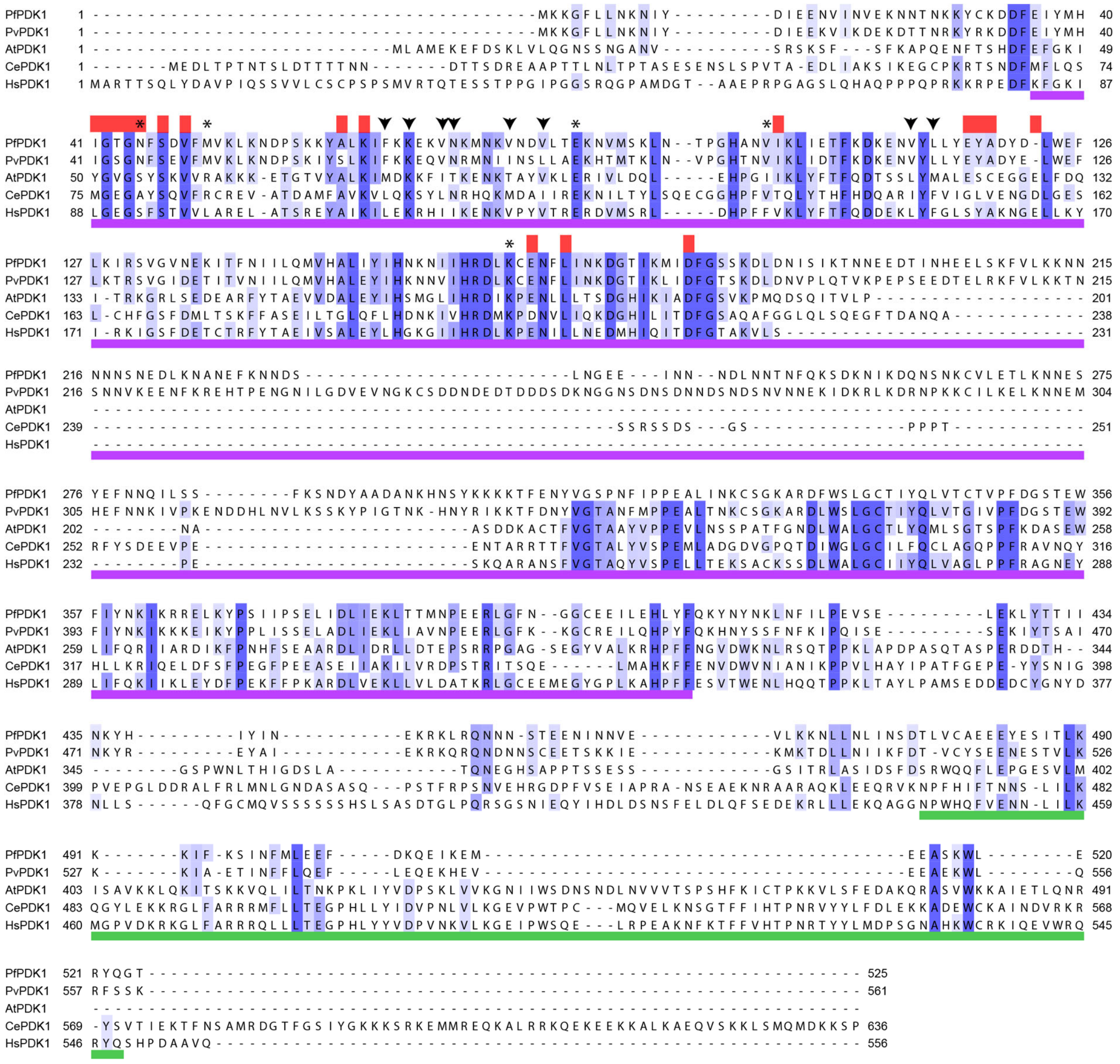

**Supplementary Figure 7 Sequence alignment of PDK1 orthologs.** Clustal Omega [1] multiple sequence alignment of PfPDK1 (Pf3D7_1121900/UniProt ID Q8IIE7) and PvPDK1 (PVX_091715/UniProt ID A5K4N1) with well characterised PDK1 homologues from *Arabidopsis thaliana* (UniProt ID Q9XF67), *Caenorhabditis elegans* (UniProt ID Q9Y1J3) and humans (UniProt ID O15530). Residues are highlighted in blue gradient depending on fractional conservation. The kinase catalytic domain spans residues 82-342 in human PDK1 [2] (purple section underneath the sequences), which also includes a C-terminal pleckstrin-homology (PH) domain spanning residues 446-548 [3] (green section). Arrowheads denote residues forming the PIF-binding pocket in human PDK1 [4], asterisks denote residues mutated in PfPKAc OE survivors identified in this study and red bars denote residues that form part of the ATP-binding cleft.

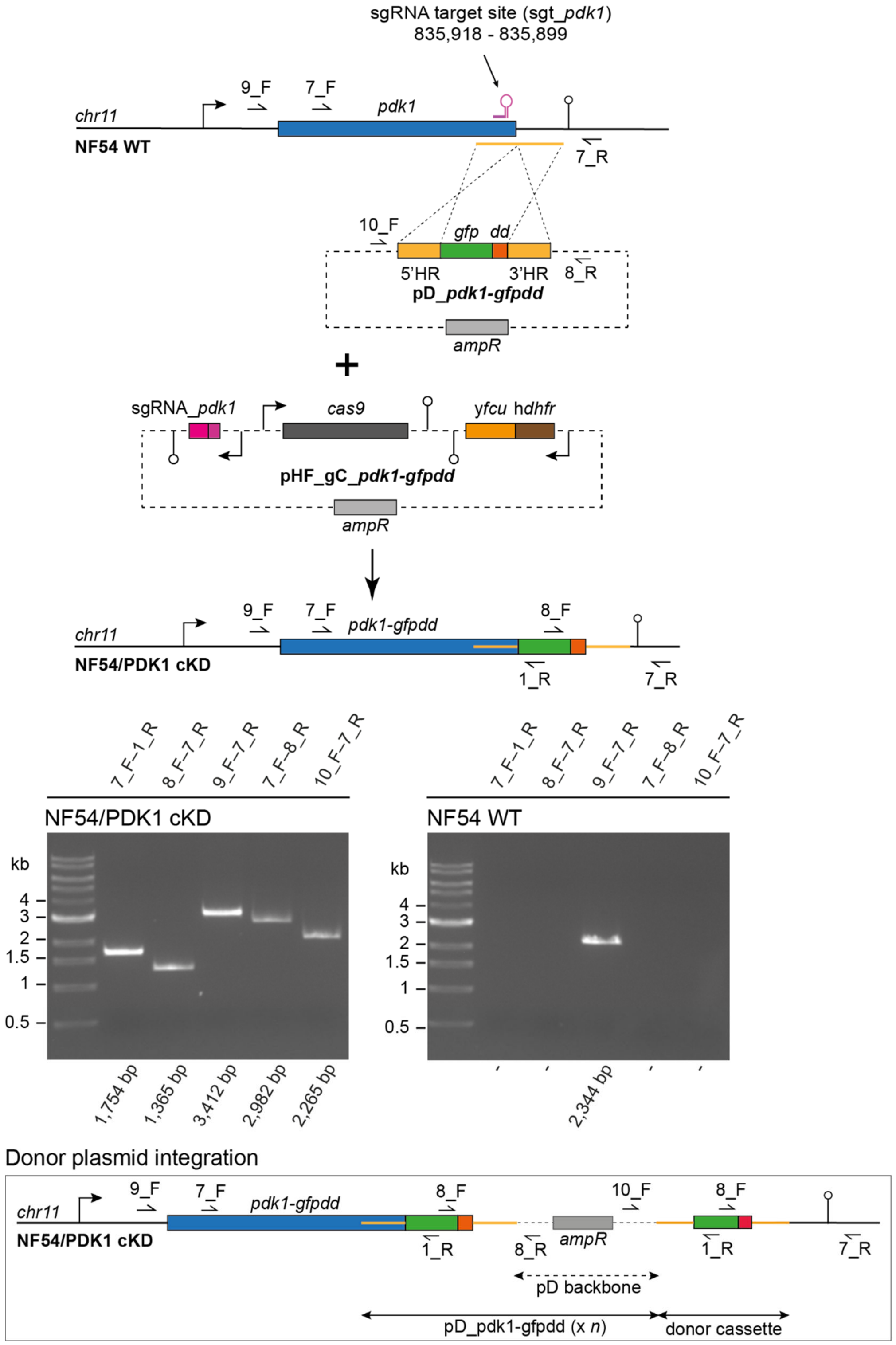


**Supplementary Figure 8 CRISPR/Cas9-based engineering of the NF54/PDK1 cKD parasite line.**

Top: Scheme depicting the wild type *pfpdk1* locus, the donor (pD_pdk1-gfpdd) and the suicide (pHF_gC_pdk1-gfpdd) constructs transfected into NF54 WT parasites to generate the NF54/PDK1 cKD parasite line and the edited *pfpdk1* locus. Primers used for diagnostic PCRs are indicated. Middle:

Results of PCR reactions performed on gDNA of NF54/PDK1 cKD and NF54 WT control parasites confirm correct editing of the *pfpdk1* locus as well as plasmid concatemer integration. Bottom: Schematic map illustrating the integration of a donor plasmid concatemer based on double-crossover recombination of non-adjacent homology regions on the concatemer. To simplify the schematic, the integration of a tandem assembly only is shown.

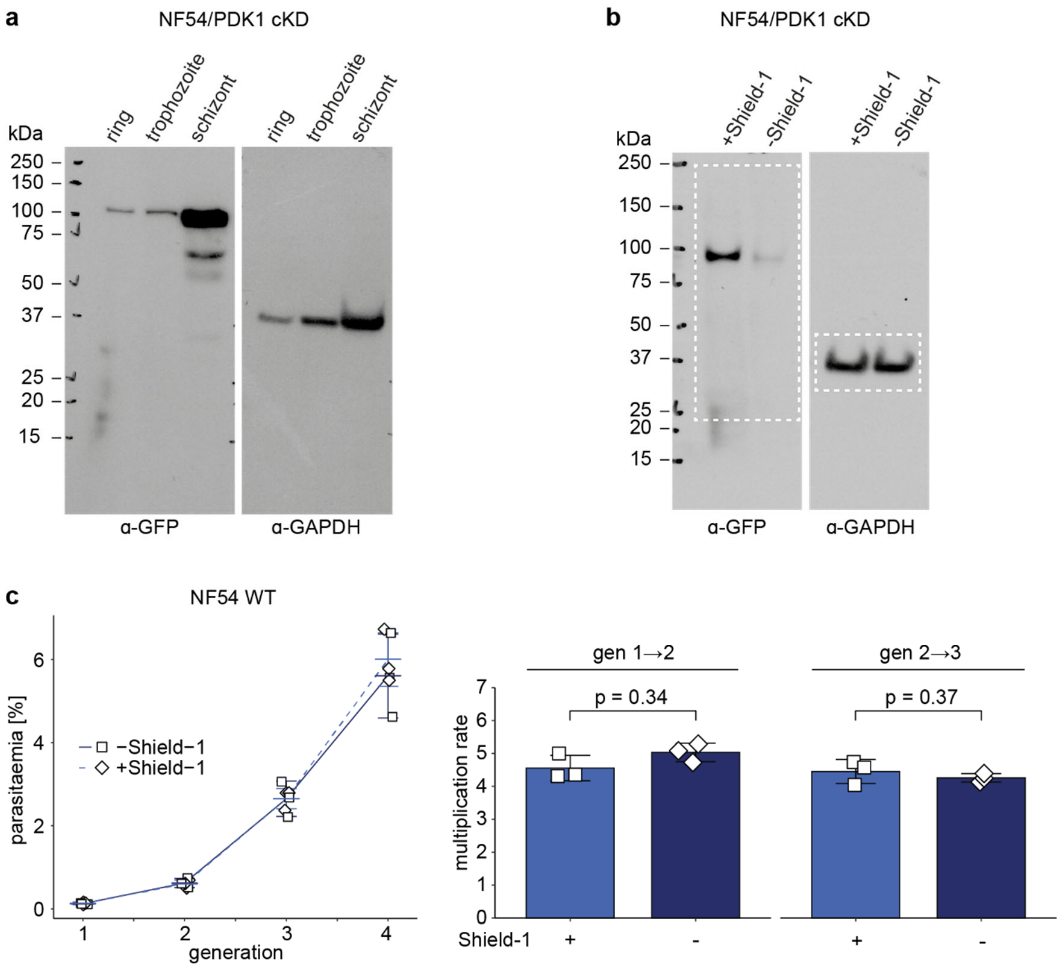


**Supplementary Figure 9 Expression of PfPDK1-GFPDD in NF54/PDK1 cKD parasites and multiplication rates of NF54 WT parasites in presence and absence of Shield-1. a** Expression of PfPDK1-GFPDD in ring (18-24 hpi), trophozoite (24-30 hpi) and schizont (42-48 hpi) stages of NF54/AP2-G-mScarlet/PDK1 cKD parasites under protein-stabilizing (+Shield-1) conditions as assessed by Western blot analysis. Lysates derived from equal numbers of parasites were loaded per lane. The membrane was first probed with α-GFP followed by α-GAPDH control antibodies. MW PfPDK1-GFPDD = 101.1 kDa, MW PfGAPDH = 36.6 kDa. **b** Full size Western blot shows expression of PfPDK1-GFPDD in NF54/AP2-G-mScarlet/PDK1 cKD parasites under protein-depleting (–Shield-1) and control (+Shield-1) conditions. Synchronous parasites (0-8 hpi) were split (±Shield-1) 40 hours before collection of the samples. Lysates derived from equal numbers of parasites were loaded per lane. The membrane was first probed with α-GFP followed by α-GAPDH control antibodies. MW PfPDK1-GFPDD = 101.1 kDa, MW PfGAPDH = 36.6 kDa. Dashed lines mark the blot sections shown in Fig.

4b. **c** Increase in parasitaemia (left) and corresponding parasite multiplication rates (right) of NF54 WT parasites cultured in presence (+Shield-1) and absence of Shield-1 (–Shield-1). Synchronous parasites (0-6 hpi) were split (±Shield-1) 18 hours before the first measurement in generation 1. Open squares represent data points for individual replicates and the means and SD (error bars) of three biological replicates are shown. Differences in multiplication rates have been compared using a paired two-tailed Student's t test (statistical significance cut-off: $p<0.05$).

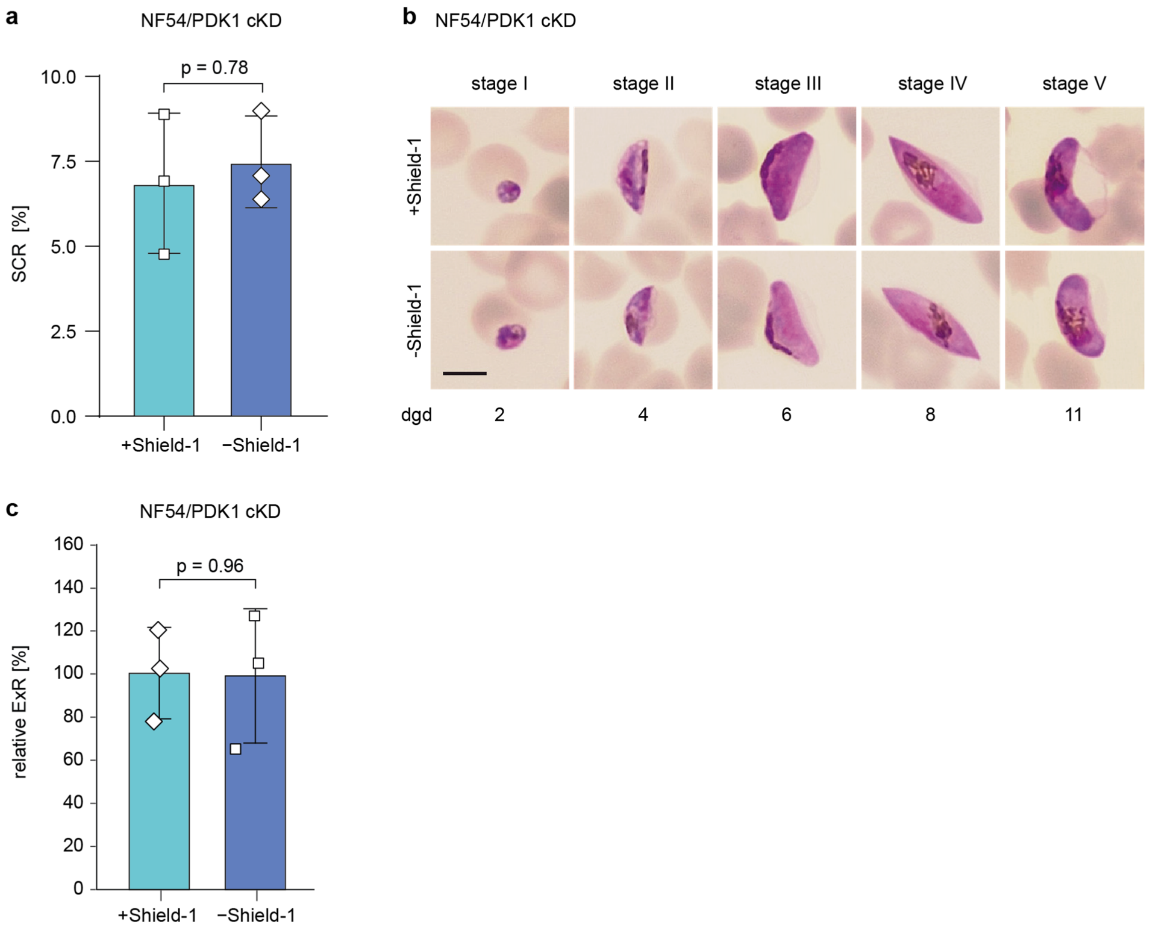


**Supplementary Figure 10 Sexual commitment rates, gametocytogenesis and male gametogenesis of NF54/PDK1 cKD parasites. a** Sexual commitment rates (SCRs) of NF54/PDK1 cKD parasites cultured in the presence (+Shield-1) or absence of Shield-1 (–Shield-1). Open squares represent data points for individual replicates and the means and SD (error bars) of three biological replicate experiments are shown. Differences in SCRs have been compared using a paired two-tailed Student's t test (statistical significance cut-off: $p<0.05$). **b** Representative images captured from Giemsa-stained thin blood smears showing the distinct morphology of stage I-V gametocytes cultured under PfPDK1-GFPDD-depleting (–Shield-1) and control (+Shield-1) conditions over eleven days of maturation. Synchronous parasites were split (±Shield-1) as sexual/asexual ring stage parasites 24 hours after the induction of sexual commitment in the preceding IDC. To eliminate asexual parasites, gametocytes were cultured in +SerM supplemented with 50 mM GlcNAc from day one to six of gametocytogenesis. Scale bar = 5 µm. dgd, day of gametocyte development. **c** Relative exflagellation rates (ExRs) of mature NF54/PDK1 cKD stage V gametocytes (day 14) cultured in presence (+Shield-1) and absence of Shield-

1 (–Shield-1). Synchronous parasites were split (±Shield-1) and cultured as described in panel b. Open squares represent data points for individual replicates and the means and SD (error bars) of three biological replicate experiments are shown. Differences in exflagellation rates have been compared using an unpaired two-tailed Student's t test (statistical significance cut-off: $p<0.05$).

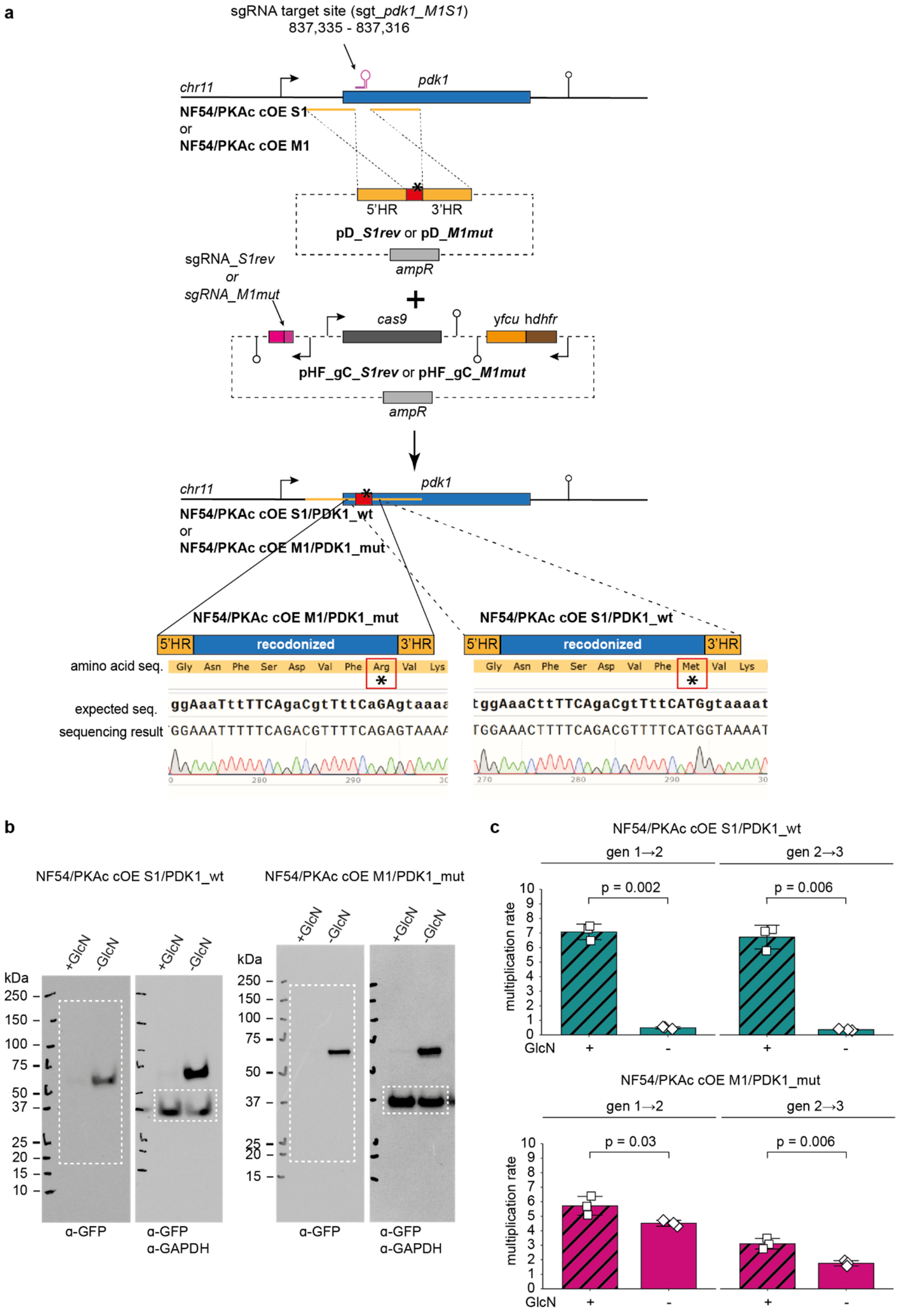


**Supplementary Figure 11 CRISPR/Cas9-based engineering and characterisation of the NF54/PKAc cOE M1/PDK1_mut and NF54/PKAc cOE S1/PDK1_wt lines. a** Top: Scheme depicting the *pfpdk1* locus of NF54/PKAc cOE S1 or M1 parasites, the donor (pD_S1rev or pD_M1mut) and the suicide (pHF_gC_S1rev or pHF_gC_M1mut) constructs transfected into either NF54/PKAc cOE S1 or NF54/PKAc cOE M1 parasites to generate the NF54/ PKAc cOE S1/PDK1_wt and NF54/PKAc cOE M1/PDK1_mut parasite line, respectively, and the edited *pfpdk1* locus. Bottom: Sanger sequencing results of modified *pfpdk1* genes after targeted mutagenesis in NF54/PKAc cOE S1/PDK1_wt and NF54/PKAc cOE M1/PDK1_mut parasites confirms correct editing. The expected sequences after successful editing, the corresponding amino acid changes and sequencing chromatograms are indicated. The asterisk marks the mutated residues (M51R or R51M). Capital letters highlight the synonymous nucleotide substitutions introduced by CRISPR/Cas9 editing to destroy the sgRNA target site and to introduce the aspired amino acid change. **b** Full size Western blots showing expression of PfPKAc-GFP in NF54/PKAc cOE S1/PDK1_wt (left) and NF54/PKAc cOE M1/PDK1_mut (right) parasites under OE-inducing (–GlcN) and control conditions (+GlcN). Synchronous parasites (0-8 hpi) were split (±GlcN) 40 hours before sample collection. Lysates derived from equal numbers of parasites were loaded per lane. The membranes were first probed with α-GFP followed by α-GAPDH control antibodies. MW PfPKAc-GFP = 67.3 kDa, MW PfGAPDH = 36.6 kDa. Dashed lines mark the blot sections shown in Figs. 5a and 5b. **c** Parasite multiplication rates of NF54/PKAc cOE S1/PDK1_wt (top) and NF54/PKAc cOE M1/PDK1_mut (bottom) parasites under OE-inducing (–GlcN) and control conditions (+GlcN) over two generations. Open squares represent data points for individual replicates and the means and SD (error bars) of three biological replicates are shown. Differences in multiplication rates have been compared using a paired two-tailed Student's t test (statistical significance cut-off: $p<0.05$). Note that the same data is presented as an increase in parasitaemia over time in Figs. 5c and 5d.

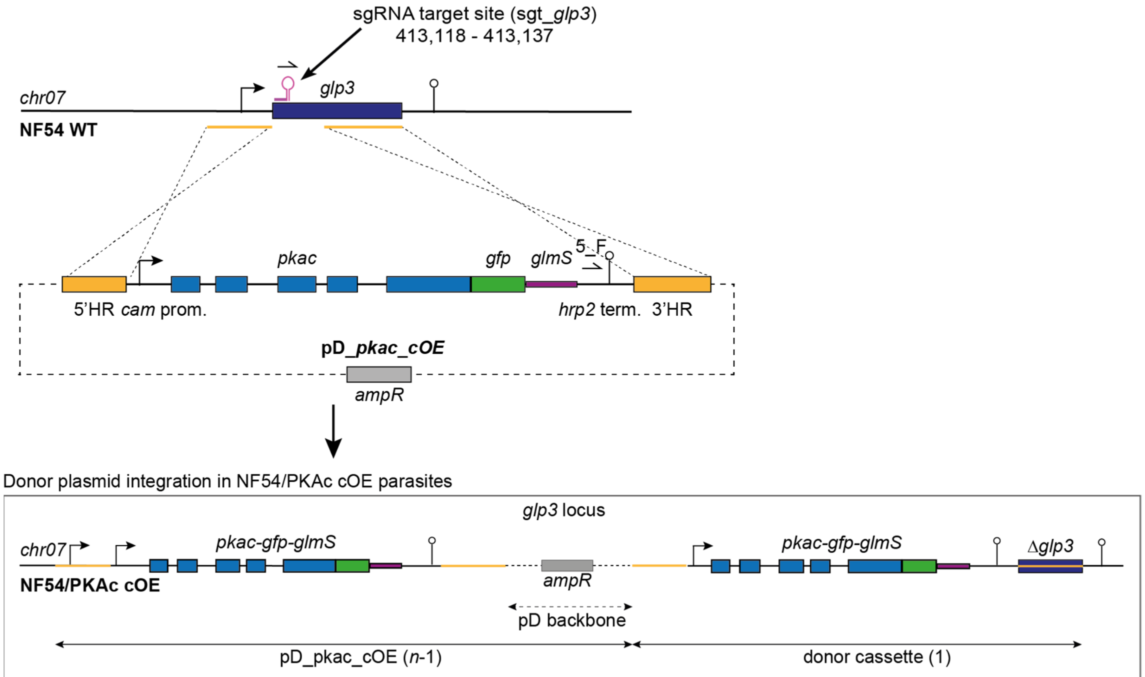


| Parasite name | PfPDK1 mutation | mean copy number PfPKAc cOE cassettes (*n*) (WGS data) |
|---|---|---|
| NF54/PKAc cOE M1 | none | 7.8 |
| NF54/PKAc cOE M2 | none | 7.2 |
| NF54/PKAc cOE S1 | M51R | 2 |
| NF54/PKAc cOE S2 | V99F | 6.3 |
| NF54/PKAc cOE S3 | V99F | 2.6 |
| NF54/PKAc cOE S4 | E84D | 6.1 |
| NF54/PKAc cOE S5 | N45S | 1 |
| NF54/PKAc cOE S6 | E84D/K164R | 5.9 |

**Supplementary Figure 12 Integration of multiple PfPKAc cOE cassettes into the *glp3* locus in NF54/PKAc cOE M1, M2 and S1-S6 parasites.** Top: Scheme depicting the wild type *glp3* target locus and the pD_pkac_cOE donor plasmid used to integrate a PfPKAc cOE cassette into the *glp3* locus in NF54 WT parasites to generate the NF54/PKAc cOE parasite line (see also Supplementary Fig. 5). Bottom: The boxed schematic illustrates the integration of pD_pkac_cOE donor plasmid concatemers into the *glp3* locus based on double-crossover recombination of non-adjacent homology regions on the concatemer. For reasons of simplicity, the integration of a tandem assembly only is shown. *n*-1, number of integrated donor plasmids. Estimated mean copy numbers of integrated PfPKAc cOE cassettes (*n*) are shown for the two unselected NF54/PKAc cOE clones (M1, M2) and the six independently grown survivor populations (S1-S6), alongside the PfPDK1 mutations identified in the six NF54/PKAc cOE survivors (see also Fig. 3). Copy numbers of PfPKAc cOE cassettes were calculated from WGS data and the analysis steps are described in the Methods section.

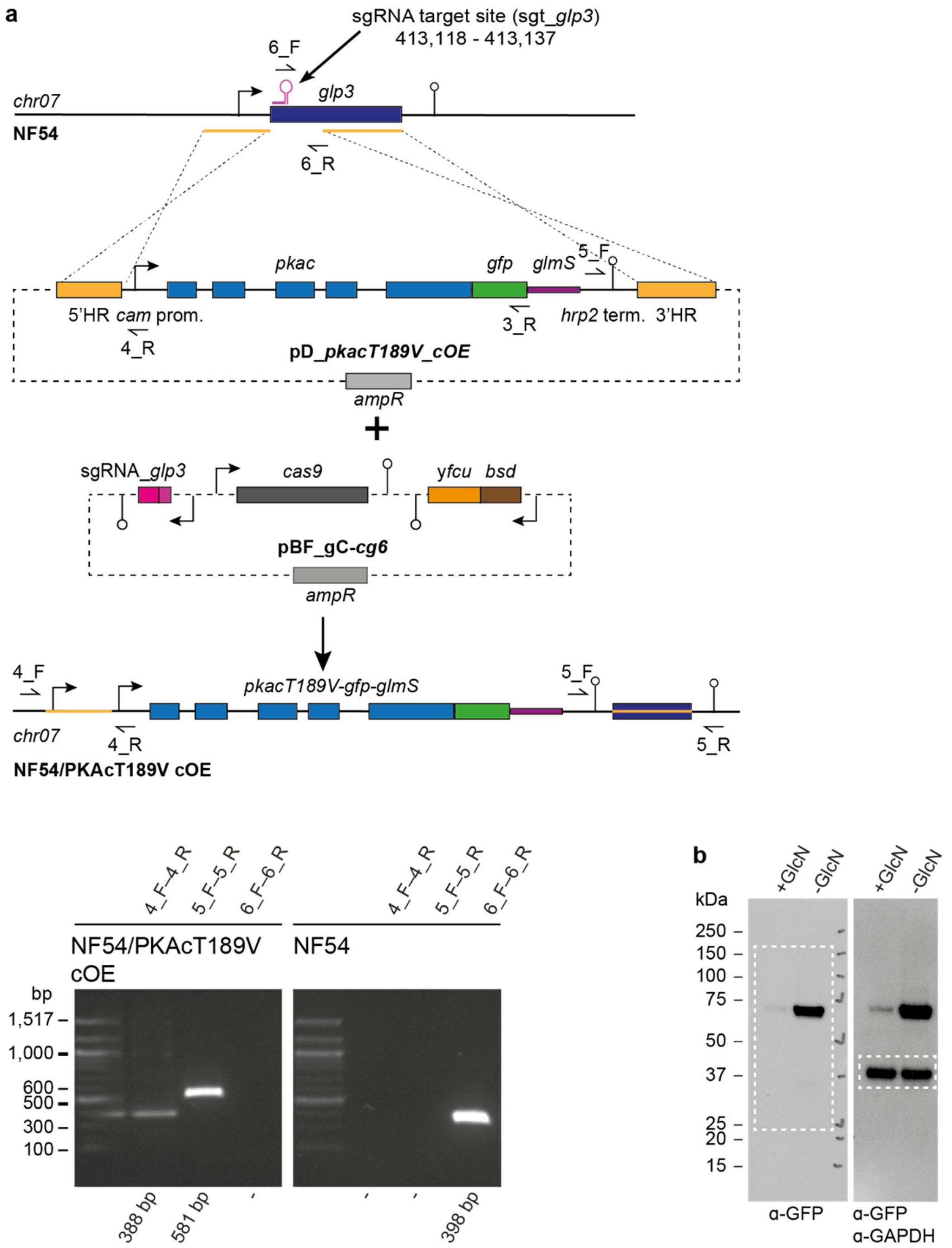


**Supplementary Figure 13 CRISPR/Cas9-based engineering and characterisation of the NF54/PKAcT189V cOE line. a** Top: Scheme depicting the wild type *glp3* target locus, the donor (pD_pkacT189V_cOE) and pBF_gC-cg6 suicide constructs transfected into NF54 WT parasites to generate the NF54/PKAcT189V cOE parasite line and the edited *glp3* locus. Primers used for diagnostic PCRs are indicated. Bottom: Results of PCR reactions performed on gDNA of the NF54/PKAcT189V cOE line and NF54 WT control parasites confirm successful insertion of the PfPKAcT189V cOE cassette into the *glp3* locus. **b** Full size Western blot showing expression of PfPKAcT189V-GFP in NF54/PKAcT189V cOE parasites under OE-inducing (–GlcN) and control conditions (+GlcN). Synchronous parasites (0-8 hpi) were split (±GlcN) 40 hours before collection of the samples. Lysates derived from equal numbers of parasites were loaded per lane. The membrane was first probed with α-GFP followed by α-GAPDH control antibodies. MW PfPKAcT189V-GFP = 67.3 kDa, MW PfGAPDH = 36.6 kDa. Dashed lines mark the blot sections shown in Fig. 6a.

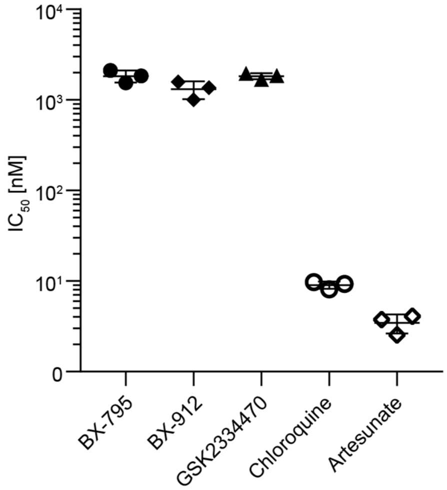


**Supplementary Figure 14 Activity of inhibitors of human PDK1 on *P. falciparum* blood stage parasite multiplication.** $IC_{50}$ values for the three human PDK1 inhibitors BX-795, BX-912 and GSK2334470 and the two antimalarial control compounds Chloroquine and Artesunate on *P. falciparum* asexual blood stage parasite multiplication. Symbols represent individual $IC_{50}$ values calculated from two technical replicate dose response assays each. Drug dose response assays were performed in three biological replicates, means and SD (error bars) are indicated.

**Supplementary Table 1 Oligonucleotides used for cloning of transfection constructs.**

| Oligo-nucleotide name | Oligonucleotide sequence 5' → 3' | Plasmids | Cell lines |
|---|---|---|---|
| PCRA_F | CTGGCGTAATAGCGAAGAGG | SLI_PKAc_cKD; pD_S1rev; pD_M1mut | NF54/AP2-G-mScarlet/PKAc cKD; NF54/ PKAc cOE S1/PDK1_wt; NF54/PKAc cOE M1/PDK1_mut |
| PCRA_R | CATTAATGAATCGGCCAACG | SLI_PKAc_cKD; pD_S1rev; pD_M1mut | NF54/AP2-G-mScarlet/PKAc cKD; NF54/ PKAc cOE S1/PDK1_wt; NF54/PKAc cOE M1/PDK1_mut |
| gfp_F | CGTTGGCCGATTCATTAATGactagtagcataggatccagtggaatgagt | SLI_PKAc_cKD | NF54/AP2-G-mScarlet/PKAc cKD |
| dd_R | CGCTACCACTTTCCAGTTTTAGAagctcc | SLI_PKAc_cKD | NF54/AP2-G-mScarlet/PKAc cKD |
| 2A_F | TCTAAAACTGGAAAGTGGTAGCGgagaaggaaga | SLI_PKAc_cKD | NF54/AP2-G-mScarlet/PKAc cKD |
| bsd_R | GAACAAGATTACTCGAGTTAGCCCtcc | SLI_PKAc_cKD | NF54/AP2-G-mScarlet/PKAc cKD |
| glmS1_F | GGGCTAACTCGAGTAATCTTGTTCttattttctcaatagg | SLI_PKAc_cKD | NF54/AP2-G-mScarlet/PKAc cKD |
| term_R | CCTCTTCGCTATTACGCCAGgtcgacgaattctagatttaataaatatgttctt | SLI_PKAc_cKD | NF54/AP2-G-mScarlet/PKAc cKD |
| ydhodh_F | AGAAATATATCATGACAGCCAGtttaactaccaagt | SLI_PKAc_cKD | NF54/AP2-G-mScarlet/PKAc cKD |
| ydhodh_R | ATATCCTTAATTAATTAAATGCTGTTCaacttcccacg | SLI_PKAc_cKD | NF54/AP2-G-mScarlet/PKAc cKD |
| pbdt3_F | GAACAGCATTTAATTAATTAAGGATATggcagcttaatgttc | SLI_PKAc_cKD | NF54/AP2-G-mScarlet/PKAc cKD |
| pbdt3_R | TCGCTATTACGCCAGgtcgacctaccctgaagaagaaaagtcc | SLI_PKAc_cKD | NF54/AP2-G-mScarlet/PKAc cKD |
| cam1_F | ATTTCTCATTATATATAAGAACATATTTATTAAATgctatccgatcttataaggaaattcc | SLI_PKAc_cKD | NF54/AP2-G-mScarlet/PKAc cKD |
| cam1_R | CTGGCTGTCATGATATATTTCTattaggtatttattattataaaatataaatcttg | SLI_PKAc_cKD | NF54/AP2-G-mScarlet/PKAc cKD |
| pka3'_F | CGTTGGCCGATTCATTAATGagctaaaatagtcgagacgagaac | SLI_PKAc_cKD | NF54/AP2-G-mScarlet/PKAc cKD |
| pka3'_R | TTCTCCTTTACTCATTCCACTGgatccccaatcataaaatggatcattttcatttg | SLI_PKAc_cKD | NF54/AP2-G-mScarlet/PKAc cKD |
| glp3_F | CTTCTTCAGGGTAGCATGaac | pD_pkac_cOE; | NF54/PKAc cOE |
| glp3_R | GGATAGCTACATGTTCATAtttatatttatttattc | pD_pkac_cOE; | NF54/PKAc cOE |
| cam_F | TATGAACATGTAGCTATCCgatcttataaggaaattccc | pD_pkac_cOE; | NF54/PKAc cOE |
| hrp2_R | CATGCTACCCTGAAGAAGgaattctagatttaataaatatgttc | pD_pkac_cOE; | NF54/PKAc cOE |
| glmS_F | GCATGGATGAACTATACAAAtaatcttgttcttattttctcaatag | pD_pkac_cOE; | NF54/PKAc cOE |
| glmS_R | CTCATATACTTCCCTAGATGAGatttttcttcctcctaagattg | pD_pkac_cOE; | NF54/PKAc cOE |
| clon_F | TAATAAATACCTAATAGAAATATATC*actagtagtggatccagtgga* | pD_pkac_cOE; | NF54/PKAc cOE |
| clon_R | GAAAAGTTCTTCTCCTTTACTCAT*tccactggatccactactagt* | pD_pkac_cOE; | NF54/PKAc cOE |
| pka_F | TACCTAATAGAAATATATCACTAGTatgcagtttattaaaaatttgc | pD_pkac_cOE; pD_pkacT189V_cOE | NF54/PKAc cOE |
| pka_R | CTCCTTTACTCATTCCACTGgatccccaatcataaaatggatcat | pD_pkac_cOE; pD_pkacT189V_cOE | NF54/PKAc cOE |
| T189V_F | ATGTTTTATGTGGAACTCCAgaatatatc | pD_pkacT189V_cOE | NF54/PKAcT189V cOE |
| T189V_R | TGGAGTTCCACATAAAACATaagttctcgtctcgac | pD_pkacT189V_cOE | NF54/PKAcT189V cOE |
| hr1_F | CGTTGGCCGATTCATTAATGgtgtaaaagttaaaaaaacatgc | pD_S1rev; pD_M1mut | NF54/PDK1_S1rev; NF54/PKAc cOE M1/PDK1_mut |
| rev_F | CTTTTCAGACGTTTTCATGGtaaaattaaaaaatgatccttcaaaaaaatatg | pD_S1rev | NF54/PKAc cOE S1/PDK1_wt |
| rev_R | CCATGAAAACGTCTGAAAAGtttccagttcctatgtgcatatatatttc | pD_S1rev | NF54/PKAc cOE S1/PDK1_wt |
| mut_F | TTTTTCAGACGTTTTCAGAGtaaaattaaaaaatgatccttcaaaaaaatatg | pD_M1mut | NF54/PKAc cOE M1/PDK1_mut |
| mut_R | CTCTGAAAACGTCTGAAAAAtttccagttcctatgtgcatatatatttc | pD_M1mut | NF54/PKAc cOE M1/PDK1_mut |
| hr2_R | CCTCTTCGCTATTACGCCAGagctattatgtttatttgcatctg | pD_S1rev; pD_M1mut | NF54/PKAc cOE S1/PDK1_wt; NF54/PKAc cOE M1/PDK1_mut |
| hr1KD_F | CGTTGGCCGATTCATTAATGtagtgacatgtaccgttcc | pD_pdk1-gfpdd | NF54/PDK1 cKD |
| hr1KD_R | GTTCCCTGGTATCTCTCAAGccacttacttgcttcttccatttc | pD_pdk1-gfpdd | NF54/PDK1 cKD |
| gfpdd_F | CTTGAGAGATACCAGGGAACtagtggatccagtggaatgagtaaag | pD_pdk1-gfpdd | NF54/PDK1 cKD |
| gfpdd_R | CTATCATTCTAATTTTAGAAGCTCCACac | pD_pdk1-gfpdd | NF54/PDK1 cKD |
| hr2KD_F | TGGAGCTTCTAAAATTAGAATGATAGaatataacatatataaataaaaacaatttcttttac | pD_pdk1-gfpdd | NF54/PDK1 cKD |
| hr2KD_R | CCTCTTCGCTATTACGCCAGgtgtttcacaagaacttaagg | pD_pdk1-gfpdd | NF54/PDK1 cKD |
| sgRNA_S1rev_F | TATT*gaatttcagtgatgtgttta* | pHF_gC_S1rev | NF54/PKAc cOE S1/PDK1_wt |
| sgRNA_S1rev_R | AAAC*taaacacatcactgaaattc* | pHF_gC_S1rev | NF54/PKAc cOE S1/PDK1_wt |
| sgRNA_M1mut_F | TATT*gaatttcagtgatgtgttta* | pHF_gC_M1mut | NF54/PKAc cOE M1/PDK1_mut |
| sgRNA_M1mut_R | AAAC*taaacacatcactgaaattc* | pHF_gC_M1mut | NF54/PKAc cOE M1/PDK1_mut |
| sgRNA_pdk1_F | TATT*aaatggttagaacgatatca* | pHF_gC_pdk1-gfpdd | NF54/PDK1 cKD |
| sgRNA_pdk1_R | AAAC*tgatatcgttctaaccattt* | pHF_gC_pdk1-gfpdd | NF54/PDK1 cKD |

Names and sequences of oligonucleotides, plasmids and cell lines are indicated. Sequences essential for Gibson assembly reactions (Gibson overhangs) or for T4 DNA ligase-dependent cloning of double-stranded sgRNA-encoding fragments (5' and 3' overhangs) are highlighted with capital letters. Italicized letters highlight the annealed sequences (sgRNAs) and colour-highlighted letters represent introduced sequence mutations.

**Supplementary Table 2 Primers used for diagnostic PCRs on gDNA of transgenic parasite lines.**

| Primer name | Sequence 5' → 3' | Cell line |
|---|---|---|
| 1_F | atgcagtttattaaaaatttgc | NF54/AP2-G-mScarlet/PKAc cKD |
| 1_R | gtgtgagttatagttgtattcc | NF54/AP2-G-mScarlet/PKAc cKD |
| 2_F | gcgaggaagcggaagagc | NF54/AP2-G-mScarlet/PKAc cKD |
| 2_R | cattatcaaaacaggcaattg | NF54/AP2-G-mScarlet/PKAc cKD |
| 3_F | ggatcattcaaagatgactc | NF54/AP2-G-mScarlet/PKAc cKD |
| 3_R | gtatgtgaaaacaactaaaacatg | NF54/AP2-G-mScarlet/PKAc cKD |
| 4_F | attatgggaaaataatccttac | NF54/PKAc cOE M1;<br>NF54/PKAc cOE M2<br>NF54/PKAcT189V cOE |
| 4_R | gctcagagattgcatgcaag | NF54/PKAc cOE M1;<br>NF54/PKAc cOE M2<br>NF54/PKAcT189V cOE |
| 5_F | ctttaattttattttggtcatg | NF54/PKAc cOE M1;<br>NF54/PKAc cOE M2<br>NF54/PKAcT189V cOE |
| 5_R | cttttacaatatgaacataaagtac | NF54/PKAc cOE M1;<br>NF54/PKAc cOE M2<br>NF54/PKAcT189V cOE |
| 6_F | gttcatgctcctcaacaaag | NF54/PKAc cOE M1;<br>NF54/PKAc cOE M2<br>NF54/PKAcT189V cOE |
| 6_R | gaacaaatacataagagcgc | NF54/PKAc cOE M1;<br>NF54/PKAc cOE M2<br>NF54/PKAcT189V cOE |
| 7_F | aaaccaggacatgcaaatgttatt | NF54/PDK1 cKD |
| 7_R | tctaatgaattgtccatcatgc | NF54/PDK1 cKD |
| 8_F | ggttatgtacaggaaagaac | NF54/PDK1 cKD |
| 8_R | attcgccattcaggctgc | NF54/PDK1 cKD |
| 9_F | gatcgaaccaagcttatattaac | NF54/PDK1 cKD |
| 10_F | gcgaggaagcggaagagc | NF54/PDK1 cKD |

**Supplementary Data 1 Multiple nucleotide sequence alignment of Pf3D7_1121900/*pfpdk1* sequences of the NF54/PKAc cOE clones M1 and M2 and the six PfPKAc OE-tolerant survivor populations S1-S6 determined by WGS.** The *pfpdk1* coding sequences of the NF54/PKAc cOE M1 and M2 clones are identical to the Pf3D7_1121900 reference sequence retrieved from PlasmoDB (www.plasmodb.org) (top row). *pfpdk1* coding sequences of the NF54/PKAc cOE survivor populations S1-S6 are shown and deviations from the reference sequence are highlighted in green. NF54/PKAc cOE survivor S6 consists of two subpopulations with one carrying the c.252A>T mutation and the other one carrying the c.491A>G mutation (as verified by inspection of the sequencing read pairs).
